# Clinical relevance of diffusion tensor imaging-based parameters estimated with deep learning: the effect of model retraining

**DOI:** 10.64898/2026.11.05.764523

**Authors:** Dominika Ciupek, Antonio Tristán Vega, Maciej Malawski, Tomasz Pieciak

## Abstract

Deep learning (DL)-based approaches have recently become powerful tools for characterizing microstructural properties of brain tissue from diffusion-weighted magnetic resonance imaging data. These techniques enable depicting microstructural features with significantly reduced acquisition schemes while preserving relatively high image-quality metrics compared with non-DL-based approaches. Learning-based methods have increasingly been applied to diffusion tensor imaging (DTI) and more advanced multicompartmental biophysical models, with the potential to facilitate multiparameter analyses of brain microstructure in clinical settings. However, independently trained DL models–while maintaining image-quality properties–do not appear to preserve the statistical properties of groupwise analyses across independently trained models.

In this study, we systematically evaluate the statistical properties of DL-based DTI parameters across independently trained neural networks under the same architecture to preserve group-level effects in bipolar disorder. We examine the impact of model retraining and its dependence on reducing the number of gradient directions compared with the nonlinear least-squares (NLLS) procedure. Our results show that initializing model weights randomly from a uniform distribution, while keeping the network architecture and training hyperparameters the identical in the experiment, leads to significant variation in the statistical properties of DTI measures inferred from independently trained models. The analyses showed that the true positive rate in groupwise analysis using the DL-based DTI parameters ranges from 0% to 100% compared with the NLLS-based analysis. As the number of gradient directions increased, the NLLS procedure yielded more stable groupwise statistical results, whereas independently trained DL models showed greater variability despite similar image-quality agreement with reference maps. The results indicate that the DTI maps retrieved from a single trained network should be treated with great caution before drawing clinical conclusions.

## 1. Introduction

Diffusion-weighted magnetic resonance imaging (DW MRI) is a well-established, non-invasive medical imaging modality that enables to examine tissue microstructure by measuring the diffusive motion of water molecules (Basser et al., 1994; Le Bihan et al., 2001). In biological tissues, such as neural tissue, water diffusion is restricted or hindered by cellular membranes, fibres, and organelles. As water diffusion is highly sensitive to these microscopic environments, DW MRI is an effective tool for characterizing brain alterations in ageing (Westlye et al., 2010; Cox et al., 2016; Pieciak et al., 2023; Kim et al., 2026), the impact of environmental factors (Lewandowska et al., 2025), and various neurodegenerative and neuropsychiatric conditions (Kubicki et al., 2007; Pasternak et al., 2018; Carreira Figueiredo et al., 2022).

To quantitatively characterize tissue properties, one can model the diffusion-weighted MR signal using various signal representations, with diffusion tensor imaging (DTI) being the most widely studied technique (Basser et al., 1994; Westin et al., 2002). DTI provides quantitative features that enable characterization of water diffusion magnitude through mean diffusivity (MD) and the directional variance of diffusion *via* fractional anisotropy (FA). These parameters serve as sensitive indicators of tissue organization, cellular complexity, and the integrity of white matter microstructure (Basser and Jones, 2002; Alexander et al., 2007; Minati and Węglarz, 2007).

Standard parametric DTI estimation approaches (Basser et al., 1994; Koay et al., 2006; Maximov et al., 2011) are based on clinically accepted protocols. While a six-direction protocol is mathematically correct for estimating DT (Basser and Pierpaoli, 1998), the estimation may be sensitive to noise and other confounding factors. Some analytical frameworks have been proposed to regularize the estimation procedure while preserving tissue boundaries (Fillard et al., 2007; Liu et al., 2013). However, as demonstrated by Ni et al. (2006); Lebel et al. (2012), relying on six diffusion-sensitizing gradient directions schemes can lead to unreliable results in voxel-based analyses. As a remedy, at least 20 (and 30) non-collinear sampling orientations were suggested to robustly estimate the FA (and MD) parameter in the brain, thereby minimizing their uncertainty (Jones, 2004; Ni et al., 2006).

Recently, deep learning (DL)-based methods have become increasingly favored for estimating DTI and DTI-based parameters (Golkov et al., 2016; Gibbons et al., 2019; Aliotta et al., 2021; Li et al., 2021; Park et al., 2021; Goodwin-Allcock et al., 2022; Gaviraghi et al., 2022; Aja-Fernández et al., 2023; Gaviraghi et al., 2024; Martin et al., 2025; Ametepe et al., 2026; Chen et al., 2026). Numerous studies have shown that these procedures outperform parametric DTI fitting techniques, particularly when only a few diffusion gradient directions are available. To this end, a wide range of DL architectures have been explored, including multilayer perceptrons (Golkov et al., 2016; Aliotta et al., 2021; Aja-Fernández et al., 2023; Ametepe et al., 2026) and convolutional neural networks (CNNs) (Gibbons et al., 2019; Park et al., 2021; Goodwin-Allcock et al., 2022; Aja-Fernández et al., 2023), with U-Net models (Aliotta et al., 2021; Li et al., 2021; Gaviraghi et al., 2022; Aja-Fernández et al., 2023; Gaviraghi et al., 2024) being the most widely adopted. More recent studies employ simulation-based inference (Eggl and De Santis, 2026) and advanced generative architectures, such as conditional diffusion probabilistic models, which additionally enable estimation of parameter uncertainty (Martin et al., 2025), or Kolmogorov-Arnold networks (Chen et al., 2026).

Despite a wide range of DL-based estimation techniques used for DTI estimation, the performance of quantitative descriptors is typically evaluated from a signal processing perspective using metrics derived from natural image analysis, such as mean squared error (MSE) or mean structural similarity index measure (MSSIM) (Aliotta et al., 2021; Park et al., 2021; Goodwin-Allcock et al., 2022; Martin et al., 2025; Ametepe et al., 2026; Chen et al., 2026). Besides, the evaluations remain largely limited to datasets from healthy individuals (Goodwin-Allcock et al., 2022; Martin et al., 2025; Ametepe et al., 2026; Chen et al., 2026). Studies evaluating the clinical applicability of DL-based DTI features are limited to gliomas (Aliotta et al., 2021), stroke (Gibbons et al., 2019; Li et al., 2021), multiple sclerosis (Golkov et al., 2016; Park et al., 2021; Gaviraghi et al., 2022, 2024), and migraine assessment (Aja-Fernández et al., 2023). Mostly, the models were developed using data from healthy individuals (Li et al., 2021; Gaviraghi et al., 2022; Aja-Fernández et al., 2023; Gaviraghi et al., 2024) or synthetic datasets (Park et al., 2021), with limited testing on clinical data. Only a few studies trained DL models using clinical data (Golkov et al., 2016) or a combination of data from healthy controls and patients with particular conditions (Gibbons et al., 2019; Aliotta et al., 2021). Other works compared the distributions of estimated parameters within specific regions of interest (Gaviraghi et al., 2022, 2024). While these studies provide useful insights, they do not assess whether the estimated maps preserve subtle, disease-related microstructural differences or maintain the statistical properties critical for clinical analyses. Recent analyses have shown that nearly identical conventional image quality metrics can lead to noticeably different outcomes in downstream statistical analyses (Aja-Fernández et al., 2023; Gaviraghi et al., 2024).

Unlike clinical conditions with clearly visible structural abnormalities, such as tumors or large ischemic lesions, neurodegenerative and neuropsychiatric disorders often exhibit subtle brain alterations that can only be detected through grouplevel statistical analyses. Bipolar disorder (BD) is one such example, which presents a significant clinical diagnosis challenge, primarily because it often begins with a depressive episode that can be indistinguishable from unipolar depression (Grande et al., 2016). Although there are no clinically validated biomarkers or objective genetic tests available, DTI findings have indicated significant changes in several major white matter tracts in BD patients, both inter- and intra-hemispheric, such as the corpus callosum (Barysheva et al., 2013; Magioncalda et al., 2018; Lee et al., 2020; Koshiyama et al., 2020), internal capsule (Barysheva et al., 2013; Lee et al., 2020; Koshiyama et al., 2020), and corona radiata (Barysheva et al., 2013; Magioncalda et al., 2018; Lee et al., 2020). These alterations are often observed as reduced FA and positive or negative alterations in MD or radial diffusivity (RD) (Barysheva et al., 2013; Magioncalda et al., 2018; Lee et al., 2020). As the differences are subtle and localized to specific white matter tracts, even small prediction errors introduced by a DL-based approach can significantly affect the results of statistical analyses, despite the model demonstrating high agreement with reference maps using standard image-quality metrics.

Motivated by this limitation, the present study systematically investigates the clinical relevance of DL-based DTI quantitative maps estimated from reduced acquisition schemes. We evaluate DTI parameters estimated with independently trained DL models under the same configuration using voxel-wise tract-based spatial statistics (TBSS) and ROI-based approaches. This study quantifies the extent to which clinically significant group differences are preserved between patients with BD and healthy controls (HC).

The main contributions of this work can be summarized as follows:

1. We experimentally demonstrate that although DTI measures predicted by a DL model trained on experimental data (i.e., individuals with BD) are characterized with stable image-quality metrics (i.e., MSE, MSSIM) across repeated training runs, their statistical properties vary considerably from run to run.
2. We reveal that models trained with the same neural network architecture but different weight initializations can exhibit markedly different statistical properties in groupbased statistical analyses.
3. We suggest that DL-based methods for predicting DTI measures may not be universally applicable when the number of diffusion gradients is high, as the true positive rate (TPR) and false positive rate (FPR) in group-based analyses show greater variability than TPR and FPR obtained from parametric DTI.

Overall, this work highlights the importance of evaluating DL-based parameter estimation using application-specific clinical analyses rather than relying exclusively on global image quality metrics, and the lack of repeatability in groupwise analyses when DTI parameters are inferred from independently trained DL models under the identical underlying architecture.

## 2. Materials and methods

### 2.1. Population study and data

We conducted the experimental study using the publicly available UCLA Consortium for Neuropsychiatric Phenomics MRI dataset (Poldrack et al., 2016). The cohort includes 130 HCs (68M/62F, mean age 31 ±9) and 49 individuals diagnosed with BD (28M/21F, mean age 35 ±9). Participants in the HC group were excluded if they had lifetime diagnoses of schizophrenia or other psychotic disorders, bipolar I or II disorder, or substance abuse or dependence (excluding caffeine or nicotine). They were also excluded if they had current major depressive disorder, suicidality, anxiety disorders, or attention deficit hyperactivity disorder (ADHD). The BD group excluded individuals with any of these other diagnoses (such as schizophrenia and ADHD), although stable medications were allowed for the patients. Participants were also excluded if they were left-handed, believed they might be pregnant, or had other contraindications to scanning (such as claustrophobia, having metal in their body, or a body size too large to fit in the scanner). After screening the dataset, we further excluded one BD subject (ID: sub-60048) due to excessively noisy data.

DW MR data were acquired on a 3T Siemens Trio scanner (Siemens, Erlangen, Germany) using an echo-planar sequence with the following parameters: repetition time (TR) = 9000 ms, echo time (TE) = 93 ms, *b*-value = 1000 s/mm^2^, 64 gradient directions, and one non-diffusion-weighted (*b*-value = 0), image slice thickness = 2 mm, acquisition matrix = 96 ×96, and a flip angle of 90°.

### 2.2. Data preprocessing

The released DW MR data were initially corrected for eddy currents and head motion artifacts using the FSL v6 eddy (Jenkinson et al., 2002). In addition, we further preprocessed the dataset using the following pipeline: 1) denoising in magnitude domain *via* Marčenko-Pastur principal component analysis using the MRtrix3 `dwidenoise` (Veraart et al., 2016a,b), 2) Gibbs ringing artifact correction through the local subvoxelshifts method *via* `mrdegibbs` (Kellner et al., 2016), and 3) normalization of spatial intensity variations using the MRtrix3 `dwibiascorrect` with the fast command as provided in FSL (Zhang et al., 2002).

### 2.3. Experimental configurations

We consider three experimental configurations to generate pseudo-ground-truth DTI metrics and investigate the statistical properties of DL-reconstructed DTI measures. Each configuration represents a separate approach for generating the reduced acquisition schemes and defining the pseudo-reference DTI maps. The configurations differ in how they compute the reference maps for the DL procedure and the subsampling scenario. Specifically, in **Configuration 1**, we first resample the dataset into an incremental 64-direction dataset, then subsample the data for the analysis by selecting the first *k* volumes from the resampled dataset. In **Configuration 1**, the dataset is resampled each time a subsampled analysis is performed. The resampling and subsampling procedures used in each configuration are described in detail in the Supplementary materials (section 1.1). Fig. 1 provides a schematic overview of two configurations used in this work, namely **Configuration 1** and **Configuration 2**, while Fig. S12 in the Supplementary materials illustrates **Configuration 3**.

**Figure 1:**
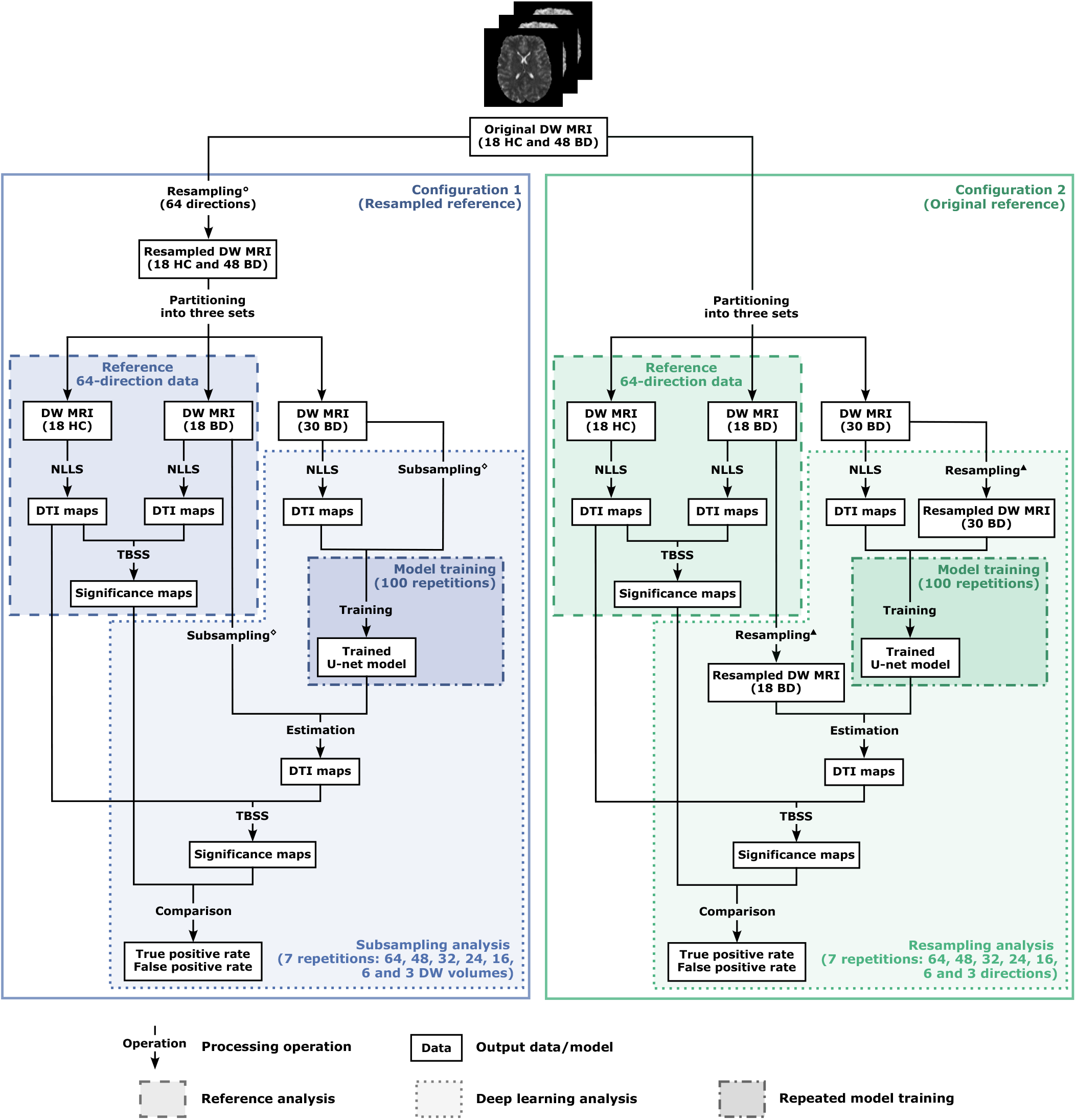
Overview of the two experimental configurations used for deep learning-based diffusion tensor imaging (DTI): **Configuration 1** (Resampled reference) and **Configuration 2** (Original reference). The key difference between these two scenarios lies in the stage at which the data are subsampled. In Configuration 1, the original data are first resampled to an incremental 64-direction gradient scheme. The resulting dataset is then used to generate both the reference DTI maps and to prepare reduced inputs for model training, with the latter generated by selecting the first *k* (*k* = {64, 48, 32, 24, 16, 6, 3}) gradient directions at a single shell. In Configuration 2, the original 64-direction acquisition is used directly to generate the reference DTI maps, while reduced datasets are generated separately by resampling the original dataset to seven subsampled datasets with *k* (*k* = {64, 48, 32, 24, 16, 6, 3}) gradient directions. Each configuration consists of three main stages: reference analysis, deep learning analysis, and repeated model training. The reference analysis is performed once for the 64-direction data in both configurations. At this stage, DTI maps are estimated using the non-linear least squares (NLLS) method, and the resulting statistical significance maps serve as the reference. The deep learning analysis is performed separately for each of the seven input schemes. For each scheme, the model training is repeated 100 times, with a separate initialization weights instance, resulting in 100 independently trained U-Net models. Each trained model is then used to infer deep learning-based DTI maps. From these maps, statistical significance maps are obtained and compared with the corresponding reference maps. The agreement between the deep learning-based and reference significance maps is evaluated using the true positive and false positive rates. The symbols used for resampling and subsampling refer to the corresponding procedures described in the Supplementary materials (see section 1.1 therein). Resampling° denotes *Data resampling (Configuration 1)*, Subsampling^⋄^ denotes *Data subsampling (Configuration 1)*, and Resampling^▲^ denotes *Data resampling (Configuration 2)*.

1. **Configuration 1 (Resampled reference):** The 64direction dataset resampled to the incremental 64-direction dataset serves as the reference for the DL-based DTI. RD and FA maps were computed using the NLLS fitting procedure, as implemented in the DIPY library v1.11.0 (`dipy.reconst.dti`; Garyfallidis et al. (2014)), and then used as the pseudo-ground truth for the neural network training procedure. We prepared the reduced inputs with *k* = {48, 32, 24, 16, 6, 3} gradient directions by selecting the first *k* volumes from the resampled dataset.
2. **Configuration 2 (Original reference):** RD and FA maps computed from the original dataset using the NLLS algorithm serve as the reference for the DL-based DTI. Reduced datasets were generated independently by resampling the original data to new gradient tables with *k* = {48, 32, 24, 16, 6, 3} gradient directions.
3. **Configuration 3 (Rotated gradients):** To assess the sensitivity of DTI parameters estimated with the NLLS algorithm to gradient distribution, we also perform a rotation-based evaluation. Specifically, this analysis focuses exclusively on gradient distribution schemes that include *k* = {64, 48, 32, 24, 16, 6} directions. We exclude the 3-direction scheme because the NLLS algorithm requires at least six diffusion gradient directions to estimate the diffusion tensor. For each gradient setup, we generate 100 rigid rotations. Rotation matrices are sampled from the special orthogonal group SO(3) using a Haar-uniform distribution (SciPy v.1.16.0, `scipy.stats.special_ortho_group`). To ensure reproducibility, we generated all rotations using a fixed random seed, resulting in a deterministic set of 100 rotation matrices shared across all experiments. We then used the rotated gradient tables to recompute DTI maps using the NLLS fitting procedure.

**Configuration 1** serves as the primary experimental setup for all analyses reported in this study. **Configuration 2** and **Configuration 3** are employed as complementary analyses. In particular, Configuration 2 is used to verify that the trends observed in the main experiments are not specific to a particular resampling scheme or pseudo-reference DTI maps. Configuration 3 is introduced to characterize the variability of DTI estimates obtained with the NLLS method under the rotated gradient tables.

### 2.4. Dataset preparation for training procedure

The study cohort was divided into three subsets: 1) 18 HC, 2) 18 patients with BD, and 3) 30 additional individuals with BD. The 18 HC subset matches as closely as possible to the 18 BD group in terms of age and sex, as both groups are subsequently used in the between-group analyses.

- In the RD analysis, the 18 HC and 18 BD groups has the same sex distribution (12 males and 6 females) and a mean age of 38 ± 8 years. In contrast, the 30 BD group included 15 males and 15 females, with a mean age of 34 ± 9 years.
- For the FA analysis, the sex distributions in the HC and BD groups were 10 males and 8 females, with mean ages of 34 ± 10 and 32 ±8 years, respectively. The 30 BD group had 17 male and 13 female patients, with a mean age of 37 ± 9 years.

The reason for using different cohort partitions for the RD and FA analyses is explained in detail in section 4.5.

Reference statistical analysis: The groups of 18 HC and 18 BD participants were used for the reference tract-based spatial statistics (TBSS) (Smith et al., 2006, 2004) (dashed rectangles in Fig. 1) and DL-derived TBSS (dotted rectangles in Fig. 1), as well as for the region of interest (ROI)-based analyses.

Training dataset: The remaining 30 BD subjects serve as the training dataset for the DL models. To prepare the dataset for training, only slices with a significant portion of the brain tissue are selected. In total, we select 40 central axial slices. The DW MRI data are normalized by dividing the diffusion-weighted MR signal at each diffusion gradient direction by the corresponding non-diffusion-weighted slice. The reference RD maps were normalized by the maximum physiologically observed RD value (3.0 × 10^−3^ mm^2^*/*s). We divided each selected slice into four overlapping 2D patches of size 64 × 64 pixels, yielding 160 images per subject and 4800 images in total for the entire training dataset. This approach provides sufficient data to ensure convergence of the training procedure. Finally, we did not use the 18 BD subset for model training. Instead, we used it as an independent evaluation dataset to assess the performance of the trained neural networks.

### 2.5. Deep learning-based DTI maps estimation

The employed DL architecture follows a U-Net scheme (Ronneberger et al., 2015). We trained a separate U-Net model for each acquisition scheme (we call it a “**model instance**”), with 64 input channels for the full scheme and 48, 32, 24, 16, 6, or 3 input channels for the reduced acquisitions. Each input channel represents a diffusion-weighted 2D image acquired along a single gradient direction. The encoder begins with an initial block with 64 features, and the number of features doubles until it reaches 512 at the bottleneck. The decoder mirrors this structure, starting from the bottleneck and producing the output DTI maps. A customized loss function, inspired by Tian et al. (2020); Aliotta et al. (2021), is employed, which combines the mean absolute error (MAE) with the structural (dis-)similarity

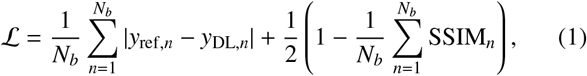

where *y*_ref,*n*_ represents the intensity of the pseudo-ground truth DTI map at position *n* estimated with the NLLS method, *y*_DL,*n*_ is the corresponding predicted intensity with the neural network, and *N*_*b*_ is the total number of samples in a batch. The SSIM_*n*_ denotes the structural similarity index (Wang et al., 2004) computed over a local window centered at pixel *n*

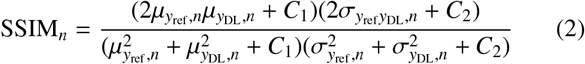

with *µ*_*y* ref, *n*_ and *µ*_*y* DL, *n*_ being the sample first-order moments (i.e., local means) computed within a fixed spatial window around the pixel *n*, 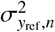 and 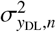 are the corresponding sample variances, and *σ*_*y* ref *y*DL, *n*_ is the sample covariance, *C*_1_ and *C*_2_ are the stability constants defined as *C*_1_ = (*k*_1_*L*)^2^ and *C*_1_ = (*k*_2_*L*)^2^, where *L* is the dynamic range of the pixel-values, and *k*_1_ = 0.01 and *k*_2_ = 0.03. The SSIM parameter is computed using a uniform sliding window of size 7 ×7.

The more similar the pseudo-ground-truth reference and predicted maps are, the closer the structural similarity index is to unity. Thus, our cost function given by Eq. (1) minimizes the Quantity 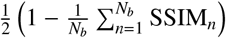, which we call the structural dissimilarity (DSSIM) index. The DSSIM assesses differences in structural information, such as local contrast and texture. We note that while the MAE may oversmooth the estimated maps by focusing solely on pixel-wise differences, incorporating the DSSIM term helps to preserve anatomical features, edges, and fine-grained details, which are critical for clinical interpretation.

The training parameters for all model instances were as follows: number of epochs: 200, batch size: 8, learning rate: 0.001, and NAdam optimizer (Dozat, 2016). Instead of selecting the model from the final training epoch, we selected the model checkpoints corresponding to the lowest validation loss for all analyses. This selection strategy minimizes the influence of potential training instabilities or overfitting in later epochs, ensuring that estimation accuracy is evaluated using the bestperforming model.

The presented network architecture was implemented using the PyTorch library v.2.5.1 (https://pytorch.org/).

### 2.6. Weights initialization

The weights of DL model instances were initialized using the He uniform initialization scheme (He et al., 2015). This initialization has been designed to preserve the variance of activations and gradients across neural network layers, thereby mitigating vanishing and exploding gradient problems during the training procedure (Glorot and Bengio, 2010). The initial weights for each run were sampled from a uniform distribution *U* as:

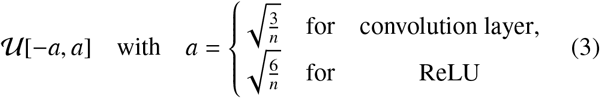

with *n* being the number of input units to the layer. The parameters in Eq. (3) come from a theoretical formula for the variance of a uniform distribution and enable maintaining a constant variance through activations and gradients across model layers.

To assess the replicability of DL-based DTI parameters, each model instance was trained 100 times independently for every acquisition scheme considered in the study, each using a different random seed for weight initialization. We call each trained network the “**run**”.

Random numbers were generated using the Mersenne Twister pseudorandom number generator (Matsumoto and Nishimura, 1998) using torch.manual_seed function delivered with the PyTorch library. The random seed initializes the generator and determines the sequence of random numbers used during weight initialization. As a result, although all neural networks were initialized using the same strategy and probability distribution, each training instance was initiated with a unique independent set of initial weights, preserving the theoretical statistical properties across runs.

### 2.7. Tract-based spatial statistics

The TBSS analysis of DTI parameters was performed using the FSL v6 software (Smith et al., 2006, 2004). First, the FA parameters for all subjects were aligned into a common space using a nonlinear registration following the standard procedure. Next, a mean FA image was generated and thinned to produce a skeleton representing the centres of all tracts common to the group. We applied a threshold of 0.2 to this skeleton to include major white matter bundles while excluding peripheral tracts. All aligned FA data were subsequently projected onto the mean FA skeleton. The registration and projection information obtained from the FA analysis was applied to the RD and FA. We conducted voxel-wise cross-subject statistics using the FSL randomise function (*p <* 0.05, 5000 permutations). The results were corrected for multiple comparisons with thresholdfree cluster enhancement (Winkler et al., 2014). Group differences between HC and patients with BD were assessed using a voxel-wise general linear model (GLM):

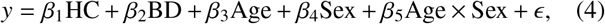

where *y* denotes the voxel-wise DTI parameter, *β*_*i*_ are the regression coefficients estimated by the model, and *ϵ* is the residual error. Age, sex, and their interaction (i.e., Age × Sex) were included as nuisance covariates to adjust for their potential effects on white matter measures. Regions of interest (ROIs) were defined as the intersection of the mean FA skeleton and the corresponding white matter labels from the Johns Hopkins University (JHU) white matter atlas (Mori et al., 2005).

### 2.8. ROI-based statistical analysis

Using the same ROIs as in the TBSS analysis, we computed mean DTI parameters for each subject by averaging voxel values within the corresponding ROI. These ROIs were defined as the intersection of the mean FA skeleton and the corresponding white matter labels from the JHU atlas. Similarly to the TBSS analysis, only voxels with the mean FA greater than 0.2 were included. Group differences between HC and BD were evaluated using a GLM, with age, sex, and their interaction (i.e., Age × Sex) included as nuisance regressors, consistent with the TBSS (see Eq. (4)). The fitted GLM produced an observed *t*statistic for the group contrast, indicating the magnitude of the adjusted group difference relative to its standard error.

### 2.9. Evaluation metrics

We assess predicted DL DTI parameters using two categories of metrics. The former includes image-based similarity metrics commonly used in natural image analysis, such as the mean squared error (MSE) and mean SSIM (MSSIM). These metrics quantify the voxel-wise agreement between the predicted and reference DTI parameters, though they do not directly indicate clinical relevance. The latter category includes TBSS and ROIbased analyses, which evaluate whether DL-predicted DTI parameters maintain group-level differences between BD patients and HC.

#### Image-based similarity metrics

Both metrics were computed within a brain mask only to exclude background voxels and to prevent artificially inflated values from large, homogeneous non-brain areas. The MSSIM was calculated as the mean value of the SSIM metric over the brain mask using the formula given in Eq. (2).

#### TBSS-based evaluation

We compared the parameters predicted by the DL model with those obtained from the pseudoground truth using standard NLLS estimation. To this end, we used two measures: TPR and FPR. Formally speaking, let *S*_ref_ represent the set of voxels displaying statistically significant group differences in the reference analysis with the NLLS method, and *S*_DL_ denote the corresponding set derived from the predicted DTI maps. The TPR is then defined as follows:

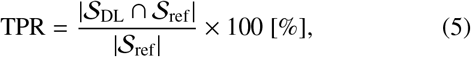

where | · | denotes set cardinality. Similarly, the FPR is given by

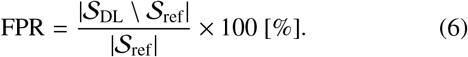

As the FPR is normalized by the number of significant voxels identified in the reference analysis, its value is not bounded by 100%. Consequently, FPR values greater than 100% indicate that the estimated analysis identified more false-positive voxels than there were significant voxels in the reference analysis. Fig. S1 in the Supplementary materials illustrates this.

ROI-based analysis: For the ROI-based statistical analysis, we obtain the standard *t*-statistic and the residualized Cohen’s *d* to quantify the magnitude of the adjusted group difference:

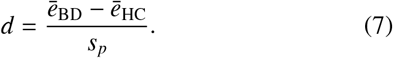

with the pooled standard deviation *s*_*p*_ of the residuals computed as

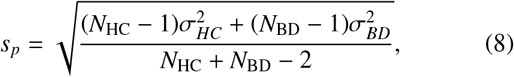

where 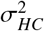 and 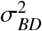 are the sample variances of the residuals in the HC and BD groups, respectively, and *N*_HC_ and *N*_BD_ denote the numbers of subjects in each group. The quantities *ē*_HC_ and *ē*_BD_ denote the mean residuals of the HC and BD groups

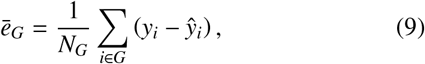

where *N*_*G*_ is the number of subjects in group *G* ∈ {{HC_1_, …, HC_18_}, {BD_1_, …, BD_18_}}, and *y*_*i*_ is the observed mean DTI metric value for *i*–th subject (reference one) and *ŷ*_*i*_ is the predicted metric by the reduced model, i.e., the model from Eq. (4) without group variables HC and BD. We note that Eq. (9) is computed separately for HD and BD groups.

To isolate the group effect from confounding variables, the reduced model includes only the intercept and nuisance covariates, and is formulated as:

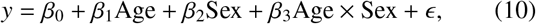

where *β*_0_ is the intercept.

## 3. Results

In this section, we present experimental results for DL-based DTI using all three configurations defined in section 2.3.

### 3.1. Between-group differences

We begin with a benchmark experiment to determine whether the initial resampling scheme affects the TBSS analysis. To this end, we performed a reference analysis with RD and FA maps estimated using the NLLS method from the resampled 64-direction dataset into the incremental gradient distribution scheme (**Configuration 1**). This analysis includes 18 individuals with BD and 18 HCs. We compare the statistically significant voxel-wise differences from the resampled 64-direction dataset with those obtained from the original 64-direction scheme. The results of this experiment are summarized in Table 1. The experiment with no resampling is presented in the Supplementary materials (see Table S1 and Table S4).

**Table 1:**
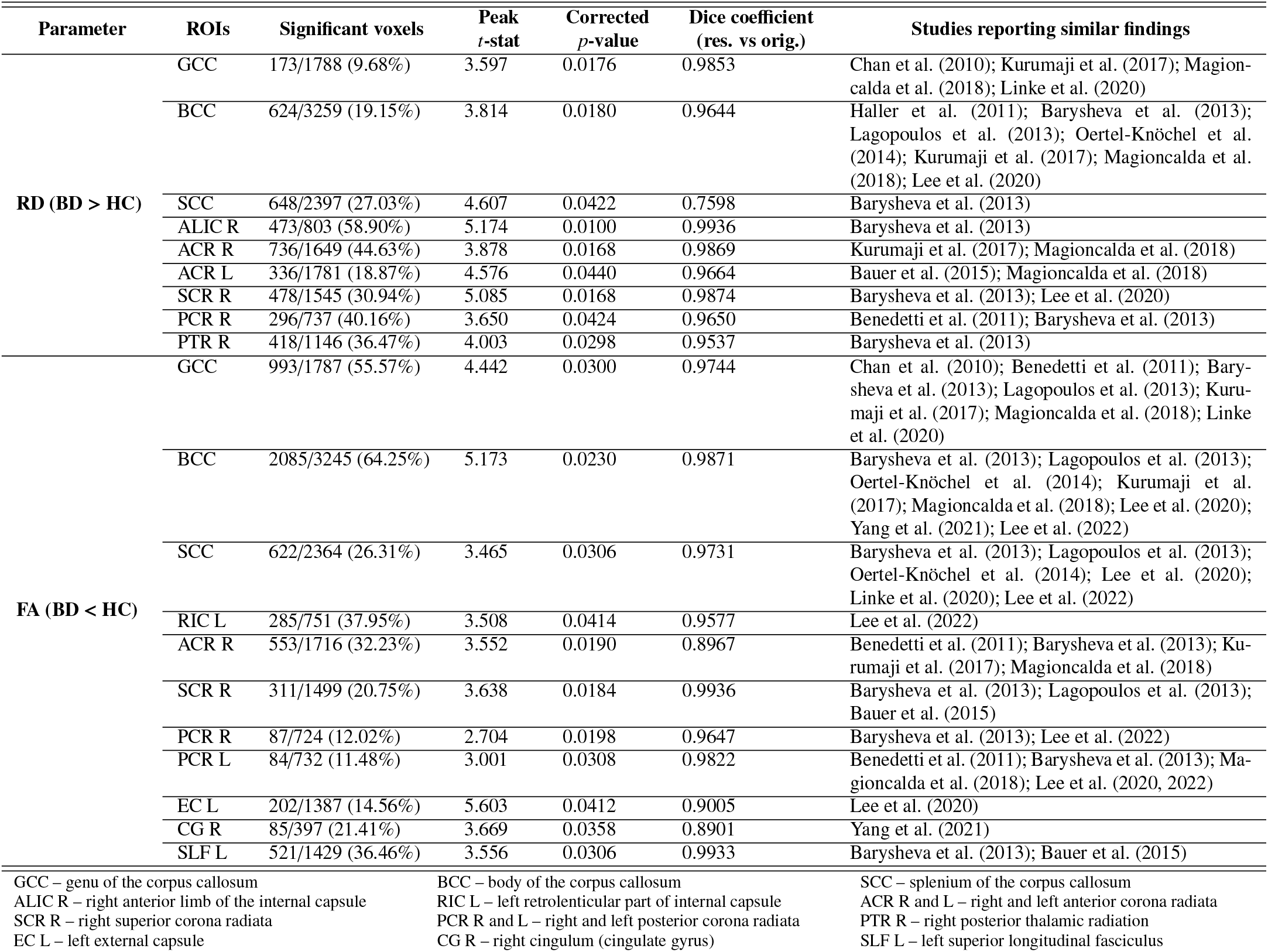
White matter regions identified by tract-based spatial statistics (TBSS) analysis of radial diffusivity (RD) and fractional anisotropy (FA) in patients with bipolar disorder (BD) compared with healthy controls (HC) for **Configuration 1**. The table includes only regions that show statistically significant results in RD (BD > HC) and FA (BD < HC) relative to HC after the threshold-free cluster enhancement (TFCE) correction (*p <* 0.05). Significant voxels are reported as *n/N* (%), where *n* is the number of voxels indicating a statistically significant difference, and *N* is the total number of voxels within the corresponding white matter region. Peak *t*-stat represents the maximum *t*-statistic within the significant portion of the region. The corrected *p*-value is the TFCE family-wise error-corrected *p*-value at the peak voxel. Dice coefficient (resampled versus original) measures the spatial overlap between significant voxels from the resampled 64-direction dataset (Configuration 1) and those from the original 64-direction dataset (Configuration 2) in the reference analysis. A Dice coefficient of 1 indicates complete spatial overlap, while 0 indicates no overlap.

| **Parameter** | **ROIs** | **Significant voxels** | **Peak *t*-stat** | **Corrected *p*-value** | **Dice coefficient (res. vs orig.)** | **Studies reporting similar findings** |
|---|---|---|---|---|---|---|
| **RD (BD > HC)** | GCC | 173/1788 (9.68%) | 3.597 | 0.0176 | 0.9853 | Chan et al. (2010); Kurumaji et al. (2017); Magion-calda et al. (2018); Linke et al. (2020) |
|  | BCC | 624/3259 (19.15%) | 3.814 | 0.0180 | 0.9644 | Haller et al. (2011); Barysheva et al. (2013); Lagopoulos et al. (2013); Oertel-Knöchel et al. (2014); Kurumaji et al. (2017); Magioncalda et al. (2018); Lee et al. (2020) |
|  | SCC | 648/2397 (27.03%) | 4.607 | 0.0422 | 0.7598 | Barysheva et al. (2013) |
|  | ALIC R | 473/803 (58.90%) | 5.174 | 0.0100 | 0.9936 | Barysheva et al. (2013) |
|  | ACR R | 736/1649 (44.63%) | 3.878 | 0.0168 | 0.9869 | Kurumaji et al. (2017); Magioncalda et al. (2018) |
|  | ACR L | 336/1781 (18.87%) | 4.576 | 0.0440 | 0.9664 | Bauer et al. (2015); Magioncalda et al. (2018) |
|  | SCR R | 478/1545 (30.94%) | 5.085 | 0.0168 | 0.9874 | Barysheva et al. (2013); Lee et al. (2020) |
|  | PCR R | 296/737 (40.16%) | 3.650 | 0.0424 | 0.9650 | Benedetti et al. (2011); Barysheva et al. (2013) |
|  | PTR R | 418/1146 (36.47%) | 4.003 | 0.0298 | 0.9537 | Barysheva et al. (2013) |
| **FA (BD < HC)** | GCC | 993/1787 (55.57%) | 4.442 | 0.0300 | 0.9744 | Chan et al. (2010); Benedetti et al. (2011); Bary-sheva et al. (2013); Lagopoulos et al. (2013); Kuru-maji et al. (2017); Magioncalda et al. (2018); Linke et al. (2020) |
|  | BCC | 2085/3245 (64.25%) | 5.173 | 0.0230 | 0.9871 | Barysheva et al. (2013); Lagopoulos et al. (2013); Oertel-Knöchel et al. (2014); Kurumaji et al. (2017); Magioncalda et al. (2018); Lee et al. (2020); Yang et al. (2021); Lee et al. (2022) |
|  | SCC | 622/2364 (26.31%) | 3.465 | 0.0306 | 0.9731 | Barysheva et al. (2013); Lagopoulos et al. (2013); Oertel-Knöchel et al. (2014); Lee et al. (2020); Linke et al. (2020); Lee et al. (2022) |
|  | RIC L | 285/751 (37.95%) | 3.508 | 0.0414 | 0.9577 | Lee et al. (2022) |
|  | ACR R | 553/1716 (32.23%) | 3.552 | 0.0190 | 0.8967 | Benedetti et al. (2011); Barysheva et al. (2013); Ku-rumaji et al. (2017); Magioncalda et al. (2018) |
|  | SCR R | 311/1499 (20.75%) | 3.638 | 0.0184 | 0.9936 | Barysheva et al. (2013); Lagopoulos et al. (2013); Bauer et al. (2015) |
|  | PCR R | 87/724 (12.02%) | 2.704 | 0.0198 | 0.9647 | Barysheva et al. (2013); Lee et al. (2022) |
|  | PCR L | 84/732 (11.48%) | 3.001 | 0.0308 | 0.9822 | Benedetti et al. (2011); Barysheva et al. (2013); Ma-gioncalda et al. (2018); Lee et al. (2020, 2022) |
|  | EC L | 202/1387 (14.56%) | 5.603 | 0.0412 | 0.9005 | Lee et al. (2020) |
|  | CG R | 85/397 (21.41%) | 3.669 | 0.0358 | 0.8901 | Yang et al. (2021) |
|  | SLF L | 521/1429 (36.46%) | 3.556 | 0.0306 | 0.9933 | Barysheva et al. (2013); Bauer et al. (2015) |

GCC – genu of the corpus callosum
BCC – body of the corpus callosum
SCC – splenium of the corpus callosum
ALIC R – right anterior limb of the internal capsule
RIC L – left retrolenticular part of internal capsule
ACR R and L – right and left anterior corona radiata
SCR R – right superior corona radiata
PCR R and L – right and left posterior corona radiata
PTR R – right posterior thalamic radiation
EC L – left external capsule
CG R – right cingulum (cingulate gyrus)
SLF L – left superior longitudinal fasciculus

We found statistically significant higher values in the BD group for RD parameter (BD > HC), with the most pronounced effects observed in the GCC, BCC, and SCC regions, as well as in the ALIC R, ACR R and ACR L, SCR R, PCR R, and PTR R. Similar findings have been already reported in previous studies using other BD datasets (Chan et al., 2010; Benedetti et al., 2011; Haller et al., 2011; Barysheva et al., 2013; Lagopoulos et al., 2013; Oertel-Knöchel et al., 2014; Bauer et al., 2015; Kurumaji et al., 2017; Magioncalda et al., 2018; Lee et al., 2020; Linke et al., 2020).

For the FA parameter, significantly lower values are observed in the BD group (BD < HC) within the GCC, BCC, and SCC, RIC L, ACR R, SCR R, PCR R, PCR L, EC L, CG R, and SLF L. These results are also consistent with previous reports (Chan et al., 2010; Benedetti et al., 2011; Barysheva et al., 2013; Lagopoulos et al., 2013; Oertel-Knöchel et al., 2014; Bauer et al., 2015; Kurumaji et al., 2017; Magioncalda et al., 2018; Lee et al., 2020; Linke et al., 2020; Yang et al., 2021; Lee et al., 2022).

The TBSS results for FA and RD demonstrate high consistency between the resampled 64-direction and the original 64direction dataset, as demonstrated by the Dice similarity coefficient. For most ROIs, the corresponding significance maps exhibit a Dice coefficient greater than 0.95. The only notable exception is the SCC region in the RD parameter, which shows a decrease to a Dice coefficient of 0.75. Overall, these findings suggest that the initial resampling has little effect on detecting group differences.

The identified ROIs in this analysis serve as the reference pattern for between-group differences and are also used in analyses based on RD and FA maps obtained from DL models.

### 3.2. Qualitative evaluation of estimated DTI maps

Before we proceed to key statistical experiments, we visually inspect RD and FA maps estimated using DL models trained separately for four acquisition schemes (64, 16, 6, and 3 gradient directions). This inspection checks whether the observations reported later in the TBSS and ROI-based analyses may be due to spurious peaks in the estimated parameters or estimation errors. Fig. 2 shows representative maps alongside their corresponding voxel-wise error maps, calculated as Parameter_ref_ −Parameter_DL_, with the reference estimated using a resampled 64-gradient scheme (**Configuration 1**).

**Figure 2:**
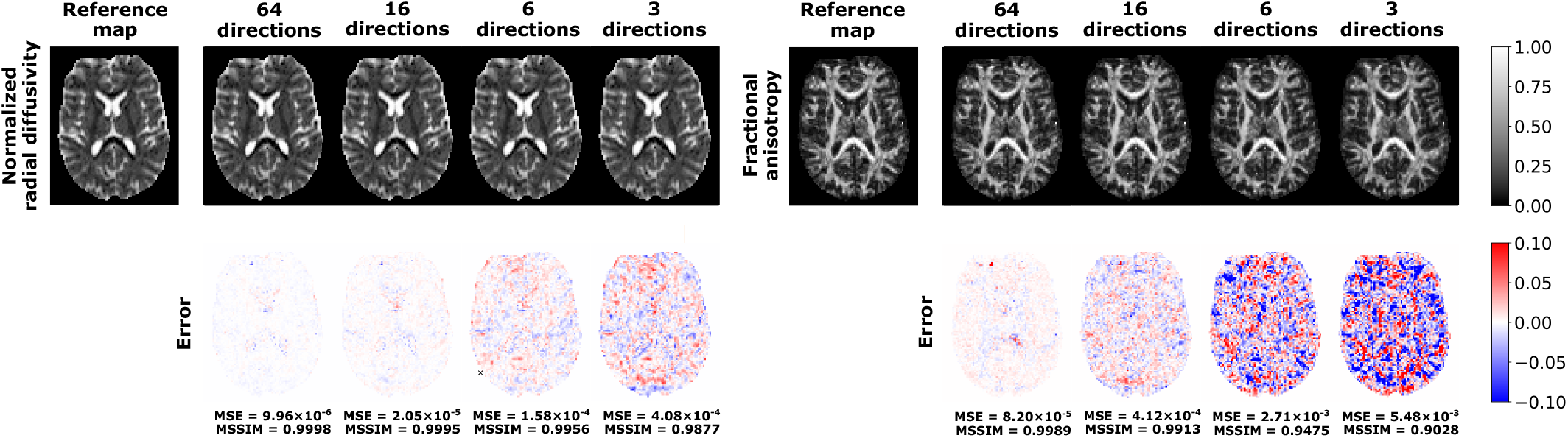
Visual inspection of **normalized radial di**ff**usivity** and **fractional anisotropy** estimated using single runs of the deep learning model and the corresponding voxel-wise differences between DL-based maps and the reference map derived from a resampled 64-direction scheme using the NLLS estimation (**Configuration 1**). Four input diffusion gradient schemes are considered for DL-based estimation: 64, 16, 6, and 3 directions. The brain-masked MSE and MSSIM values are reported for each estimated map.

Visually, the estimated RD maps closely resemble the reference maps for all four model instances. No obvious anatomical distortions or systematic estimation errors arise, even under the most limited gradient sampling scheme. A similar observation applies to FA, which also shows no evident systematic errors. However, the estimation error is larger than that observed for RD.

MSE analysis reveals that the estimation error increases as the number of gradient directions decreases, as expected. For the 64-and 16-direction schemes, the voxel-wise differences are generally small, with MSE below 10^−4^ for RD and below 10^−3^ for FA. For 6 gradient directions, the estimation error becomes more evident, with an increase in the MSE, which is approximately one order of magnitude higher than that obtained with 16 directions. This change is also reflected in the FA MSSIM value, which decreases from 0.99 to 0.95. The 3-direction scheme shows the largest estimation errors, with MSE values of approximately 4 ×10^−4^ (RD) and 5.5 ×10^−3^ (FA). Despite the larger errors observed for the most reduced scheme, they remain spatially dispersed rather than concentrated in specific brain regions.

### 3.3. Voxel-wise evaluation of deep learning-based predicted DTI maps

We now assess whether DL-predicted DTI parameters retain clinically relevant information. We trained separate DL models to estimate RD and FA for each gradient direction configuration using the BD training dataset described in Section 2.4. A total of 100 independent models were trained for each acquisition scheme, using different instances of model weights. We then applied the trained models to the 18 BD participants included in the reference analysis presented in Table 1. We kept the HC data from the reference NLLS-based estimation and applied the incremental 64-gradient scheme (**Configuration 1**).

Fig. 3 summarizes the results for the RD parameter within the BCC, including four different numbers of gradient directions (64, 16, 6, and 3). The histograms present voxels with statistically significant results in the reference analysis in blue and voxels with non-statistically significant results in orange. We selected three representative runs to show the range of variation observed in the statistical analyses. To further illustrate the problem, we selected runs with substantial underestimation of true positives (low TPR) and runs in which the model produced many false positives (high FPR).

**Figure 3:**
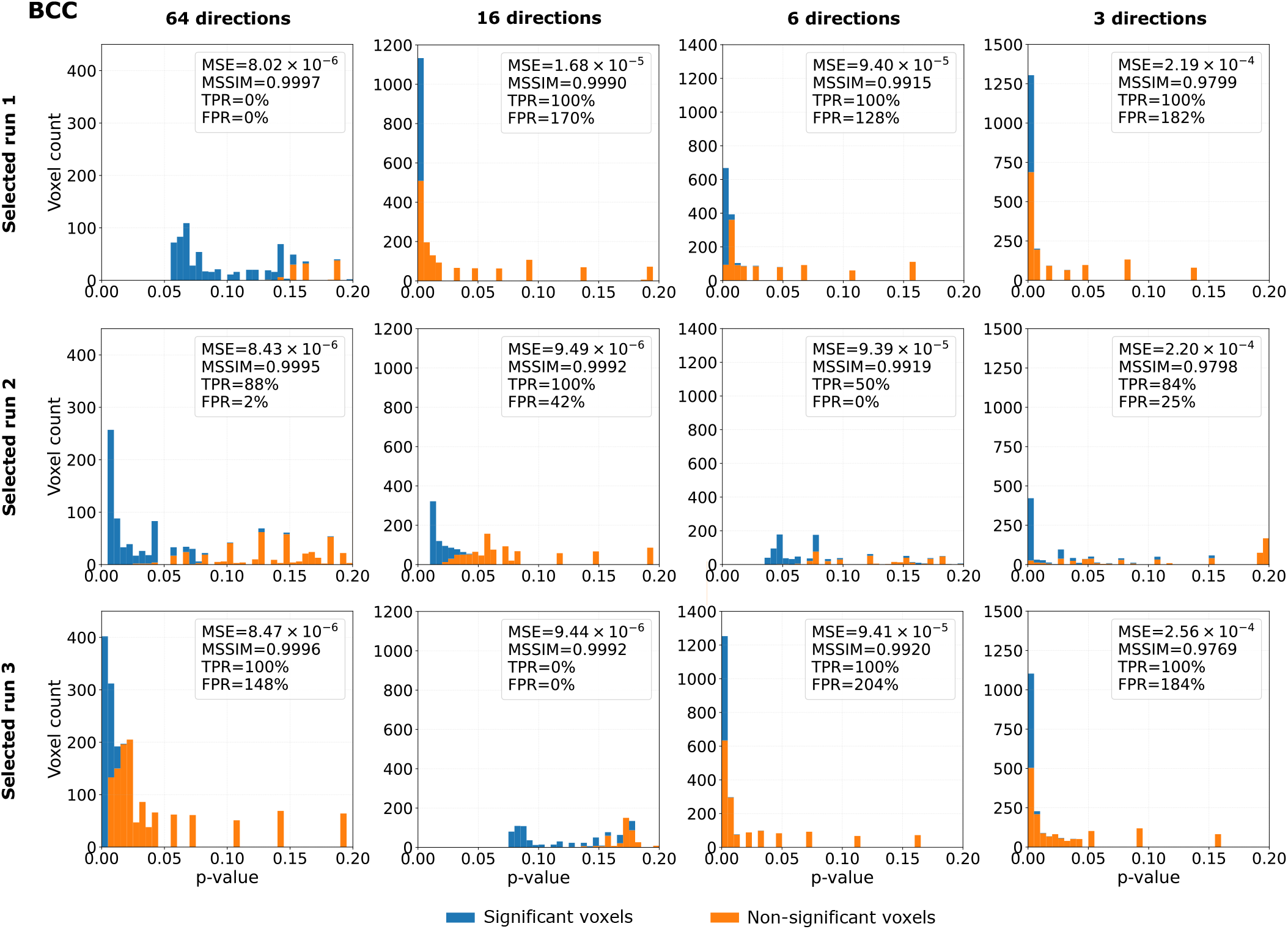
Histograms of voxel-wise *p*-values obtained from the tract-based spatial statistics (TBSS) analysis for the RD parameter estimated using a deep learning model following **Configuration 1**. The figure presents three selected runs out of 100 runs characterized with a comparable mean squared error (MSE) and mean structural similarity index measure (MSSIM), while varying true positive and false positive rates (TPR, FPR). For each run, we considered four diffusion gradient distributions of input data: 64, 16, 6, and 3. Blue bins indicate voxels that were statistically significant in the TBSS analysis based on standard NLLS-derived DTI (no deep learning applied), while orange bins indicate voxels that were statistically non-significant in the TBSS analysis based on NLLS-derived DTI.

These examples highlight a considerable discrepancy between group-level differences detected by TBSS and the preservation of image-quality metrics. Across all gradient schemes, the estimated RD parameters closely match the pseudo-groundtruth data. Even in the most challenging scenario, which in-volves 3 gradient directions, the MSE remains around 10^−4^, while MSSIM is close to 0.98. These results suggest that the DL models produce visually and quantitatively accurate RD estimates. In contrast, results derived from the TBSS analysis exhibit significant run-to-run variability. For instance, in the 64-gradient-directions variant, all three runs show nearly identical MSE and MSSIM values, while the corresponding TPR ranges from 0% to 100%, and the FPR from 0% to 148%. The 6-direction scheme shows an even more pronounced effect, with the FPR increasing from 0% to over 200% despite only small changes in MSE and MSSIM.

To better understand variability in TBSS results across runs, Fig. 4 shows TPR as a function of FPR across all 100 runs for the BCC and ACR R regions. For the BCC, the 64- and 16-direction schemes yield FPR values ranging from 0% to around 150%. As the number of gradient directions decreases, the FPR shifts toward higher values and becomes more concentrated. For example, for the 6- and 3-direction schemes, most runs yield FPR values between approximately 50% and 170%. A similar trend is observed for TPR. In the 64- and 16-direction schemes, TPR spans the full range from 0% to 100%, indicating considerable variability in preserving significant voxels across runs. Conversely, for the 6- and 3-direction schemes, the TPR distribution becomes more concentrated at higher values, with most runs falling between approximately 95% and 100%. Notably, for the 64- and 16-direction schemes, many runs cluster around two key boundaries: FPR at 0% or TPR at 100%. This pattern indicates that the RD values are either consistently underestimated or overestimated. As a result, the number of significant voxels uniformly decreases or increases. In contrast, only a small number of runs reveal a combination of high FPR and low TPR across all schemes, suggesting more selective misestimation of RD values.

**Figure 4:**
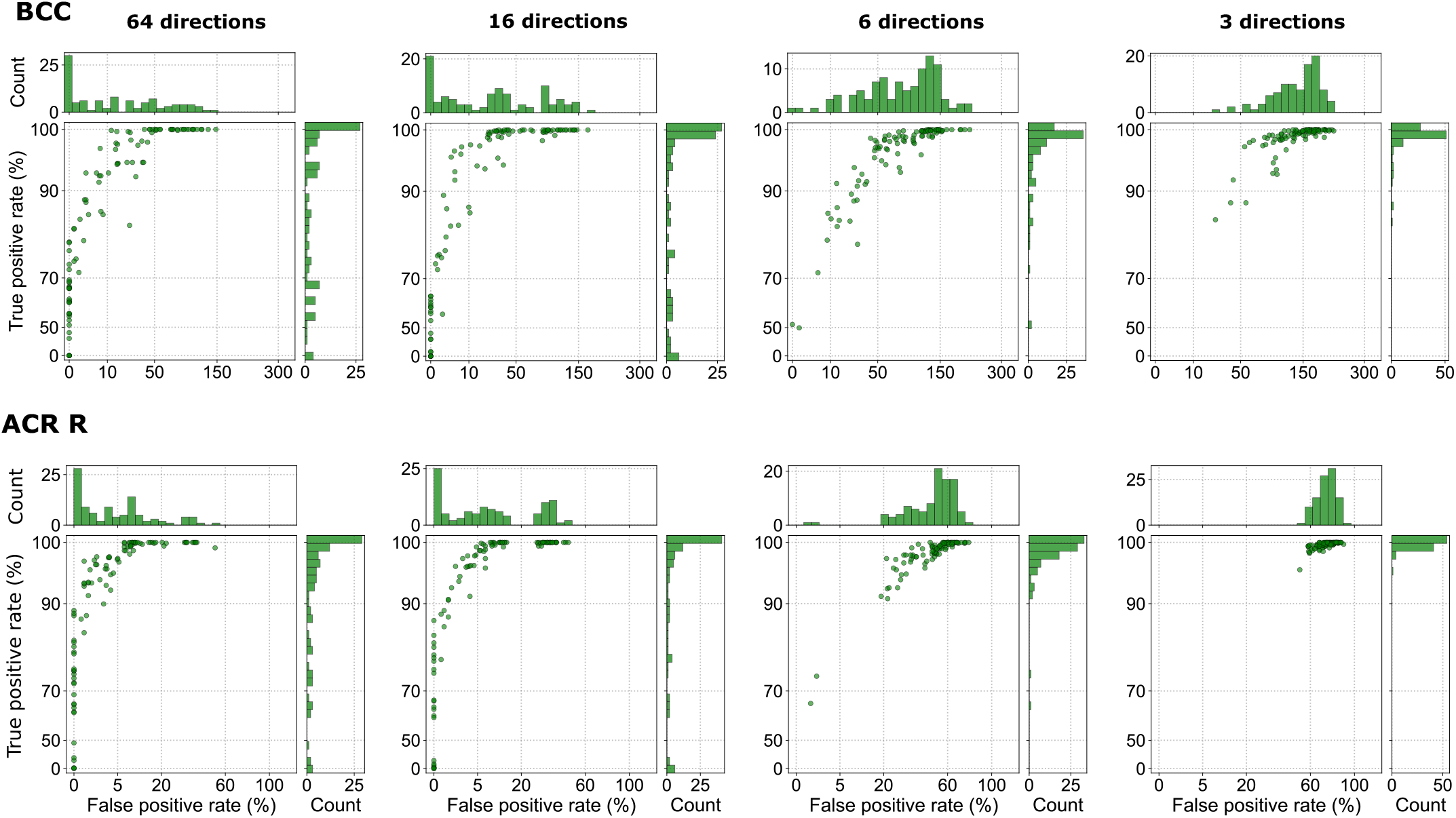
Relationship between the true positive rate and the false positive rate, obtained from the tract-based spatial statistics (TBSS) analysis for RD parameter, in the body of the corpus callosum (BCC) and right anterior corona radiata (ACR R) using **Configuration 1**, with four different numbers of input data gradient directions (64, 16, 6, and 3). Each point refers to a single run. In total, 100 runs were considered. The FPRs greater than 100% result from computing the values relative to the number of significant voxels identified in the reference analysis (see the explanation included in Fig. S1 of the Supplementary materials).

The patterns observed in the BCC also appear in ACR R, indicating that the behavior is not specific to a single ROI. The primary difference between the regions is the range of FPR values observed in the reduced schemes. These differences likely stem mainly from ROI size and the baseline number of statistically significant voxels identified in the reference analysis, rather than from region-specific differences in RD estimation accuracy. Since FPR was normalized by the number of voxels, regions with a smaller reference set are inherently more sensitive to extra significant voxels. As a result, comparable levels of overestimation can lead to considerably higher FPR values in BCC compared to ACR R.

To examine the spatial distribution of FP voxels identified in the TBSS analyses across different runs and acquisition schemes, we visualize their occurrence frequency within the BCC area in Fig. 5. The color indicates the number of runs in which a particular voxel is identified as statistically significant in the DL-based analysis but not in the reference analysis presented in Table 1.

**Figure 5:**
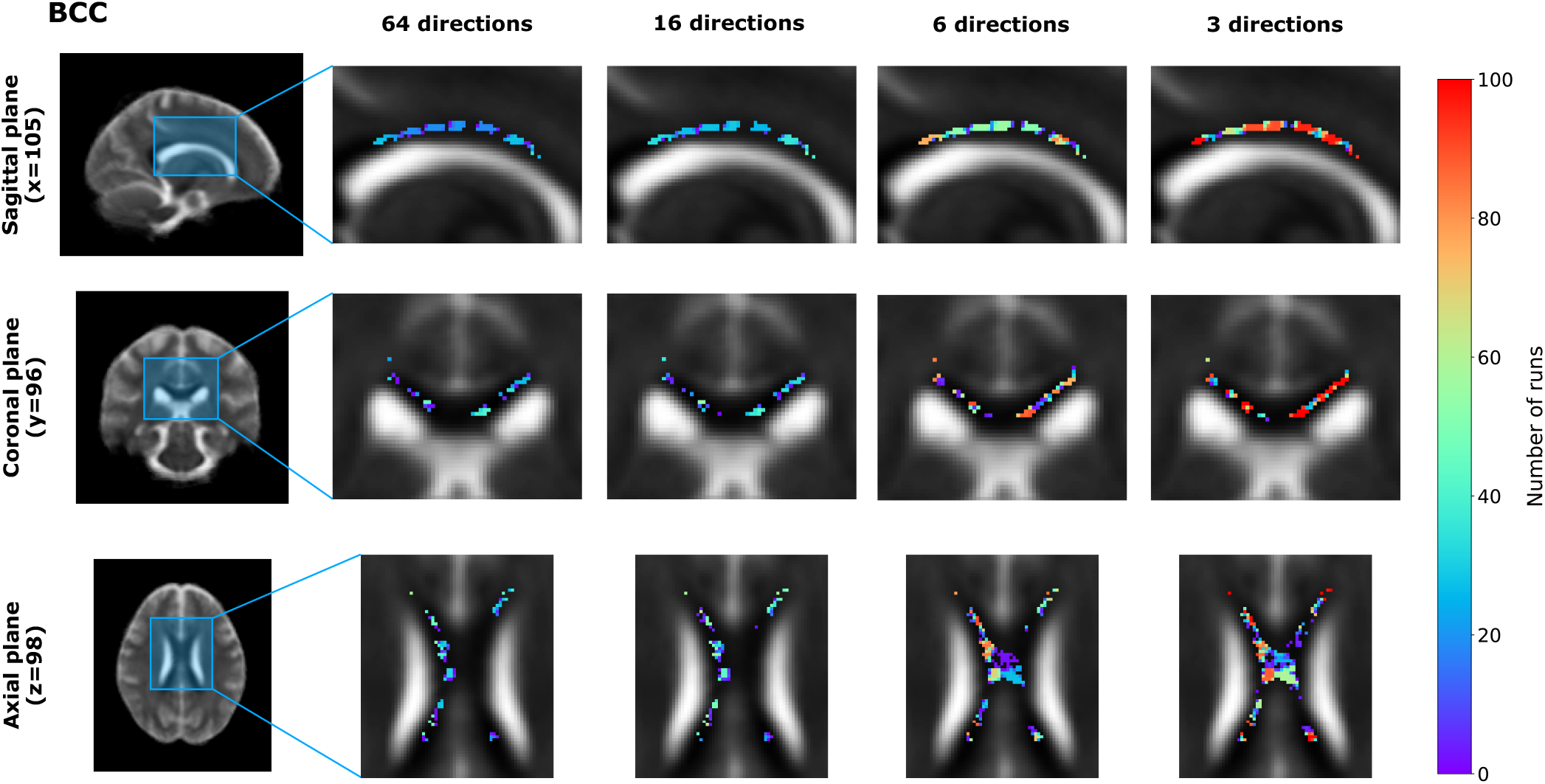
Visual representation of false positive voxels, obtained from the tract-based spatial statistics (TBSS) analysis for the RD parameter, in the body of the corpus callosum (BCC) shown in the sagittal, coronal, and axial planes for four input gradient directions: 64, 16, 6, and 3 (**Configuration 1**). False positive voxels correspond to locations identified as exhibiting statistically significant group differences in the estimated RD maps but not in the reference analysis. The color indicates how many runs classified a given voxel as a false positive.

The voxels incorrectly identified as statistically significant overlap spatially across training repetitions rather than occurring at entirely different locations in each run. Their occurrence frequency generally increases as the number of gradient directions decreases. Some voxels identified as FP in approximately 10–20 runs with 64 directions are identified in approximately 20–35, 50–70, and nearly all runs with 16, 6, and 3 directions, respectively (see the axial plane in Fig. 5). Additional FP voxels also emerge under reduced acquisition schemes. These results demonstrate that the spatial distribution of FP voxels varies with the acquisition scheme and that some locations are repeatedly identified as significant across runs.

Overall, the results presented in Figs. 3, 4, and 5 demonstrate that the statistical agreement between TBSS analyses performed on the estimated and reference RD maps can vary substantially across different runs, even when the corresponding image similarity metrics remain nearly identical. As the number of gradient directions decreases, the distribution of results shifts toward higher FPR values while generally preserving high TPR values.

The additional analyses included in the Supplementary materials demonstrate a consistent pattern across various white matter regions (see Figs. S2 and S3).

### 3.4. Comparison of three experimental configurations

To determine whether the findings under **Configuration 1** are specific to the selected resampling strategy and reference RD definition, we conduct auxiliary analyses using **Configuration 2** and **Configuration 3**. In the former scenario, the pseudoground truth RD maps employed for the DL training procedure are derived using the NLLS algorithm applied to the original 64-direction dataset rather than the resampled dataset. The latter configuration examines the stability of standard NLLSbased result estimation under different gradient-scheme reori-entations. The results for all three configurations are summarized in Fig. 6. To better visualize how TPR, FPR, MSE, and MSSIM depend on the number of gradient directions, we fitted second-order polynomial regression curves to the median values of each metric.

**Figure 6:**
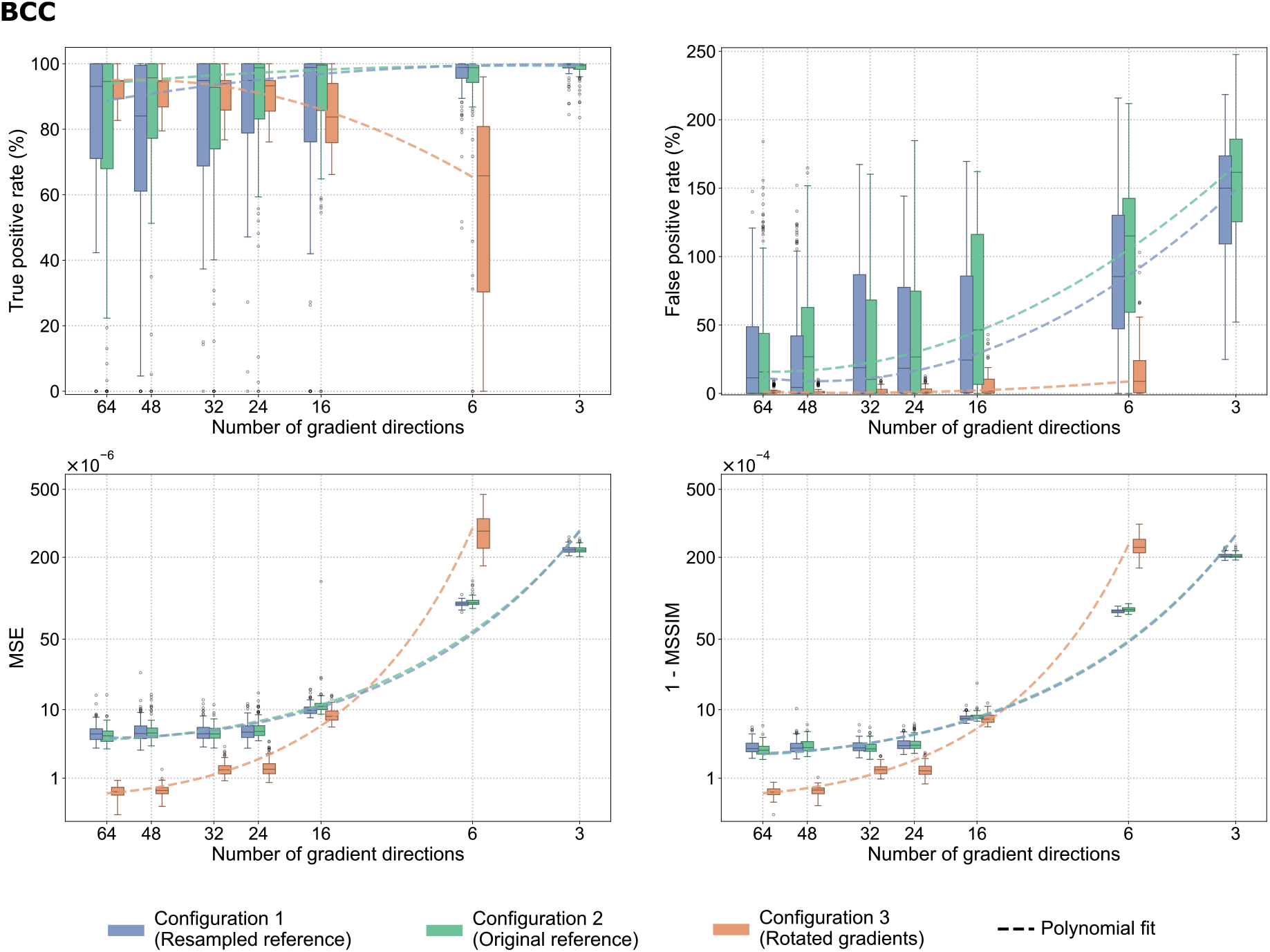
Boxplots presenting distributions of true positive rate, false positive rate, mean squared error (MSE), and 1 −mean structural similarity index measure (1 −MSSIM) for RD parameter across the body of the corpus callosum (BCC) for the three experimental configurations: **Configuration 1 (Resampled reference)** – reference maps were derived using the NLLS from the resampled 64-direction dataset, **Configuration 2 (Original reference)** – reference maps were derived using the NLLS from the original 64-direction dataset, **Configuration 3 (Rotated gradients)** – a non-DL baseline representing RD calculated using the NLLS fitting on rotated gradient sets. Boxes indicate the interquartile range (IQR), horizontal lines denote the median, whiskers extend to 1.5 ×IQR, and individual points indicate outliers. The dashed line represents a second-order polynomial regression fitted to the median values of the metrics, illustrating the overall trend of changes.

**Configuration 1** and **Configuration 2** show similar trends across all metrics, suggesting that the patterns observed in the previous analyses are unaffected by the choice of reference RD maps used for the training procedure. Although minor differences in the TPR and FPR distributions are noticeable between these two configurations, no consistent trend indicates that either configuration reliably reduces result variability. In both configurations, TPR increases significantly when the number of gradient directions is reduced to 6 or fewer. A corresponding rise in FPR also occurs, with the greatest increase in the 6- and 3-direction schemes. Similar trends appear for the two imagequality indices, MSE and (1 −MSSIM), indicating a gradual decline in image-level agreement with the reference RD maps as the number of gradient directions decreases.

In contrast, **Configuration 3** shows markedly different behavior, particularly in TPR and FPR. Under 24 or more gradient directions, the TPR distributions are less dispersed than those observed for the DL models (i.e., Configurations 1 and 2). However, when the number of gradient directions falls below 16, TPR variability increases significantly, and the median FPR remains close to 0% across all schemes. This indicates that reducing the number of gradient directions primarily affects the NLLS-based method’s ability to replicate the significant voxels identified in the reference analysis. The image similarity metrics further underscore the differences between the DL-based and NLLS approaches in obtaining the RD parameter. For gradient schemes with more than 16 directions, Configuration 3 achieves MSE and (1 − MSSIM) values roughly an order of magnitude lower than those for Configurations 1 and 2, showing substantially better agreement with the reference RD maps. However, for significantly reduced gradient schemes, the image similarity metrics decline much more rapidly for Configuration 3.

Results for other ROIs under Configurations 2 and 3 are provided in the Supplementary materials (see Figs. S6–S9 and S13–S16).

### 3.5. Preservation of group differences in ROI-based analysis

To complement the TBSS analysis and assess whether the observed trends were specific to the voxel-wise approach, we compared BD and HC groups using an ROI-based approach. The trained DL models were used to estimate the DTI parameters for the 18 BD participants, while the HC data were retained from the reference NLLS-based estimation using the resampled 64-gradient scheme (**Configuration 1**). We obtained mean parameter values within each ROI and calculated *t*-statistics using the same GLM and covariates as in the TBSS analysis (Eq. 4). Cohen’s *d* values were estimated using the reduced GLM defined in Eq. 10. Fig. 7 illustrates the distributions of the *t*-statistics and Cohen’s *d* values across 100 runs. The dashed line represents the corresponding values obtained from the reference analysis using the NLLS method. For selected ROIs, we compared the statistical significance from each of the 100 runs with the reference-analysis decision. The percentage of runs that result in the same decision as the reference analysis, i.e., both identifying the ROI as statistically significant or non-significant, is given as the *decision agreement* and reported in Table 2.

**Table 2:**
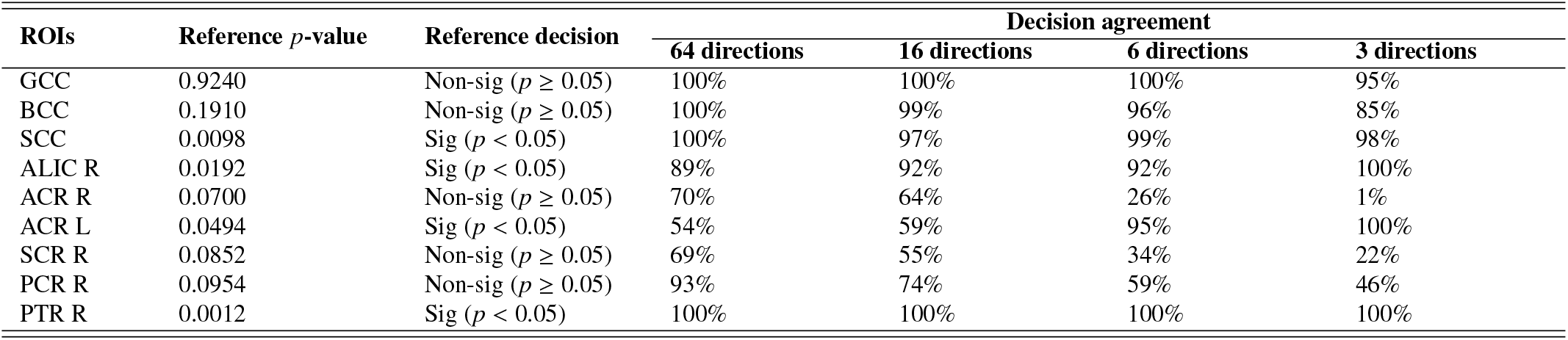
Decision agreement between DL-based and reference ROI analyses across four different numbers of input data gradient directions (64, 16, 6, and 3) for the RD (**Configuration 1**). The reference *p*-value and significance decision are reported for each ROI, while decision agreement indicates the percentage of runs out of 100 that reproduced the statistical significance of the reference analysis (*p <* 0.05 or *p* ≥ 0.05).

**Figure 7:**
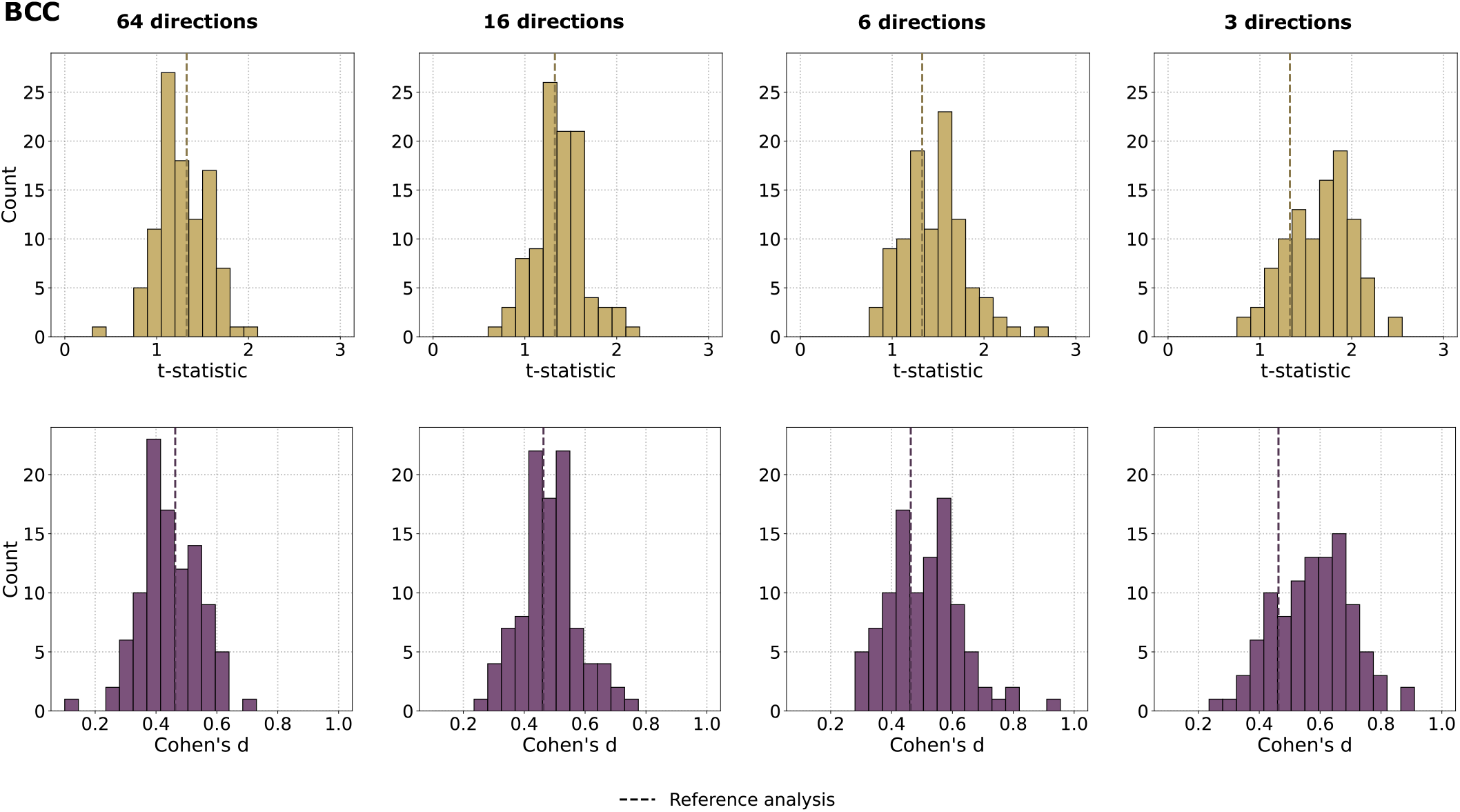
Distributions of *t*-statistics (top row) and Cohen’s *d* values (bottom row) in the body of the corpus callosum (BCC) for the RD parameter estimated using a deep learning model, with four different numbers of input data gradient directions (64, 16, 6, and 3) (**Configuration 1**). These statistics were calculated by comparing the mean RD values between patients with bipolar disorder (BD) and healthy controls (HC). Each histogram summarizes the results obtained from 100 runs. The dashed vertical line indicates the corresponding value from the reference analysis using the NLLS estimation.

The ROI-based analysis confirms the trends observed in the previous TBSS evaluation, although the variability is less pronounced. For both the *t*-statistic and Cohen’s *d*, the distributions obtained with the 64-direction scheme are approximately centered around the reference values. This observation aligns with the TBSS results, which indicate that independent runs tend to both underestimate and overestimate the statistical differences compared to the pseudo-ground truth analysis. As the number of gradient directions decreases, the distributions shift toward higher values. This shift is especially pronounced for the 6- and 3-direction cases, where higher *t*-statistics and Cohen’s *d* values are observed. Additionally, the distributions become increasingly asymmetric and exhibit greater variability. These findings demonstrate that the observed sensitivity to different runs and the number of gradient directions is not related to artefacts, but remains evident in ROI-based statistical comparisons. The Supplementary Materials further confirm this trend for additional ROIs (Fig. S4), where the shift in *t*-statistics and Cohen’s *d* toward higher values, as well as the increased asymmetry of the distributions, are much more pronounced.

In the *p*-value analysis, agreement between the DL-based and reference analyses varies across ROIs and gradient schemes. For GCC, BCC, SCC, ALIC R, and PTR R, agreement remain relatively high across all acquisition schemes, with at least 85% of runs matching the decision from the reference analysis. In contrast, ACR R demonstrate a significant decrease in agreement for a reduced number of gradient directions, falling from 70% agreement for the 64-direction scheme to only 1% for the 3-direction one. A similar trend is observed for SCR R and PCR R regions, although the reduction in decision agreement is less pronounced. In contrast, ACR L shows a different pattern: agreement is relatively low for 64 and 16 gradient directions but increases to 100% for the most reduced acquisition scheme. These results highlight that reducing the number of gradient directions does not consistently affect agreement with the reference, i.e., in some regions, we observe an increase in the number of matching runes, whereas in others, we observe the opposite effect.

The degree of agreement between the DL-based and reference analyses varies by region and reference *p*-value. For example, the reference *p*-value for SCC is well below the 0.05 significance threshold, and therefore, nearly all runs (across all acquisition schemes) reproduce the same decision. In contrast, the reference *p*-value for ACR R is 0.07, which is relatively close to the threshold, making the decision more sensitive to small variations in the estimated values.

### 3.6. Agreement between estimated and reference DTI parameters

To better understand the variability in our statistical analyses, we present a complementary experiment using Bland-Altman plots. We plot the mean RD values over the BCC region from the DL models across 100 runs against the results from the standard NLLS method applied to a resampled 64-direction dataset (**Configuration 1**). The results of this experiment are depicted in Fig. 8.

**Figure 8:**
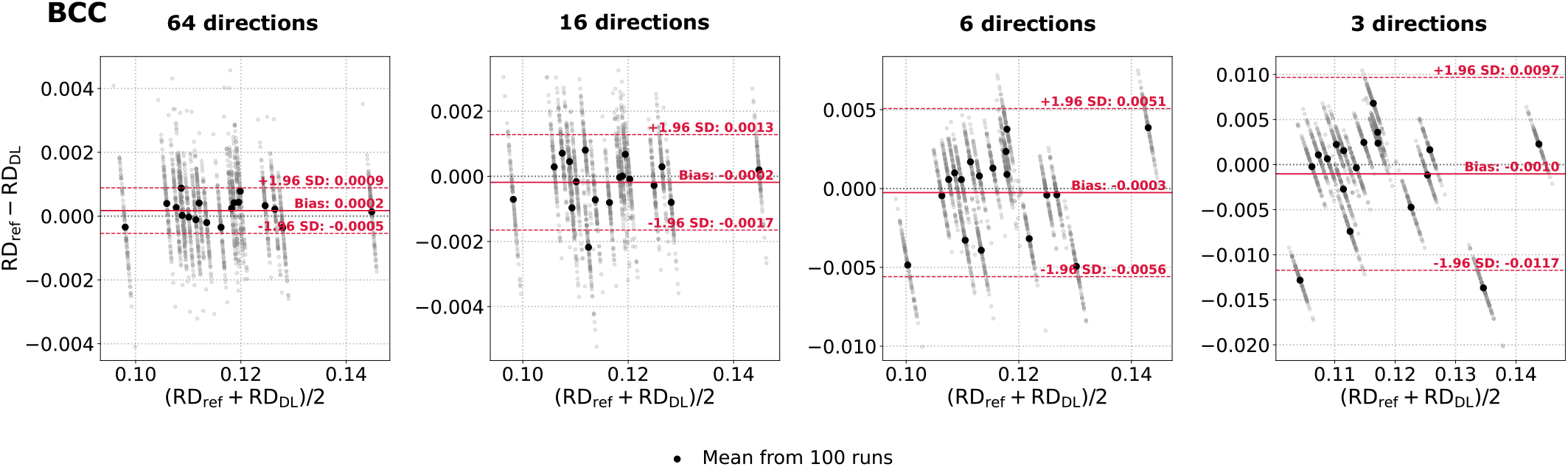
Bland-Altman plots of the RD parameter computed from the BCC region across all BD subjects. The reference values (RD_ref_) were obtained using the NLLS estimation from a resampled 64-direction scheme, whereas the predicted RD_DL_ values were generated using the deep learning model (**Configuration 1**) considering four different numbers of input data gradient directions (64, 16, 6, and 3). Gray points represent individual model runs (mean values over the BCC region). A single black point corresponds to the average across 100 runs for a single BD patient. Solid red lines indicate the bias, while dashed red lines denote 95% limits of agreement (±1.96 standard deviations).

The Bland-Altman plots illustrate significant variability across runs for the same BD patient (represented by diagonal lines). Beyond the overall variation, some run-subject combinations show substantial deviations from the corresponding averages (black dots), suggesting inherent variability in the DLbased predictions across runs rather than data-processing errors or model failures. As the number of gradient directions decreases, the variability in the estimated RD values increases notably. For the 64-direction scheme, the difference between the reference (estimated using the NLLS) and the DL model is around ±0.005. In contrast, for the 3-direction scheme, this range increases to approximately ±0.010. Besides, a systematic change in bias is observed. The mean bias is 0.0002 for the 64-direction scheme, decreases to −0.0003 for the 6-direction scheme, and declines further to approximately −0.0010 for the 3-direction scheme. This gradual shift toward negative bias indicates that DL instances tend to yield progressively higher RD values than the reference maps as the number of gradient directions decreases.

### 3.7. Evaluation of deep learning-based fractional anisotropy maps estimation

To determine whether the trends observed in the previous analyses are specific to RD or generalize to other DTI parameters, we conduct further analyses using FA maps. Fig. 9 replicates the experiment previously presented in Fig. 3 for FA within the SCR R region. For each subsampling scheme, we selected three runs to illustrate variability in the statistical results. Again, voxels identified as statistically significant in the reference analysis are marked in blue, while voxels with no significant group differences are in orange. The selected runs clearly show that the trend observed for the RD parameter is also present for FA. A similar discrepancy between the imagequality metrics and the TBSS results is observed. MSE and MSSIM values remain nearly identical across individual runs. In contrast, the TPR and FPR vary substantially between individual runs for the same acquisition scheme. This variability is particularly evident for the 16-direction scheme, for which TPR ranges from 49% to 100% and FPR from 0% to nearly 200%.

**Figure 9:**
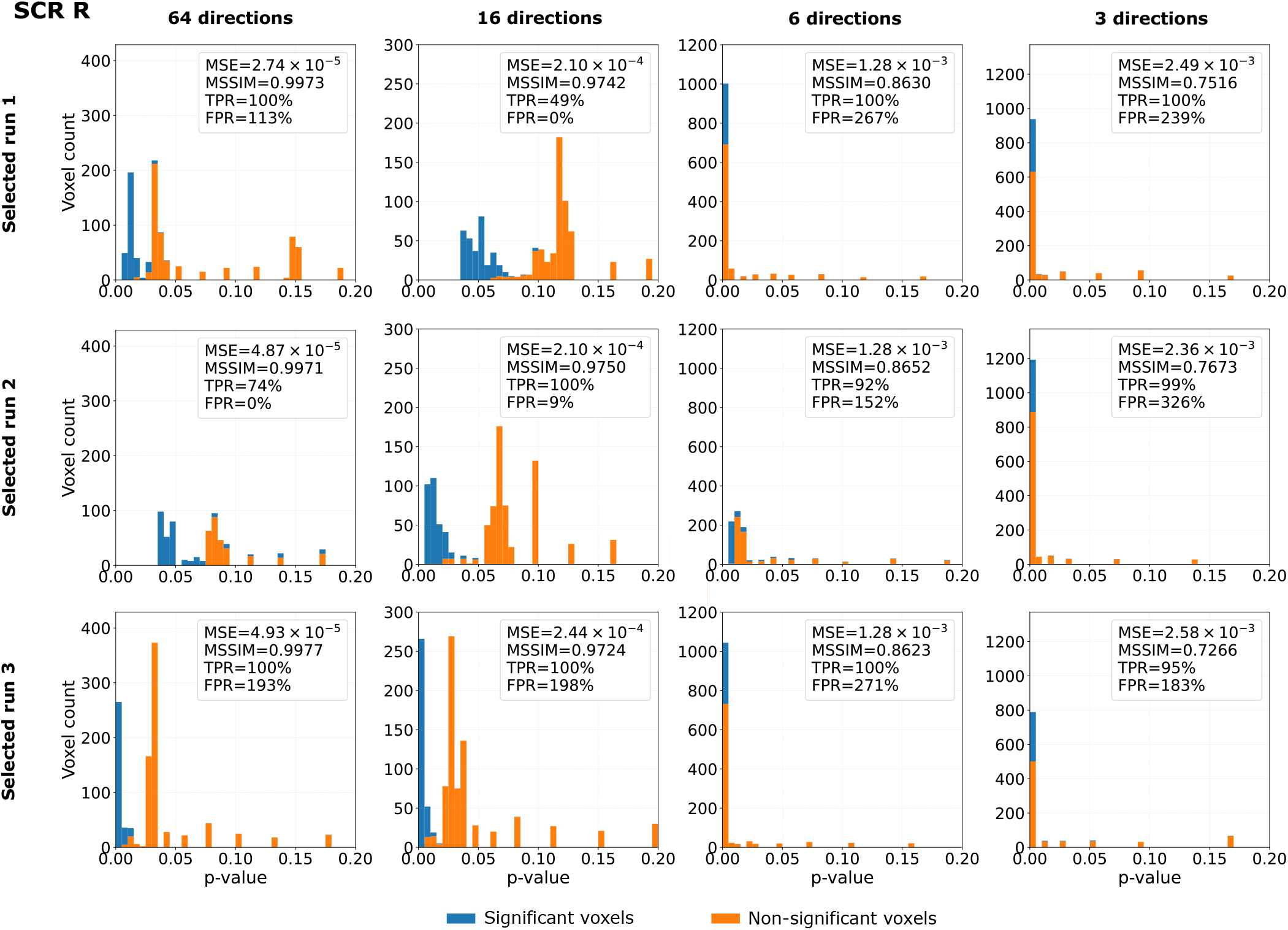
Histograms of voxel-wise *p*-values obtained from the TBSS analysis using the FA parameter over the skeleton of SCR R region for three selected deep learning training runs (**Configuration 1**). For each run, four diffusion gradient distributions of input data are considered: 64, 16, 6, and 3. The *p*-values are computed relative to the reference results derived from parametric DTI. For each run, we show the MSE, MSSIM, TPR, and FPR for the FA parameter estimated using a deep learning model. Blue-colored bins refer to statistically significant voxels, while orange bins refer to statistically non-significant voxels estimated from parametric DTI.

These trends are also evident in the extended analyses, including extra ROIs presented in the Supplementary materials (see Figs. S20-S24).

## 4. Discussion

Interest in employing DL methods to estimate DTI-based parameters has recently grown immensely. Despite this trend, DL models are often trained only on data from healthy individuals (Goodwin-Allcock et al., 2022; Martin et al., 2025; Ametepe et al., 2026; Chen et al., 2026), even when they are evaluated on a clinical cohort (Li et al., 2021; Gaviraghi et al., 2022; AjaFernández et al., 2023; Gaviraghi et al., 2024).

In this work, we systematically explored the clinical relevance of DL-based DTI parameters in BP, a condition in which disease-related microstructural differences can be relatively subtle. Our primary focus was retraining a model with different weight initializations and evaluating whether it preserves clinically significant information. Besides, we examined whether this effect changes with reduced acquisition schemes. Model retraining and its clinical validation in quantitative MRI imaging have received significantly less attention in the literature. We note that the objective of the study was not to develop a novel DL-based method that achieves the minimal possible error between pseudo-ground truth and estimated maps or generalizes across different populations, but rather to verify the correctness and consistency of DTI measures generated using the DL models from a clinical perspective in a run-to-run scenario.

### 4.1. Evaluation of deep learning-based models

The experimental results depicted in Fig. 3 indicate that different DL model runs, despite similar image-quality metrics achieved for the predicted RD parameters (e.g., MSE and MSSIM), can yield substantially different TBSS results. In some cases, the estimated maps showed a high degree of agreement with the reference analysis in terms of statistically significant voxels (e.g., run 2 for 64 gradient directions, with a TPR of 88% and an FPR of 2%). In other cases, however, similar image-level agreement was associated with markedly different statistical outcomes, ranging from nearly the entire BCC being identified as statistically significant (e.g., run 1 for 16-direction scheme) to no statistically significant voxels identified (e.g., run 3 for 16-direction scheme).

These findings highlight a key limitation of image-level metrics, which are typically used to construct loss functions and evaluate DL models for quantitative imaging. In numerous studies, the training objective is based on metrics such as MSE or MSSIM, which encourage the estimated maps to closely match the reference at the image level (Golkov et al., 2016; Gibbons et al., 2019; Aliotta et al., 2021; Li et al., 2021; Park et al., 2021; Goodwin-Allcock et al., 2022; Gaviraghi et al., 2022, 2024; Martin et al., 2025; Chen et al., 2026). Model performance is then commonly evaluated using the same or similar measures (Aliotta et al., 2021; Park et al., 2021; GoodwinAllcock et al., 2022; Gaviraghi et al., 2022, 2024; Martin et al., 2025; Ametepe et al., 2026; Chen et al., 2026). Although these metrics quantify the overall agreement between estimated and (pseudo-)reference DTI parameters, they do not directly assess whether clinically relevant statistical differences between groups are preserved (Aja-Fernández et al., 2023). Consequently, maps with similar MSE and MSSIM values can yield substantially different patterns of statistically significant voxels. Conversely, local differences that contribute to image-level error may have little effect on group comparisons if they occur outside the ROIs included in the TBSS analysis, such as in grey matter. This distinction is particularly important for DTI, where even small, spatially distributed differences within the regions can affect the statistical significance of individual voxels (Ball et al., 2013; Bach et al., 2014; Maximov et al., 2015). Therefore, high image similarity should not be interpreted as evidence that the statistical properties of the estimated maps have been preserved or that the model is suitable for clinical analyses.

Our study revealed a discrepancy between conventional image-level metrics and downstream statistical analyses, aligning with results from other research involving various neural network architectures or clinical conditions, such as migraine (Aja-Fernández et al., 2023), temporal lobe epilepsy, and multiple sclerosis (Gaviraghi et al., 2024). Importantly, these studies used different training populations than our work: DL models were trained exclusively on healthy controls, whereas our neural networks were trained exclusively on patients with BD. Despite these differences in the training data, we observed similar trends in the relationship between image-level agreement and downstream statistical results. This consistency across datasets and patient populations suggests this limitation is not data-dependent. Instead, it may also apply to DL-based DTI parameter estimation in other clinical contexts.

### 4.2. Variability in deep learning-based DTI parameters estimation due to model weight initialization

We trained our models using the identical dataset, preprocessing pipeline, network architecture, training hyperparameters, and data partitioning approach. We also controlled other potential sources of variability in the training procedure to ensure reproducibility, including deterministic computational settings and consistent data ordering. Under such a controlled scenario, the only variability introduced to our experimental design between runs was the weights initialization in a pseudo-random manner using the He scheme (He et al., 2015). However, variability in weights initialization translated to significant variability in the results of the downstream TBSS analysis in both DTI parameters. To our knowledge, this behavior has not been previously noted in quantitative MRI, such as diffusion-weighted MR imaging. In particular, for acquisition schemes with more gradient directions, TPR ranged from 0% to 100% across runs, while FPR varied from 0% to several dozen or several hundred per cent, depending on the ROI, as illustrated in Fig. 4. The substantial differences in FPR among ROIs primarily stem from how the FPR was normalized using the number of statistically significant voxels in the reference analysis. When a large proportion of voxels within an ROI are statistically significant, the FPR remains relatively low. Conversely, when only a small proportion of voxels are significant, even a small number of falsepositive voxels can lead to a vastly higher FPR. These findings demonstrate that differences arising solely from the initial weight configuration can substantially affect the preservation of statistically significant group effects. Our results suggest that a single trained model may not fully capture the range of statistical effects from DL-based DTI parameters.

Variability in model weight initialization adds another layer of uncertainty to the estimated DTI parameters, which, importantly, affects the statistical conclusions drawn from these estimates. Even when both MSE and MSSIM indicate strong agreement with the reference parameters, different model initializations can yield substantially different results in TBSS analysis. This variation can range from closely preserving the reference group differences to entirely failing to detect them, or even mistakenly identifying additional areas as statistically significant. Consequently, considering a DL-based DTI estimated from a single run with a fixed initialization may lead to an incomplete and potentially misleading evaluation of model performance. Instead, evaluating multiple runs provides a more complete picture of this uncertainty, enabling a more robust assessment of the consistency of the estimated parameters and their impact in a clinical context.

### 4.3. Preservation of statistical properties in deep learning predicted DTI parameters

A key characteristic of DL-based DTI inference is its ability to preserve a large number of statistically significant voxels identified in the reference analysis. However, we also observed additional statistically significant voxels in the subsampled datasets. The DL-based estimates generally maintain the voxels identified as significant in the reference analysis while also identifying larger clusters of statistically significant voxels. Notably, these extra statistically significant voxels are not randomly distributed across the runs. Although their exact spatial locations vary from run-to-run, they consistently appear in the same anatomical regions, and their occurrence generally increases as the number of gradient directions decreases. This observation suggests that the extra statistical effects are not merely due to random voxel-wise variations but may reflect systematic characteristics of the DL-based estimation. The spatial distribution of these additional effects could be influenced by local diffusion-signal properties, signal-to-noise ratio, anatomical boundaries, or spatial features learned by the DL model. Therefore, a high TPR does not confirm that the statistical information in the reference maps has been fully preserved, as the presence of additional spatially structured statistically significant voxels can alter the extent of the observed group differences.

### 4.4. Differences in DTI parameters estimated using deep learning and NLLS

The effect of reducing the number of gradient directions reduction on DTI has been well documented in previous studies (Jones, 2004; Ni et al., 2006; Landman et al., 2007; Giannelli et al., 2010; Lebel et al., 2012). In addition, the benefits of DL-based approaches for purely image-quality metrics under reduced acquisition schemes have also been studied (Gibbons et al., 2019; Aliotta et al., 2021; Li et al., 2021; Park et al., 2021; Goodwin-Allcock et al., 2022; Gaviraghi et al., 2022, 2024; Martin et al., 2025). Our findings further demonstrate that NLLS and neural networks exhibit different responses to progressively reduced numbers of gradient directions, as illustrated in Fig. 6. Hence, reducing the amount of acquired information does not lead to the same type of degradation observed in these two approaches.

For the NLLS method, image-level metrics remained relatively stable across different numbers of gradient directions. A significant increase in error occurred only with the most reduced acquisition schemes, particularly with 6 gradient directions. In contrast, TBSS results remained consistent across most acquisition schemes, i.e., FPR was low, suggesting that NLLS rarely introduced extra statistically significant voxels not found in the reference analysis. The largest loss of agreement occurred with the most reduced scheme, reflected in a decrease in TPR. This indicates that fewer of the reference statistically significant voxels were identified.

In comparison, DL-based estimation displayed considerable variability in maintaining downstream statistical results, even when all 64 gradient directions were available. As the number of directions decreased, this variability increased and was associated with detecting additional statistically significant voxels, as indicated by higher FPR. Meanwhile, the TPR remained relatively high, and its variability decreased with fewer gradient directions. The Bland-Altman analysis for BD patients (Fig. 8, Fig. S5 in Supplementary materials) supports these findings and may indicate differing directional biases between the two approaches. For NLLS, the loss of statistically significant voxels under highly reduced acquisition schemes corresponds to underestimating RD relative to the reference. In the case of DL models, the combination of a high TPR and increasing FPR, along with observed shifts in RD values, suggests an overestimation of RD. These interpretations should be viewed as potential explanations for the observed statistical patterns, not as evidence of systematic bias in every voxel.

The observed difference between the two approaches (i.e., NLLS vs. DL-based) highlights different failure characteristics depending on whether the diffusion acquisition data are relatively complete or substantially reduced. When more gradient directions are available, DL-based DTI parameter estimation may not provide a clear advantage over the conventional NLLS method. NLLS produced more stable downstream statistical results, while DL models introduced greater variability. Consequently, the complexity of DL-based estimation does not seem to offer a significant benefit within this range of gradient directions, while also introducing additional uncertainty. This uncertainty specifically refers to the variability in results when retraining models with different initial weights. One run may produce results that accurately match the reference statistical voxels, while another run, trained with the same experimental setup, might overlook most reference group differences or identify far more statistically significant voxels.

As the number of gradient directions decreases, comparing the results becomes more complicated. Similar errors at the image level do not necessarily reflect similar statistical behavior. A DL model might produce visually or numerically similar parameter maps from significantly reduced acquisition data, but this should not be taken as evidence that it has preserved the statistical properties of the full data. These findings indicate that the choice between DL-based methods and conventional NLLS estimation should be based not only on the number of available gradient directions and the resulting image similarity but also on the intended downstream analysis and the required level of analysis reproducibility for the specific clinical application.

### 4.5. Dataset size

The UCLA Consortium for Neuropsychiatric Phenomics MRI dataset (Poldrack et al., 2016) contains a limited number of subjects, particularly in the BD group, which includes only 48 participants after excluding one subject due to poor data quality. As a result, a compromise had to be found between the size of the training set and the independent evaluation set used for the TBSS analyses. Therefore, we chose a 30:18 split, as we believed it provided sufficient data to train the neural networks while keeping the evaluation cohort large enough to identify between-group differences.

Given the available sample size, identifying a single subset of 18 BD subjects that reveals statistically significant differences across multiple ROIs for both DTI parameters was challenging. This is consistent with previous studies reporting variability in statistically significant findings across DTI measures (Chan et al., 2010; Benedetti et al., 2011). As the aim of this study was to investigate how DL-based parameter estimation affects subsequent statistical analyses, rather than to optimize dataset partitioning or TBSS sensitivity, separate evaluation subsets were selected for RD and FA. This allowed us to examine the downstream effects of DL-based estimation in the presence of between-group differences for each parameter.

The statistical power of TBSS depends on sample size (Bach et al., 2014), and a substantially larger cohort (i.e., more than 100 subjects per group) would enhance sensitivity and robustness. To ensure that the observed trends were not solely due to a limited sample size or specific to the TBSS framework, we performed two additional complementary analyses. First, we conducted ROI-wise analyses based on mean DTI parameters, which are less sensitive to sample size than voxel-wise TBSS. Second, we directly compared the estimated and reference DTIbased parameters using Bland-Altman plots (see Fig. 8). Despite employing different evaluation strategies, all three analyses showed consistent results. This agreement suggests that the observed effects cannot be attributed solely to the limited dataset size or to the characteristics of the TBSS methodology, but instead reflect the properties of the DL-based estimation.

### 4.6. Selection of microstructural parameters

The UCLA Consortium for Neuropsychiatric Phenomics MRI dataset includes DW MRI data acquired with a single *b*value of 1000 s/mm^2^. Consequently, our analyses were limited to parameters derived from the single-shell solutions, such as DTI, since more advanced signal representations or biophysical models, e.g., the Mean Apparent Propagator MRI (Özarslan et al., 2013) or the Neurite Orientation Dispersion and Density Imaging (Zhang et al., 2012), require multi-shell acquisitions to obtain meaningful parameters. Moreover, DTI has been widely used to investigate white matter abnormalities in BD, providing a strong basis for comparison with previous studies (Chan et al., 2010; Benedetti et al., 2011; Haller et al., 2011; Barysheva et al., 2013; Lagopoulos et al., 2013; Oertel-Knöchel et al., 2014; Bauer et al., 2015; Kurumaji et al., 2017; Magioncalda et al., 2018; Lee et al., 2020; Linke et al., 2020).

Among the DTI-derived parameters, we focused in this work on RD and FA. We excluded axial diffusivity (AD) because previous studies on BD have reported both increased (OertelKnöchel et al., 2014; Kurumaji et al., 2017; Singh et al., 2023) and decreased AD tendencies (Lan et al., 2020; Linke et al., 2020; Singh et al., 2023). We also excluded mean diffusivity (MD), as it did not show statistically significant differences between groups in the UCLA dataset when the subgroups were randomly selected for the TBSS analyses.

### 4.7. Selection of data for training and TBSS analysis

In this study, we trained the DL models using only experimental group data, unlike (Aja-Fernández et al., 2023), which used the HC group exclusively. Moreover, by restricting both the training and evaluation data to the same clinical population, we minimised variance that might come from inter-scanner differences in data acquisition protocols (Ciupek et al., 2026; Fiszer et al., 2026). For the TBSS analyses, we intentionally kept the HC measures as the original NLLS-derived pseudoground truth maps, while for the BD group, the metrics were replaced with DTI maps estimated using DL models. This design allowed us to assess how DL-based DTI parameters affect the detection of group differences. Therefore, any observed differences between the two TBSS results reflect changes in the estimated parameters for the BD group. If both groups were replaced with DL-based estimates, systematic biases, such as consistent overestimation or underestimation of parameters, could affect both groups similarly. This could make it more difficult to determine how the estimation process itself affected the observed group differences. Keeping the HC group unchanged therefore provides a controlled scenario in which the effect of DL-based DTI parameters in the TBSS results can be examined directly.

### 4.8. Selection of neural network architecture

In this work, we employed the U-Net architecture rather than exploring recent advances in the field or proposed modifications to existing schemes (Chen et al., 2026; Jiang et al., 2026). The U-Net architecture (Ronneberger et al., 2015) is well established in medical imaging and remains one of the most widely used architectures for estimating DTI parameters from DW MRI data (Aliotta et al., 2021; Li et al., 2021; Gaviraghi et al., 2022, 2024). The architecture provides a favorable balance between estimation accuracy and computational efficiency, which was essential in our study, as we trained over 1000 network instances to conduct a comprehensive set of analyses. Despite recent attention to advanced generative architectures, such as generative adversarial networks (Yi et al., 2019) or diffusion models (Kazerouni et al., 2023), their significantly higher computational requirements make them less suitable for exhaustive studies, as demonstrated in this work. For example, diffusion models require iterative denoising during both training and inference, resulting in considerably longer execution times (Croitoru et al., 2023; Taragola et al., 2026). Recent reports suggest that diffusion models may require approximately 14 times longer training and up to 60 times longer inference than U-Net (Taragola et al., 2026). This makes them unsuitable for the scale of analysis presented in this work, even when employing high-performance computing infrastructure.

Another advantage of U-Net architecture is that it directly learns a deterministic mapping from the input DW MR images to the target parameter metrics. In contrast, GANs optimized with distribution-matching objectives may be more prone to hallucinations (Cohen et al., 2018). Strictly speaking, GANbased approaches may generate structures not present in the input data or suppress existing anatomical features. Although these artifacts have been studied primarily in synthesis or translation medical imaging tasks (Cohen et al., 2018; Xin et al., 2020; Zoghby et al., 2024), they are also unwelcome in parameter estimation, where even subtle alterations of the estimated maps may bias downstream statistical analyses. This issue is particularly relevant to our analysis, where differences in DTI parameters are often subtle and spatially localized.

### 4.9. Generalization to other clinical applications

The results presented in this work are based on a cohort of individuals with BD. This raises a question of whether the observed trends are data-specific or reflect more general characteristics of DL-based DTI estimation. To address this concern, we conducted additional analyses with schizophrenia patients and presented the findings in the Supplementary materials (see section 4 therein). We observed run-to-run variability in the estimated parameters and their statistical properties in the groupbased study, similar to those observed in BD. In particular, TPR and FPR varied significantly across runs, despite comparable MSE and MSSIM values. In addition, reduced acquisition schemes displayed a trend toward higher TPR and FPR values. The ROI-based analysis exhibited a similar pattern, with *t*-statistics and Cohen’s *d* values clustering around the reference values for 64 gradient directions but increasingly shifting away from them under reduced acquisition schemes.

We also observed some differences between the two disorders. In schizophrenia, the metrics derived from TBSS showed greater variability for the 3 gradient directions. These differences suggest that the distribution of the observed effects may depend, to some extent, on the underlying data and the characteristics of each disease. Nevertheless, the key properties of DL-based metrics observed in the BD cohort were reproduced in schizophrenia, supporting the hypothesis that the statistical variability observed between runs is not condition-specific.

## 5. Conclusions

Our findings show that agreement at the image level does not necessarily preserve statistical results in downstream analyses when the neural network is trained multiple times under the same configuration setup. Although DL-based DTI estimation can remain comparable to reference parameters, different runs can produce substantially dissimilar statistical results, particularly when the amount of data used for inference is reduced. This variability suggests that conventional image similarity metrics may not adequately capture the relevance of DLbased DTI parameters for repeated statistical analyses. When DL-based DTI estimation is intended for studies of conditions characterized by subtle microstructural alterations, model evaluation should therefore include application-specific analyses in addition to conventional image similarity metrics.

## Acknowledgements

This work is supported by the European Union’s Horizon 2020 research and innovation programme under grant agreement No 857533 and the International Research Agendas Programme of the Foundation for Polish Science No MAB PLUS/2019/13. This work was conducted within the project of the Ministry of Science and Higher Education “Support for the activity of Centers of Excellence established in Poland under Horizon 2020” on the basis of the contract number MEiN/2023/DIR/3796. Tomasz Pieciak and Antonio Tristán Vega acknowledge the Agencia Estatal de Investigación (Ministerio de Ciencia, Innovación y Universidades of Spain) with the research grant PID2024-158963NB-I00.

The numerical experiments were possible through computing allocation on the Ares and Athena systems at the highperformance computing ACC Cyfronet AGH centre under the grants PLG/2025/018009 and PLG/2026/019157.

## References

Aja-Fernández, S., Martín-Martín, C., Planchuelo-Gómez, Á., Faiyaz, A., Uddin, M.N., Schifitto, G., Tiwari, A., Shigwan, S.J., Singh, R.K., Zheng, T., et al., 2023. Validation of deep learning techniques for quality augmentation in diffusion MRI for clinical studies. NeuroImage: Clinical 39, 103483.

Alexander, A.L., Lee, J.E., Lazar, M., Field, A.S., 2007. Diffusion tensor imaging of the brain. Neurotherapeutics 4, 316–329.

Aliotta, E., Nourzadeh, H., Patel, S.H., 2021. Extracting diffusion tensor fractional anisotropy and mean diffusivity from 3-direction DWI scans using deep learning. Magnetic Resonance in Medicine 85, 845–854.

Ametepe, J.M., Gholam, J., Beltrachini, L., Cercignani, M., Jones, D.K., 2026. Four directions, one solution: Enabling rapid diffusion tensor MRI for ultra-low field using deep learning. Magnetic Resonance in Medicine 96, 1413–1426.

Bach, M., Laun, F.B., Leemans, A., Tax, C.M., Biessels, G.J., Stieltjes, B., Maier-Hein, K.H., 2014. Methodological considerations on tract-based spatial statistics (TBSS). Neuroimage 100, 358–369.

Ball, G., Boardman, J.P., Arichi, T., Merchant, N., Rueckert, D., Edwards, A.D., Counsell, S.J., 2013. Testing the sensitivity of tract-based spatial statistics to simulated treatment effects in preterm neonates. PLoS One 8, e67706.

Barysheva, M., Jahanshad, N., Foland-Ross, L., Altshuler, L.L., Thompson, P.M., 2013. White matter microstructural abnormalities in bipolar disorder: A whole brain diffusion tensor imaging study. NeuroImage: clinical 2, 558–568.

Basser, P.J., Jones, D.K., 2002. Diffusion-tensor MRI: theory, experimental design and data analysis–a technical review. NMR in Biomedicine: An International Journal Devoted to the Development and Application of Magnetic Resonance In Vivo 15, 456–467.

Basser, P.J., Mattiello, J., LeBihan, D., 1994. Mr diffusion tensor spectroscopy and imaging. Biophysical journal 66, 259–267.

Basser, P.J., Pierpaoli, C., 1998. A simplified method to measure the diffusion tensor from seven MR images. Magnetic resonance in medicine 39, 928–934.

Bauer, I.E., Ouyang, A., Mwangi, B., Sanches, M., Zunta-Soares, G.B., Keefe, R.S., Huang, H., Soares, J.C., 2015. Reduced white matter integrity and verbal fluency impairment in young adults with bipolar disorder: a diffusion tensor imaging study. Journal of psychiatric research 62, 115–122.

Benedetti, F., Yeh, P.H., Bellani, M., Radaelli, D., Nicoletti, M.A., Poletti, S., Falini, A., Dallaspezia, S., Colombo, C., Scotti, G., et al., 2011. Disruption of white matter integrity in bipolar depression as a possible structural marker of illness. Biological psychiatry 69, 309–317.

Carreira Figueiredo, I., Borgan, F., Pasternak, O., Turkheimer, F.E., Howes, O.D., 2022. White-matter free-water diffusion MRI in schizophrenia: a systematic review and meta-analysis. Neuropsychopharmacology 47, 1413–1420.

Chan, W.Y., Yang, G.L., Chia, M.Y., Woon, P.S., Lee, J., Keefe, R., Sitoh, Y.Y., Nowinski, W.L., Sim, K., 2010. Cortical and subcortical white matter abnormalities in adults with remitted first-episode mania revealed by Tract-Based Spatial Statistics. Bipolar Disorders 12, 383–389.

Chen, Y., Li, Z., Wang, Y., Li, Y., Zheng, J., Yang, H., Liu, M., Cukur, T., Fan, Q., Li, Z., et al., 2026. MicroKAN: Mapping human brain microstructure using diffusion MRI and adaptive nonlinear modeling. NeuroImage 337, 122032.

Ciupek, D., Malawski, M., Pieciak, T., 2026. Federated learning: A new frontier in the exploration of multi-institutional medical imaging data. Computer Methods and Programs in Biomedicine 284, 109454.

Cohen, J.P., Luck, M., Honari, S., 2018. Distribution matching losses can hallucinate features in medical image translation, in: International conference on medical image computing and computer-assisted intervention, Springer. pp. 529–536.

Cox, S.R., Ritchie, S.J., Tucker-Drob, E.M., Liewald, D.C., Hagenaars, S.P., Davies, G., Wardlaw, J.M., Gale, C.R., Bastin, M.E., Deary, I.J., 2016. Ageing and brain white matter structure in 3,513 UK Biobank participants. Nature communications 7, 13629.

Croitoru, F.A., Hondru, V., Ionescu, R.T., Shah, M., 2023. Diffusion models in vision: A survey. IEEE transactions on pattern analysis and machine intelligence 45, 10850–10869.

Dozat, T., 2016. Incorporating Nesterov Momentum into Adam, in: Proceedings of the 4th International Conference on Learning Representations, pp. 1–4.

Eggl, M.F., De Santis, S., 2026. Simulation-based inference at the theoretical limit for fast, robust microstructural MRI with minimal diffusion data. Communications Medicine.

Fillard, P., Pennec, X., Arsigny, V., Ayache, N., 2007. Clinical DT-MRI estimation, smoothing, and fiber tracking with log-Euclidean metrics. IEEE transactions on medical imaging 26, 1472–1482.

Fiszer, J., Ciupek, D., Malawski, M., Pieciak, T., 2026. Validation of eleven federated learning strategies for multi-contrast image-to-image MRI data synthesis from heterogeneous sources. Biomedical Signal Processing and Control 119, 109649.

Garyfallidis, E., Brett, M., Amirbekian, B., Rokem, A., Van Der Walt, S., Descoteaux, M., Nimmo-Smith, I., Contributors, D., 2014. Dipy, a library for the analysis of diffusion MRI data. Frontiers in neuroinformatics 8, 8.

Gaviraghi, M., Ricciardi, A., Palesi, F., Brownlee, W., Vitali, P., Prados, F., Kanber, B., Gandini Wheeler-Kingshott, C.A., 2022. A generalized deep learning network for fractional anisotropy reconstruction: Application to epilepsy and multiple sclerosis. Frontiers in neuroinformatics 16, 891234.

Gaviraghi, M., Ricciardi, A., Palesi, F., Brownlee, W., Vitali, P., Prados, F., Kanber, B., Gandini Wheeler-Kingshott, C.A., 2024. Finding the limits of deep learning clinical sensitivity with fractional anisotropy (FA) microstructure maps. Frontiers in reuroinformatics 18, 1415085.

Giannelli, M., Cosottini, M., Michelassi, M.C., Lazzarotti, G., Belmonte, G., Bartolozzi, C., Lazzeri, M., 2010. Dependence of brain DTI maps of fractional anisotropy and mean diffusivity on the number of diffusion weighting directions. Journal of applied clinical medical physics 11, 176–190.

Gibbons, E.K., Hodgson, K.K., Chaudhari, A.S., Richards, L.G., Majersik, J.J., Adluru, G., DiBella, E.V., 2019. Simultaneous NODDI and GFA parameter map generation from subsampled q-space imaging using deep learning. Magnetic resonance in medicine 81, 2399–2411.

Glorot, X., Bengio, Y., 2010. Understanding the difficulty of training deep feedforward neural networks, in: Proceedings of the thirteenth international conference on artificial intelligence and statistics, JMLR Workshop and Conference Proceedings. pp. 249–256.

Golkov, V., Dosovitskiy, A., Sperl, J.I., Menzel, M.I., Czisch, M., Sämann, P., Brox, T., Cremers, D., 2016. Q-space deep learning: twelve-fold shorter and model-free diffusion MRI scans. IEEE transactions on medical imaging 35, 1344–1351.

Goodwin-Allcock, T., McEwen, J., Gray, R., Nachev, P., Zhang, H., 2022. How can spherical cnns benefit ml-based diffusion mri parameter estimation?, in: Cetin-Karayumak, S., Christiaens, D., Figini, M., Guevara, P., Pieciak, T., Powell, E., Rheault, F. (Eds.), Computational Diffusion MRI, Springer Nature Switzerland, Cham. pp. 101–112.

Grande, I., Berk, M., Birmaher, B., Vieta, E., 2016. Bipolar disorder. The Lancet 387, 1561–1572.

Haller, S., Xekardaki, A., Delaloye, C., Canuto, A., Lövblad, K.O., Gold, G., Giannakopoulos, P., 2011. Combined analysis of grey matter voxel-based morphometry and white matter tract-based spatial statistics in late-life bipolar disorder. Journal of Psychiatry and Neuroscience 36, 391–401.

He, K., Zhang, X., Ren, S., Sun, J., 2015. Delving deep into rectifiers: Surpassing human-level performance on imagenet classification, in: Proceedings of the IEEE international conference on computer vision, pp. 1026–1034.

Jenkinson, M., Bannister, P., Brady, M., Smith, S., 2002. Improved optimization for the robust and accurate linear registration and motion correction of brain images. Neuroimage 17, 825–841.

Jiang, H., Chen, G., Ma, J., Cui, H., Zhang, S., Xia, Y., Yap, P.T., 2026. Precise estimation of tissue microstructure with hybrid graph transformer. Artificial Intelligence in Medicine 176, 103390.

Jones, D.K., 2004. The effect of gradient sampling schemes on measures derived from diffusion tensor MRI: a Monte Carlo study. Magnetic Resonance in Medicine 51, 807–815.

Kazerouni, A., Aghdam, E.K., Heidari, M., Azad, R., Fayyaz, M., Hacihaliloglu, I., Merhof, D., 2023. Diffusion models in medical imaging: A comprehensive survey. Medical image analysis 88, 102846.

Kellner, E., Dhital, B., Kiselev, V.G., Reisert, M., 2016. Gibbsringing artifact removal based on local subvoxel-shifts. Magnetic resonance in medicine 76, 1574–1581.

Kim, M.E., Gao, C., Ramadass, K., Newlin, N.R., Kanakaraj, P., Bogdanov, S., Rudravaram, G., Archer, D., Hohman, T.J., Jefferson, A.L., et al., 2026. White matter micro-and macrostructure brain charts for the human lifespan. Nature, 1–11.

Koay, C.G., Chang, L.C., Carew, J.D., Pierpaoli, C., Basser, P.J., 2006. A unifying theoretical and algorithmic framework for least squares methods of estimation in diffusion tensor imaging. Journal of magnetic resonance 182, 115–125.

Koshiyama, D., Fukunaga, M., Okada, N., Morita, K., Nemoto, K., Usui, K., Yamamori, H., Yasuda, Y., Fujimoto, M., Kudo, N., et al., 2020. White matter microstructural alterations across four major psychiatric disorders: mega-analysis study in 2937 individuals. Molecular psychiatry 25, 883–895.

Kubicki, M., McCarley, R., Westin, C.F., Park, H.J., Maier, S., Kikinis, R., Jolesz, F.A., Shenton, M.E., 2007. A review of diffusion tensor imaging studies in schizophrenia. Journal of psychiatric research 41, 15–30.

Kurumaji, A., Itasaka, M., Uezato, A., Takiguchi, K., Jitoku, D., Hobo, M., Nishikawa, T., 2017. A distinctive abnormality of diffusion tensor imaging parameters in the fornix of patients with bipolar ii disorder. Psychiatry Research: Neuroimaging 266, 66–72.

Lagopoulos, J., Hermens, D.F., Hatton, S.N., Tobias-Webb, J., Griffiths, K., Naismith, S.L., Scott, E.M., Hickie, I.B., 2013. Microstructural white matter changes in the corpus callosum of young people with bipolar disorder: a diffusion tensor imaging study. PloS one 8, e59108.

Lan, M.J., Rubin-Falcone, H., Sublette, M.E., Oquendo, M.A., Stewart, J.W., Hellerstein, D.J., McGrath, P.J., Zanderigo, F., Mann, J.J., 2020. Deficits of white matter axial diffusivity in bipolar disorder relative to major depressive disorder: no relationship to cerebral perfusion or body mass index. Bipolar Disorders 22, 296–302.

Landman, B.A., Farrell, J.A., Jones, C.K., Smith, S.A., Prince, J.L., Mori, S., 2007. Effects of diffusion weighting schemes on the reproducibility of DTI-derived fractional anisotropy, mean diffusivity, and principal eigenvector measurements at 1.5 T. Neuroimage 36, 1123–1138.

Le Bihan, D., Mangin, J.F., Poupon, C., Clark, C.A., Pappata, S., Molko, N., Chabriat, H., 2001. Diffusion tensor imaging: concepts and applications. Journal of Magnetic Resonance Imaging 13, 534–546.

Lebel, C., Benner, T., Beaulieu, C., 2012. Six is enough? comparison of diffusion parameters measured using six or more diffusion-encoding gradient directions with deterministic tractography. Magnetic resonance in medicine 68, 474–483.

Lee, D.K., Lee, H., Park, K., Joh, E., Kim, C.E., Ryu, S., 2020. Common gray and white matter abnormalities in schizophrenia and bipolar disorder. PLoS One 15, e0232826.

Lee, D.K., Lee, H., Ryu, V., Kim, S.W., Ryu, S., 2022. Different patterns of white matter microstructural alterations between psychotic and non-psychotic bipolar disorder. Plos one 17, e0265671.

Lewandowska, P., Bajada, C.J., Mysak, Y., Domagalik, A., Kossowski, B., Baumbach, C., Kaczmarek-Majer, K., Degórska, A., Skotak, K., Sitnik-Warchulska, K., et al., 2025. The impact of early life exposure to air pollution on the brain: A diffusion MRI study in 10–13-year-old children with and without ADHD diagnosis. Human Brain Mapping 46, e70306.

Li, H., Liang, Z., Zhang, C., Liu, R., Li, J., Zhang, W., Liang, D., Shen, B., Zhang, X., Ge, Y., et al., 2021. SuperDTI: Ultrafast DTI and fiber tractography with deep learning. Magnetic resonance in medicine 86, 3334–3347.

Linke, J.O., Adleman, N.E., Sarlls, J., Ross, A., Perlstein, S., Frank, H.R., Towbin, K.E., Pine, D.S., Leibenluft, E., Brotman, M.A., 2020. White matter microstructure in pediatric bipolar disorder and disruptive mood dysregulation disorder. Journal of the American Academy of Child & Adolescent Psychiatry 59, 1135–1145.

Liu, M., Vemuri, B.C., Deriche, R., 2013. A robust variational approach for simultaneous smoothing and estimation of DTI. NeuroImage 67, 33–41.

Magioncalda, P., Martino, M., Tardito, S., Sterlini, B., Conio, B., Marozzi, V., Adavastro, G., Capobianco, L., Russo, D., Parodi, A., et al., 2018. White matter microstructure alterations correlate with terminally differentiated CD8+ effector T cell depletion in the peripheral blood in mania: combined DTI and immunological investigation in the different phases of bipolar disorder. Brain, Behavior, and Immunity 73, 192–204.

Martin, P., Altbach, M., Bilgin, A., 2025. Conditional generative diffusion deep learning for accelerated diffusion tensor and kurtosis imaging. Magnetic Resonance Imaging 117, 110309.

Matsumoto, M., Nishimura, T., 1998. Mersenne twister: a 623-dimensionally equidistributed uniform pseudo-random number generator. ACM Transactions on Modeling and Computer Simulation (TOMACS) 8, 3–30.

Maximov, I.I., Grinberg, F., Shah, N.J., 2011. Robust tensor estimation in diffusion tensor imaging. Journal of Magnetic Resonance 213, 136–144.

Maximov, I.I., Thönneßen, H., Konrad, K., Amort, L., Neuner, I., Shah, N.J., 2015. Statistical instability of TBSS analysis based on DTI fitting algorithm. Journal of Neuroimaging 25, 883–891.

Minati, L., We?glarz, W.P., 2007. Physical foundations, models, and methods of diffusion magnetic resonance imaging of the brain: A review. Concepts in Magnetic Resonance Part A: An Educational Journal 30, 278–307.

Mori, S., Wakana, S., Van Zijl, P.C., Nagae-Poetscher, L., 2005. MRI atlas of human white matter. Elsevier.

Ni, H., Kavcic, V., Zhu, T., Ekholm, S., Zhong, J., 2006. Effects of number of diffusion gradient directions on derived diffusion tensor imaging indices in human brain. American Journal of Neuroradiology 27, 1776–1781.

Oertel-Knöchel, V., Reinke, B., Alves, G., Jurcoane, A., Wenzler, S., Prvulovic, D., Linden, D., Knöchel, C., 2014. Frontal white matter alterations are associated with executive cognitive function in euthymic bipolar patients. Journal of affective disorders 155, 223–233.

Özarslan, E., Koay, C.G., Shepherd, T.M., Komlosh, M.E., İrfanoğlu, M.O., Pierpaoli, C., Basser, P.J., 2013. Mean apparent propagator (MAP) MRI: a novel diffusion imaging method for mapping tissue microstructure. NeuroImage 78, 16–32.

Park, J., Jung, W., Choi, E.J., Oh, S.H., Jang, J., Shin, D., An, H., Lee, J., 2021. DIFFnet: diffusion parameter mapping network generalized for input diffusion gradient schemes and b-value. IEEE Transactions on Medical Imaging 41, 491–499.

Pasternak, O., Kelly, S., Sydnor, V.J., Shenton, M.E., 2018. Advances in microstructural diffusion neuroimaging for psychiatric disorders. Neuroimage 182, 259–282.

Pieciak, T., París, G., Beck, D., Maximov, I.I., Tristan-Vega, A., de Luis-Garcia, R., Westlye, L.T., Aja-Fernandez, S., 2023. Spherical means-based free-water volume fraction from diffusion MRI increases non-linearly with age in the white matter of the healthy human brain. Neuroimage 279, 120324.

Poldrack, R.A., Congdon, E., Triplett, W., Gorgolewski, K., Karlsgodt, K., Mumford, J., Sabb, F., Freimer, N., London, E., Cannon, T., et al., 2016. A phenome-wide examination of neural and cognitive function. Scientific data 3, 1–12.

Ronneberger, O., Fischer, P., Brox, T., 2015. U-net: Convolutional networks for biomedical image segmentation, in: International Conference on Medical image computing and computer-assisted intervention, Springer. pp. 234–241.

Singh, A., Pandey, H.R., Arya, A., Agarwal, V., Shree, R., Kumar, U., 2023. Altered white matter integrity in euthymic children with bipolar disorder: A tract-based spatial statistical analysis of diffusion tensor imaging. Journal of Affective Disorders 340, 820–827.

Smith, S.M., Jenkinson, M., Johansen-Berg, H., Rueckert, D., Nichols, T.E., Mackay, C.E., Watkins, K.E., Ciccarelli, O., Cader, M.Z., Matthews, P.M., et al., 2006. Tract-based spatial statistics: voxelwise analysis of multi-subject diffusion data. Neuroimage 31, 1487–1505.

Smith, S.M., Jenkinson, M., Woolrich, M.W., Beckmann, C.F., Behrens, T.E., Johansen-Berg, H., Bannister, P.R., De Luca, M., Drobnjak, I., Flitney, D.E., et al., 2004. Advances in functional and structural MR image analysis and implementation as fsl. Neuroimage 23, S208–S219.

Taragola, E., Neyt, S., Yu, B., Abi Akl, M., Vervenne, B., Gong, K., Vandenberghe, S., Vanhove, C., Muller, F.M., 2026. Cross-modality deep learning denoising for low-dose µSPECT: Transfer of PET-trained U-Net and diffusion models. Chemical & Biomedical Imaging 4, 1727–1739.

Tian, Q., Bilgic, B., Fan, Q., Liao, C., Ngamsombat, C., Hu, Y., Witzel, T., Setsompop, K., Polimeni, J.R., Huang, S.Y., 2020. DeepDTI: High-fidelity six-direction diffusion tensor imaging using deep learning. NeuroImage 219, 117017.

Veraart, J., Fieremans, E., Novikov, D.S., 2016a. Diffusion MRI noise mapping using random matrix theory. Magnetic resonance in medicine 76, 1582–1593.

Veraart, J., Novikov, D.S., Christiaens, D., Ades-Aron, B., Sijbers, J., Fieremans, E., 2016b. Denoising of diffusion MRI using random matrix theory. Neuroimage 142, 394–406.

Wang, Z., Bovik, A.C., Sheikh, H.R., Simoncelli, E.P., 2004. Image quality assessment: from error visibility to structural similarity. IEEE transactions on image processing 13, 600–612.

Westin, C.F., Maier, S.E., Mamata, H., Nabavi, A., Jolesz, F.A., Kikinis, R., 2002. Processing and visualization for diffusion tensor MRI. Medical image analysis 6, 93–108.

Westlye, L.T., Walhovd, K.B., Dale, A.M., Bjørnerud, A., Due-Tønnessen, P., Engvig, A., Grydeland, H., Tamnes, C.K., Østby, Y., Fjell, A.M., 2010. Life-span changes of the human brain white matter: diffusion tensor imaging (DTI) and volumetry. Cerebral cortex 20, 2055–2068.

Winkler, A.M., Ridgway, G.R., Webster, M.A., Smith, S.M., Nichols, T.E., 2014. Permutation inference for the general linear model. Neuroimage 92, 381–397.

Xin, B., Hu, Y., Zheng, Y., Liao, H., 2020. Multi-modality generative adversarial networks with tumor consistency loss for brain mr image synthesis, in: 2020 IEEE 17th international symposium on biomedical imaging (ISBI), IEEE. pp. 1803–1807.

Yang, J., Jiang, X., Wei, S., Deng, X., Zhu, Y., Chang, M., Yin, Z., Geng, H., Tang, Y., Dai, X., 2021. White matter tracts in bipolar disorder patients: a comparative study based on diffusion kurtosis and tensor imaging. Journal of Affective Disorders 292, 45–55.

Yi, X., Walia, E., Babyn, P., 2019. Generative adversarial network in medical imaging: A review. Medical image analysis 58, 101552.

Zhang, H., Schneider, T., Wheeler-Kingshott, C.A., Alexander, D.C., 2012. NODDI: practical in vivo neurite orientation dispersion and density imaging of the human brain. Neuroimage 61, 1000–1016.

Zhang, Y., Brady, M., Smith, S., 2002. Segmentation of brain MR images through a hidden markov random field model and the expectation-maximization algorithm. IEEE transactions on medical imaging 20, 45–57.

Zoghby, M., Erickson, B., Conte, G., 2024. Generative adversarial networks for brain MRI synthesis: impact of training set size on clinical application. Journal of imaging informatics in medicine 37, 1228–1238.

