## Supplementary material for "Clinical relevance of diffusion tensor imaging-based parameters estimated with deep learning: the effect of model retraining": Supplementary_materials_biorxiv.pdf

### 1. Technical details

#### *1.1. Data postprocessing*

In what follows, we present four different resampling and subsampling schemes used in our evaluation framework. This section provides technical details related to the resampling and subsampling procedures, which are then applied in the three different configurations presented in section 2.4 of the main manuscript.

Data resampling (Configuration 1): To simulate reduced gradient direction schemes while maintaining a common 64-direction reference, we resampled the original dataset onto a new 64-direction scheme following the electrostatic repulsion algorithm (Jones et al., 1999; Caruyer et al., 2013). This method generates a gradient sampling scheme such that any subset of the first $k$ directions remains approximately uniformly distributed on the sphere. To resample the data, the original DW MR signals were projected onto a real-valued spherical harmonic basis (Descoteaux et al., 2007) with a maximum order of $L_{\max} = 6$ and then resampled onto the new 64-direction gradient scheme. This resampled dataset serves as the reference for this configuration.

Data subsampling (Configuration 1): Reduced acquisition schemes were then simulated by subsampling the resampled 64-direction dataset. This involved selecting the first $k$ gradient directions and their corresponding DW volumes, where $k = \{48, 32, 24, 16, 6, 3\}$.

Data resampling (Configuration 2): To simulate reduced gradient direction schemes, the original dataset was independently resampled for each reduced value of $k$, where $k = \{48, 32, 24, 16, 6, 3\}$. The original DW MR signals were projected onto a real-valued spherical harmonic basis with a maximum order of $L_{\max} = 6$ and then resampled onto each of the six new gradient schemes. The original 64-direction dataset was retained without resampling and served as the reference for this configuration.

Data resampling (Configuration 3): To simulate reduced gradient direction schemes, the original dataset was independently resampled onto new schemes with $k$ gradient directions, where $k = \{64, 48, 32, 24, 16, 6\}$. The original DW MR signals were projected onto a real-valued spherical harmonic basis with a maximum order of $L_{\max} = 6$ and then resampled onto each of the six new gradient schemes. Additionally, each gradient scheme underwent 100 random rigid rotations, and the resampling procedure was repeated for each rotated scheme. Consequently, for each value of $k$, 100 independently rotated gradient schemes and their corresponding resampled datasets were generated. The unrotated 64-direction resampled dataset served as the reference, as in Configuration 1.

#### *1.2. TBSS-based evaluation metrics*

Figure S1 illustrates the true positive rate (TPR) and false positive rate (FPR) for the body of the corpus callosum (BCC), using two representative DL model runs with markedly different FPR values. TPR quantifies the proportion of voxels identified as statistically significant in the reference analysis that are also significant in the DL-based analysis. FPR quantifies the number of additional significant voxels identified by the DL-based analysis, relative to the number of significant voxels in the reference analysis. Because FPR is normalized by the number of reference-significant voxels rather than the total number of voxels analyzed, it is not bounded by 100%. Values above 100% indicate that the number of additional significant voxels exceeds the number of significant voxels in the reference analysis.

*Corresponding author

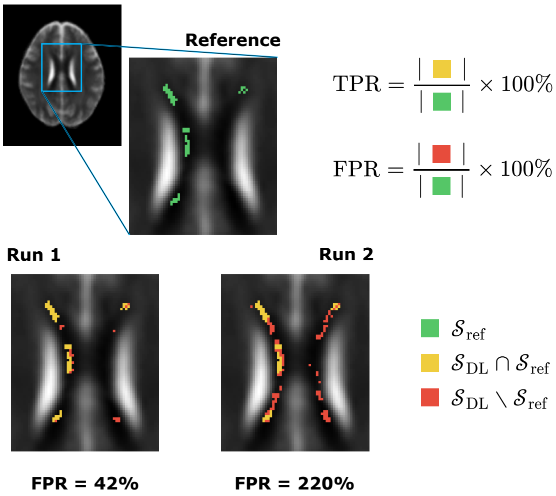


Figure S1: Visual illustration of the tract-based spatial statistics (TBSS) evaluation metrics, true positive rate (TPR) and false positive rate (FPR), for two representative deep learning model runs. The blue box on the whole-brain slice indicates the analyzed region of interest. Voxels exhibiting statistically significant group differences in the reference analysis ($\mathcal{S}_{\text{ref}}$) are shown in green. Yellow voxels correspond to true positives, defined as $\mathcal{S}_{\text{DL}} \cap \mathcal{S}_{\text{ref}}$, whereas red voxels correspond to false positives, defined as $\mathcal{S}_{\text{DL}} \setminus \mathcal{S}_{\text{ref}}$. Accordingly, TPR is calculated as the number of yellow voxels divided by the number of green voxels, and FPR as the number of red voxels divided by the number of green voxels, expressed as percentages.

### 2. Bipolar disorder: radial diffusivity

#### *2.1. Comparison of between-group differences between original and resampled dataset*

To evaluate whether the resampling procedure influenced the identification of regions with significant between-group differences, we repeated the reference Tract-Based Spatial Statistics (TBSS) analysis described in the main manuscript using the original, non-resampled dataset. Table S1 summarizes the statistically significant regions of interest (ROIs) identified in the original dataset. The same ROIs were found to be statistically significant in both the original and resampled datasets, with the group difference consistently indicating that the bipolar disorder (BD) group had higher radial diffusivity (RD) values than the healthy controls (HC) group (BD > HC). These findings suggest that the resampling procedure did not change the set of ROIs identified as showing significant between-group differences, nor did it affect the direction of these effects. As a result, the same ROIs can be utilized for the subsequent ROI-based analyses in both Configuration 1, which is based on the resampled dataset, and Configuration 2, which relies on the original dataset.

Table S1: White matter regions identified by tract-based spatial statistics (TBSS) analysis of **radial diffusivity (RD)** in patients with bipolar disorder (BD) compared with healthy controls (HC) for **Configuration 2**. The table includes only regions that show a significant increase in RD (BD > HC) relative to HC after the threshold-free cluster enhancement (TFCE) correction ($p < 0.05$). Significant voxels are reported as $n/N$ (%), where $n$ is the number of voxels showing a statistically significant difference, and $N$ is the total number of voxels within the corresponding white matter region. Peak T-stat represents the maximum $t$-statistic within the significant portion of the region. Coordinates (X, Y, Z) indicate the location of the peak voxel in MNI space. Corrected $p$-value refers to the TFCE family-wise error-corrected $p$-value at the peak voxel.

| Parameter | ROIs | Significant voxels | Peak T-stat | Coordinates (X,Y,Z) | Corrected *p*-value |
|---|---|---|---|---|---|
| **RD (BD > HC)** | GCC | 168/1788 (9.40%) | 3.613 | 16, 22, 23 | 0.0174 |
| | BCC | 583/3259 (17.89%) | 3.821 | 17, 0, 35 | 0.0176 |
| | SCC | 397/2397 (16.56%) | 4.618 | -21, -49, 22 | 0.0498 |
| | ALIC R | 467/803 (58.16%) | 5.200 | 21, 6, 13 | 0.0100 |
| | ACR R | 717/1649 (43.48%) | 3.861 | 25, 27, 4 | 0.0172 |
| | ACR L | 318/1781 (17.86%) | 4.592 | -19, 30, 17 | 0.0444 |
| | SCR R | 474/1545 (30.68%) | 5.044 | 18, 15, 34 | 0.0166 |
| | PCR R | 276/737 (37.45%) | 3.661 | 28, -34, 20 | 0.0440 |
| | PTR R | 381/1146 (33.25%) | 4.007 | 33, -62, 7 | 0.0352 |

#### 2.2. *Configuration 1 (Resampled reference)*

The resampled 64-direction dataset serves as the reference for the DL-based DTI. The reference RD maps were computed from this dataset using the nonlinear least-squares (NLLS) fitting procedure, and then used as pseudo-ground truth for the DL network training procedure. The reduced inputs with $k = \{48, 32, 24, 16, 6, 3\}$ gradient directions were prepared by selecting the first $k$ volumes from the resampled dataset.

Figures S2, S3, S4, and S5 present results that are similar to those shown in the main manuscript for additional ROIs evaluated under Configuration 1. A consistent relationship between image-level agreement and downstream statistical results was observed across these regions. Notably, different runs exhibited considerable variability in true positive rate (TPR) and false positive rate (FPR), despite showing relatively stable mean squared error (MSE) and mean structural similarity index measure (MSSIM) values. As the number of gradient directions decreased, the distributions generally shifted towards higher TPR and FPR values, with FPR becoming more concentrated at these higher values. The differences in the FPR ranges between ROIs are expected, as FPR is normalized by the number of significant voxels in the corresponding reference analysis. The ROI-based statistical analyses indicated corresponding shifts in the distributions of $t$-statistics and Cohen's $d$, with the estimated values typically being higher than those obtained from the reference analysis as the number of gradient directions decreased. Similarly, Bland-Altman analyses showed an increasing disparity in the estimated RD values compared to the reference as the number of gradient directions was reduced. Overall, these findings are consistent with those presented in the main manuscript and suggest that the variability observed in downstream statistical results is not limited to the ROIs chosen for the primary analysis.

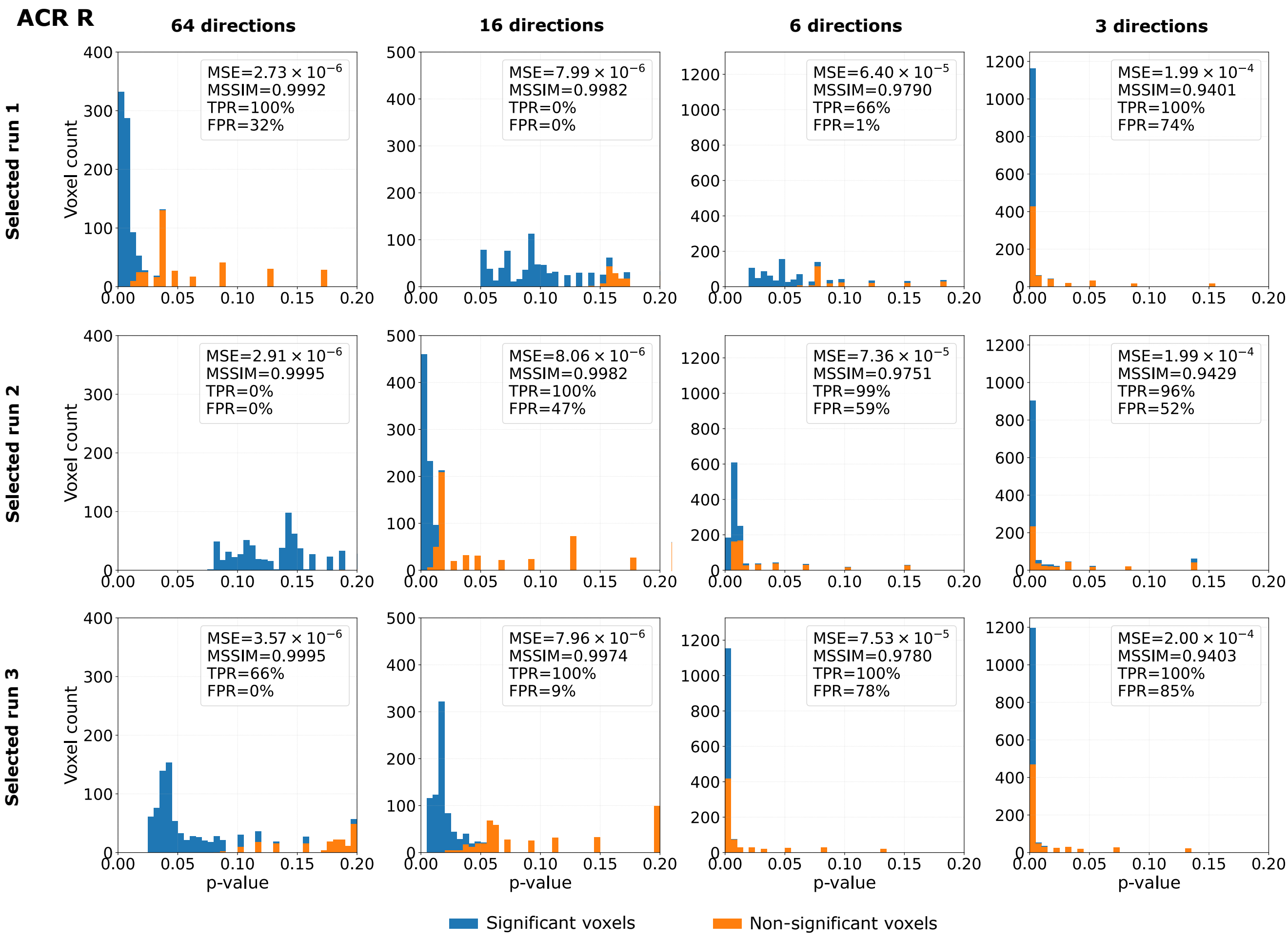


Figure S2: Histograms of voxel-wise $p$-values obtained from the tract-based spatial statistics (TBSS) analysis for the **RD parameter** estimated using a deep learning model following **Configuration 1** over the skeleton of the right anterior corona radiata (ACR R) and the genu of the corpus callosum (GCC). The figure presents three selected runs over 100 runs characterized with a comparable mean squared error (MSE) and mean structural similarity index measure (MSSIM), while variable true positive and false positive rates (TPR, FPR). For each run, four diffusion gradient distributions of input data are considered: 64, 16, 6, and 3. Bins in blue refer to the voxels, which were statistically significant in the TBSS analysis based on the standard NLLS-derived DTI (no deep learning applied), while orange bins show the voxels statistically non-significant in the TBSS analysis based on NLLS-derived DTI.

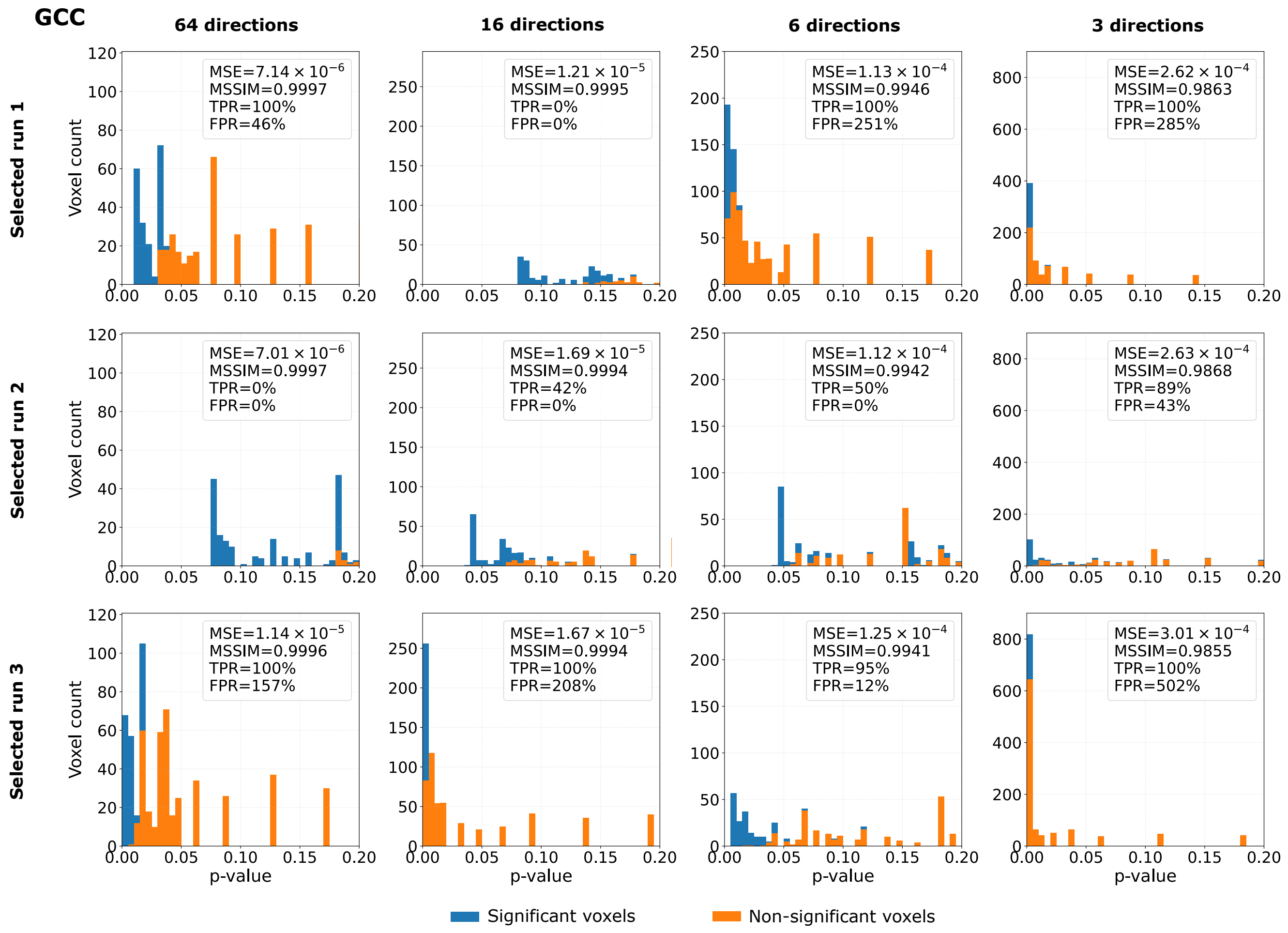


Figure S2 (continued)

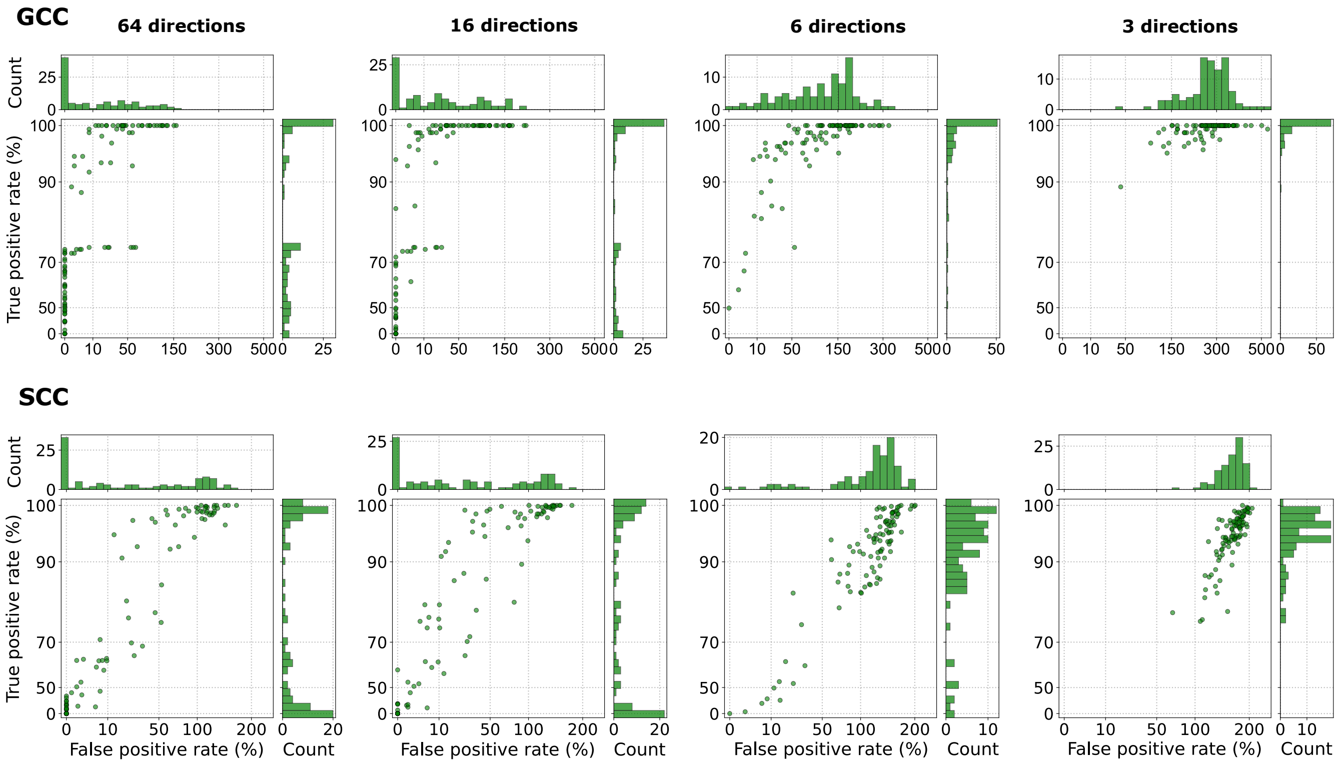


Figure S3: Relationship between the true positive rate and the false positive rate, obtained from the tract-based spatial statistics (TBSS) analysis for **RD parameter**, in the genu (GCC) and the splenium (SCC) of the corpus callosum using **Configuration 1**, with four different numbers of input data gradient directions (64, 16, 6, and 3). Each point refers to a single run. In total, 100 runs were considered. The FPRs greater than 100% result from computing the values relative to the number of significant voxels identified in the reference analysis.

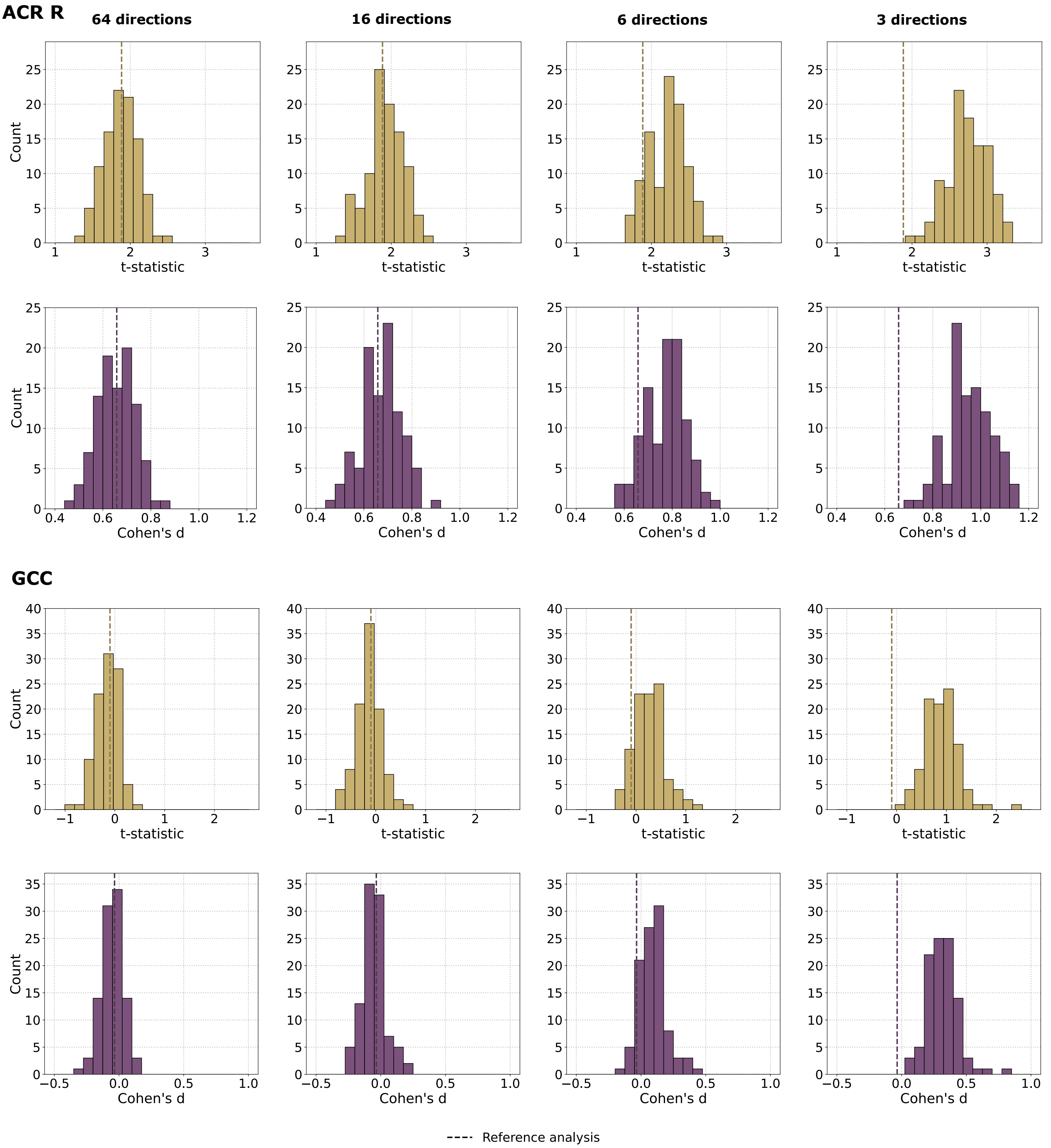


Figure S4: Distributions of $t$-statistics (top row) and Cohen's $d$ values (bottom row) in the right anterior corona radiata (ACR R) and the genu of the corpus callosum (GCC) for the **RD parameter** estimated using a deep learning model, with four different numbers of input data gradient directions (64, 16, 6, and 3) (**Configuration 1**). These statistics were calculated by comparing the mean RD values between patients with bipolar disorder (BD) and healthy controls (HC). Each histogram summarizes the results obtained from 100 runs. The dashed vertical line indicates the corresponding value from the reference analysis using the NLLS estimation.

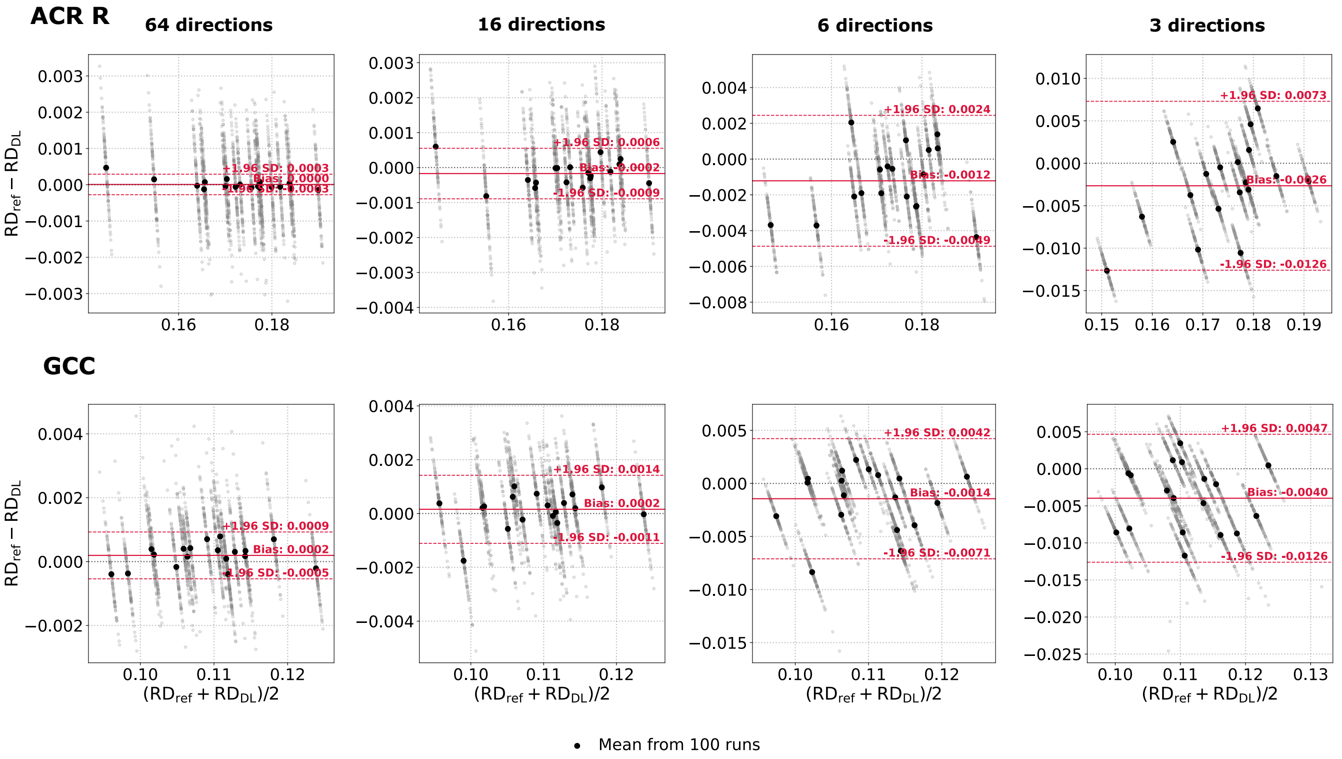


Figure S5: Bland-Altman plots of the **RD parameter** computed from the right anterior corona radiata (ACR R) and genu of the corpus callosum (GCC) across all BD subjects. The reference values ($RD_{ref}$) were obtained using the NLLS estimation from a resampled 64-direction scheme, whereas the predicted $RD_{DL}$ values were generated using the deep learning model (**Configuration 1**) considering four different numbers of input data gradient directions (64, 16, 6, and 3). Gray points represent individual model runs (mean values over the ROI). A single black point corresponds to the average across 100 runs for a single BD patient. Solid red lines indicate the bias, while dashed red lines denote 95% limits of agreement (±1.96 standard deviations).

#### 2.3. *Configuration 2 (Original reference)*

RD maps used as pseudo-ground truth for DL training were computed from the original 64-direction acquisition using the NLLS algorithm. Reduced datasets were generated independently by resampling the original data to new gradient tables with $k = \{48, 32, 24, 16, 6, 3\}$ gradient directions.

The analyses conducted for Configuration 1 were repeated using Configuration 2. The results from this analysis are illustrated in Figures S6, S7, S8, S9, S10, S11, and Table S2. Overall, the findings from Configuration 2 mirrored the patterns observed in Configuration 1. Across runs, TPR and FPR varied substantially, despite relatively consistent image-level similarity metrics. This variability became more pronounced as the number of gradient directions decreased. Additionally, the ROI-based statistical analyses reflected corresponding variability in the $t$-statistics and Cohen's $d$. The Bland-Altman analyses also indicated an increase in variability of the estimated RD values with more significantly reduced acquisition schemes. The distributions of the image-level and TBSS metrics were generally similar for both configurations. However, we observed differences in the magnitude of individual metrics for certain acquisition schemes and ROIs. Importantly, these differences did not alter the overall relationship between image-level agreement and downstream statistical results. Therefore, the results from Configuration 2 provide further evidence that the variability in downstream statistical outcomes is not specific to the procedures used to generate the reduced acquisition schemes or the definition of reference data.

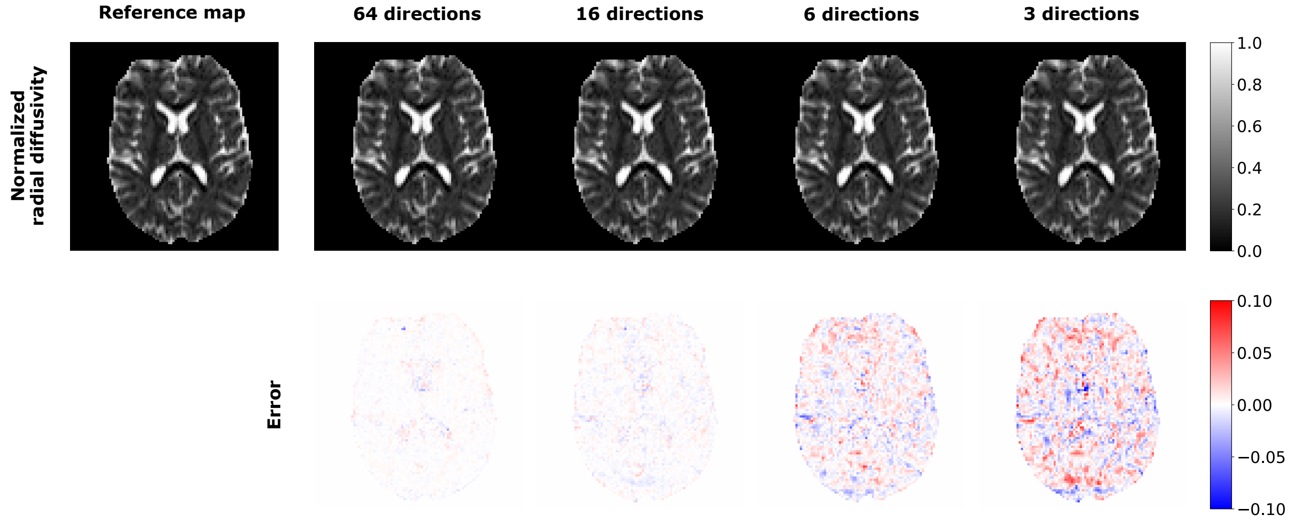


Figure S6: Visual inspection of **normalized radial diffusivity** estimated using single runs of the deep learning model and the corresponding voxel-wise differences between DL-based maps and the reference map derived from the original 64-direction scheme using the NLLS estimation (**Configuration 2**). Four input diffusion gradient schemes are considered for DL-based estimation: 64, 16, 6, and 3 directions.

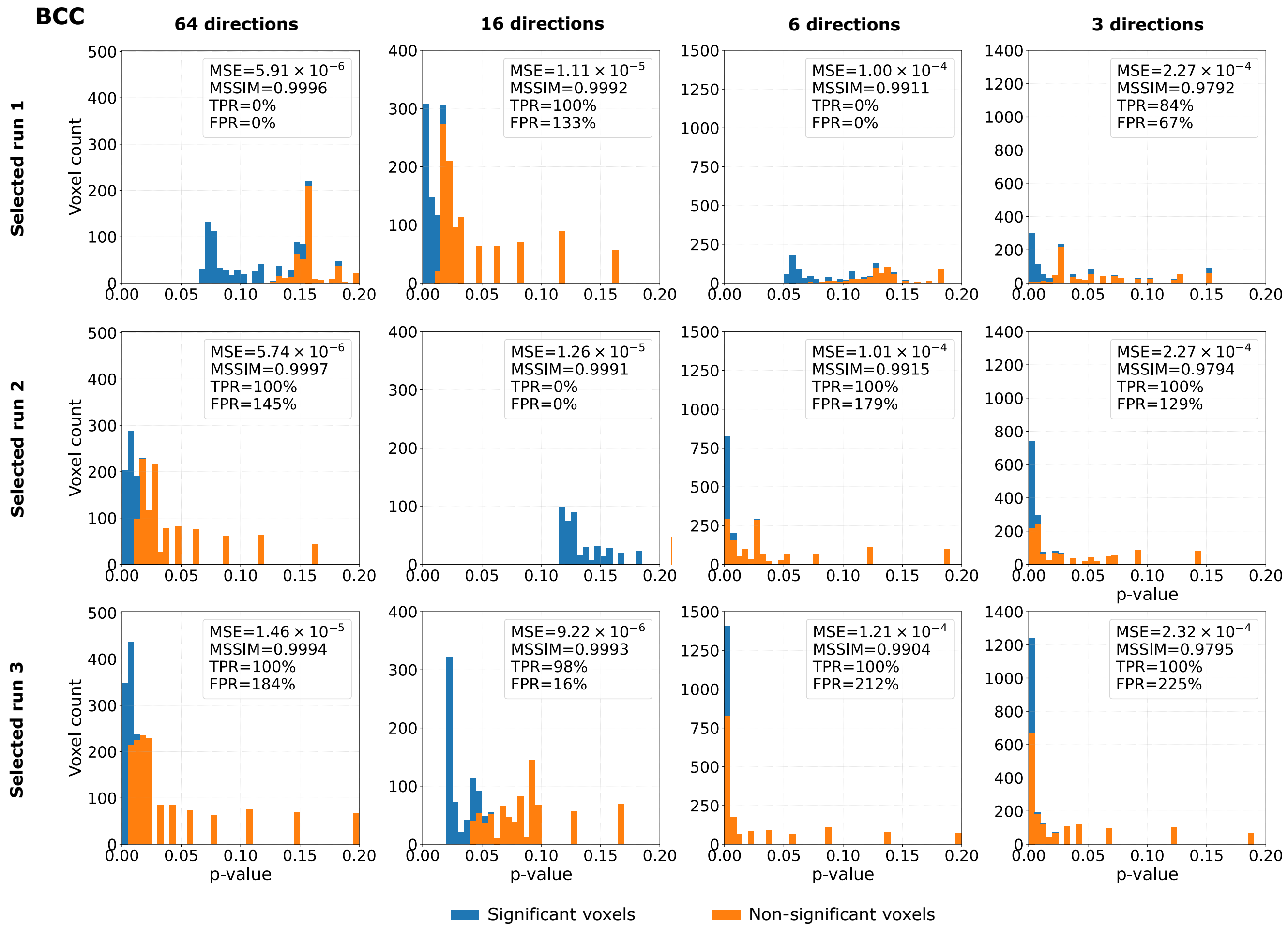


Figure S7: Histograms of voxel-wise $p$-values obtained from the tract-based spatial statistics (TBSS) analysis for the **RD parameter** estimated using a deep learning model following **Configuration 2** over the skeleton of the body (BCC) and the genu (GCC) of the corpus callosum, and the right anterior corona radiata (ACR R). The figure presents three selected runs over 100 runs characterized with a comparable mean squared error (MSE) and mean structural similarity index measure (MSSIM), while variable true positive and false positive rates (TPR, FPR). For each run, four diffusion gradient distributions of input data are considered: 64, 16, 6, and 3. Bins in blue refer to the voxels, which were statistically significant in the TBSS analysis based on the standard NLLS-derived DTI (no deep learning applied), while orange bins show the voxels statistically non-significant in the TBSS analysis based on NLLS-derived DTI.

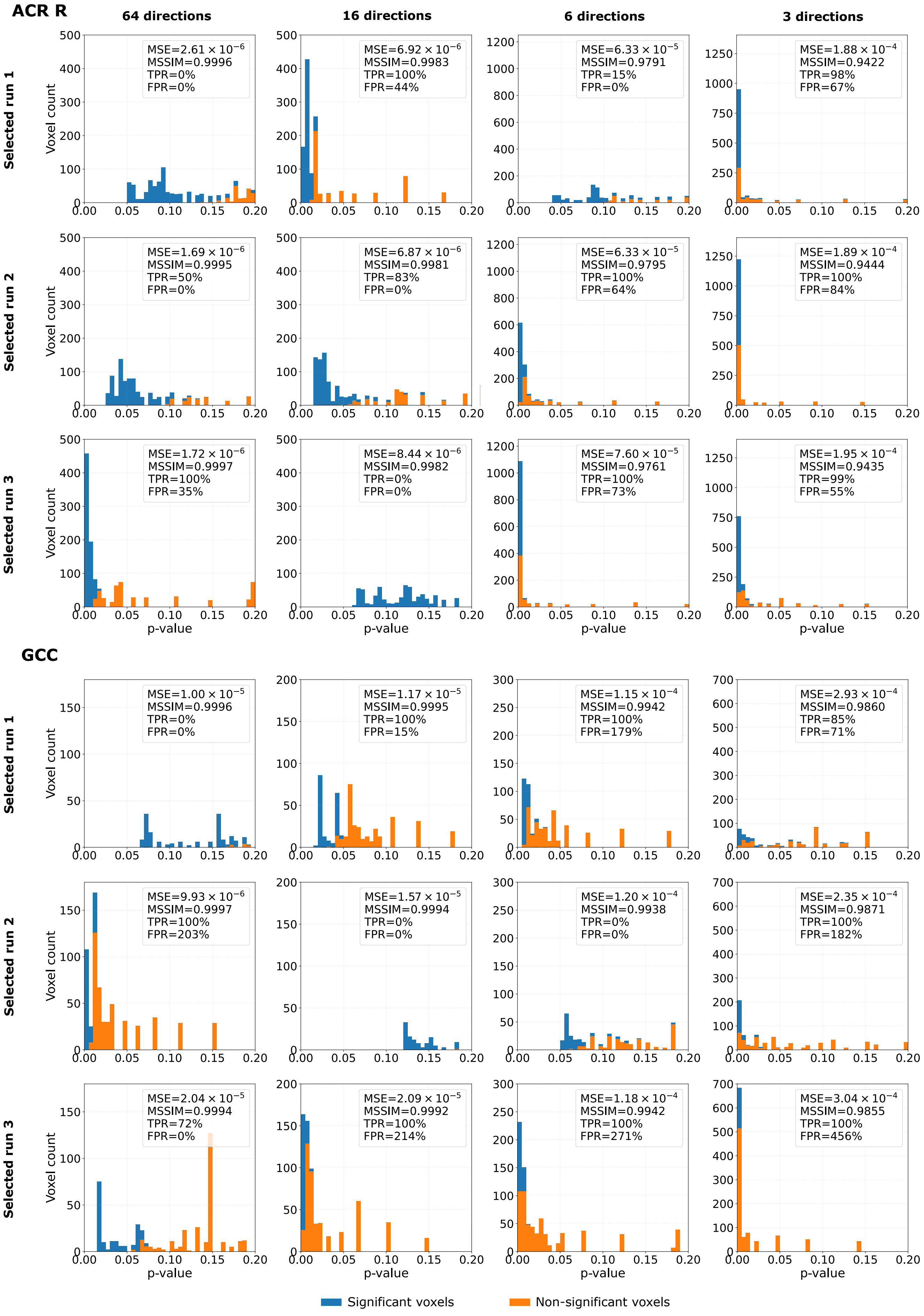


Figure S7 (continued)

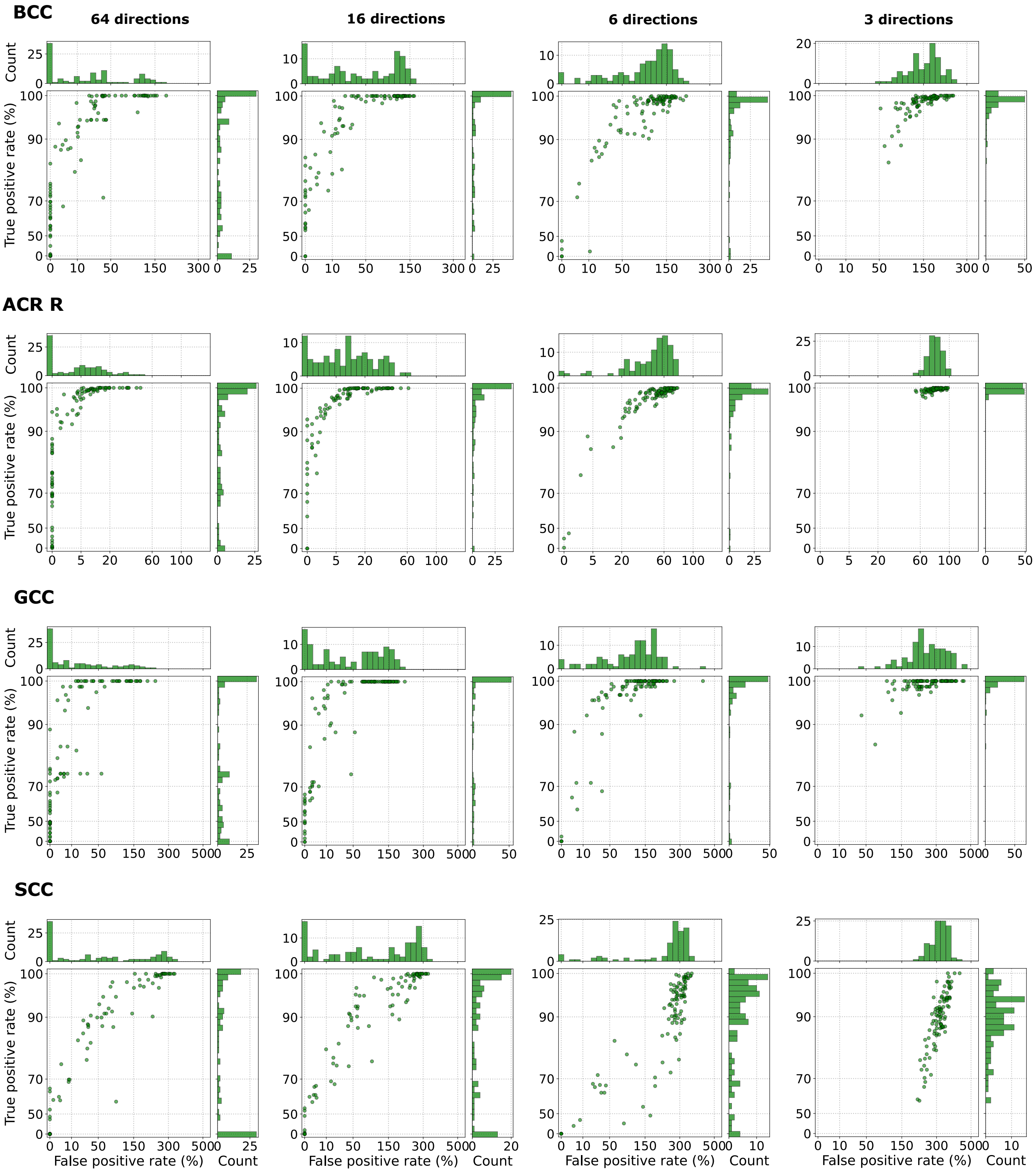


Figure S8: Relationship between the true positive rate and the false positive rate, obtained from the tract-based spatial statistics (TBSS) analysis for **RD parameter**, in the body (BCC), genu (GCC) and the splenium (SCC) of the corpus callosum, and the right anterior corona radiata (ACR R) using **Configuration 2**, with four different numbers of input data gradient directions (64, 16, 6, and 3). Each point refers to a single run. In total, 100 runs were considered. The FPRs greater than 100% result from computing the values relative to the number of significant voxels identified in the reference analysis.

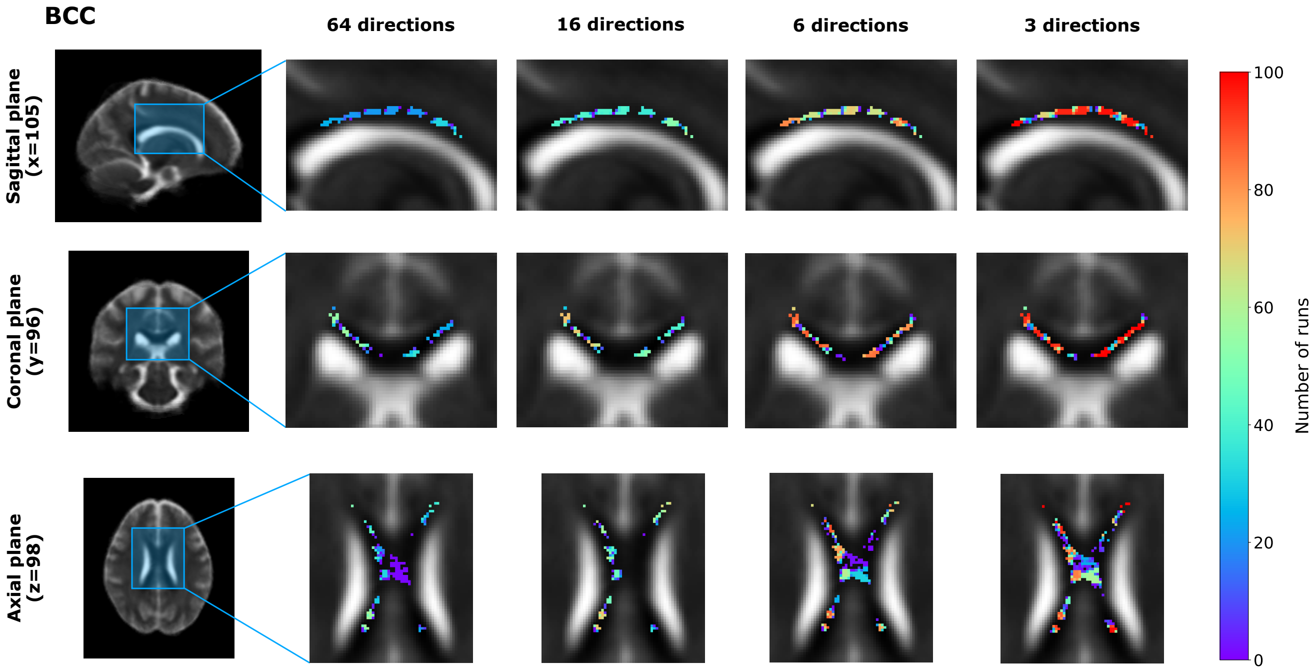


Figure S9: Visual representation of false positive voxels, obtained from the tract-based spatial statistics (TBSS) analysis for **RD parameter**, in the body of the corpus callosum (BCC) shown in the sagittal, coronal, and axial planes for four input gradient directions: 64, 16, 6, and 3 (**Configuration 2**). False positive voxels correspond to locations identified as exhibiting statistically significant group differences in the estimated RD maps but not in the reference analysis. The color indicates the number of runs in which a given voxel was classified as a false positive.

Table S2: Decision agreement between DL-based and reference ROI analyses across four different numbers of input data gradient directions (64, 16, 6, and 3) (**Configuration 2**). The reference $p$-value and significance decision are reported for each ROI, while decision agreement indicates the percentage of runs out of 100 that reproduced the statistical significance of the reference analysis ($p < 0.05$ or $p \geq 0.05$).

| **ROIs** | **Reference $p$-value** | **Reference decision** | **Decision agreement** | | | |
|---|---|---|---|---|---|---|
| | | | **64 directions** | **16 directions** | **6 directions** | **3 directions** |
| GCC | 0.9098 | Non-sig ($p \geq 0.05$) | 100% | 99% | 100% | 100% |
| BCC | 0.1956 | Non-sig ($p \geq 0.05$) | 99% | 99% | 97% | 86% |
| SCC | 0.0102 | Sig ($p < 0.05$) | 99% | 100% | 99% | 98% |
| ALIC R | 0.0214 | Sig ($p < 0.05$) | 94% | 87% | 91% | 100% |
| ACR R | 0.0744 | Non-sig ($p \geq 0.05$) | 81% | 66% | 31% | 0% |
| ACR L | 0.0530 | Non-sig ($p \geq 0.05$) | 44% | 37% | 16% | 0% |
| SCR R | 0.0808 | Non-sig ($p \geq 0.05$) | 64% | 54% | 54% | 22% |
| PCR R | 0.0964 | Non-sig ($p \geq 0.05$) | 88% | 71% | 73% | 60% |
| PTR R | 0.0014 | Sig ($p < 0.05$) | 100% | 100% | 100% | 100% |

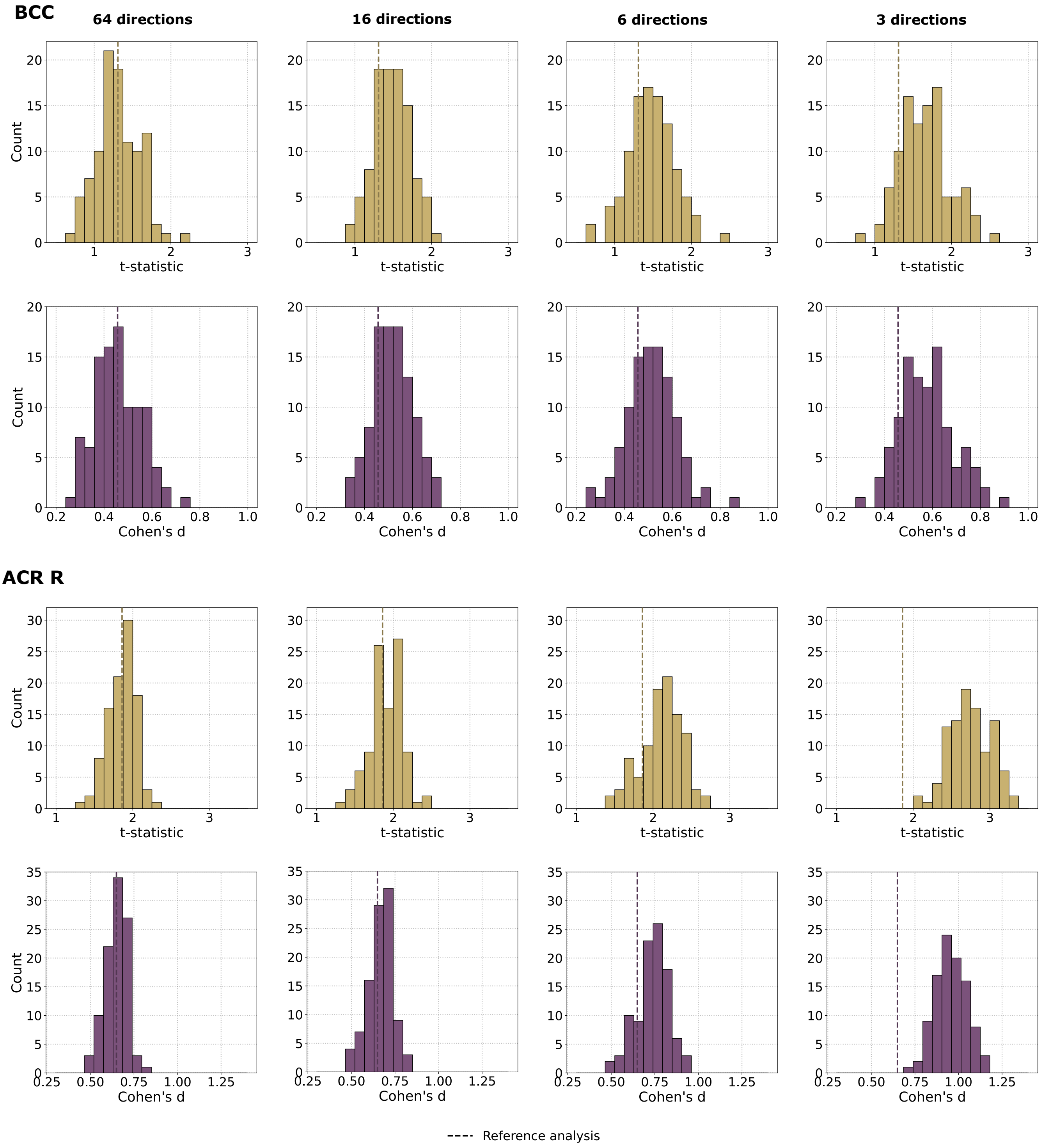


Figure S10: Distributions of $t$-statistics (top row) and Cohen's $d$ values (bottom row) in the body (BCC) and the genu (GCC) of the corpus callosum, and the right anterior corona radiata (ACR R) for the **RD parameter** estimated using a deep learning model, with four different numbers of input data gradient directions (64, 16, 6, and 3) (**Configuration 2**). These statistics were calculated by comparing the mean RD values between patients with bipolar disorder (BD) and healthy controls (HC). Each histogram summarizes the results obtained from 100 runs. The dashed vertical line indicates the corresponding value from the reference analysis using the NLLS estimation.

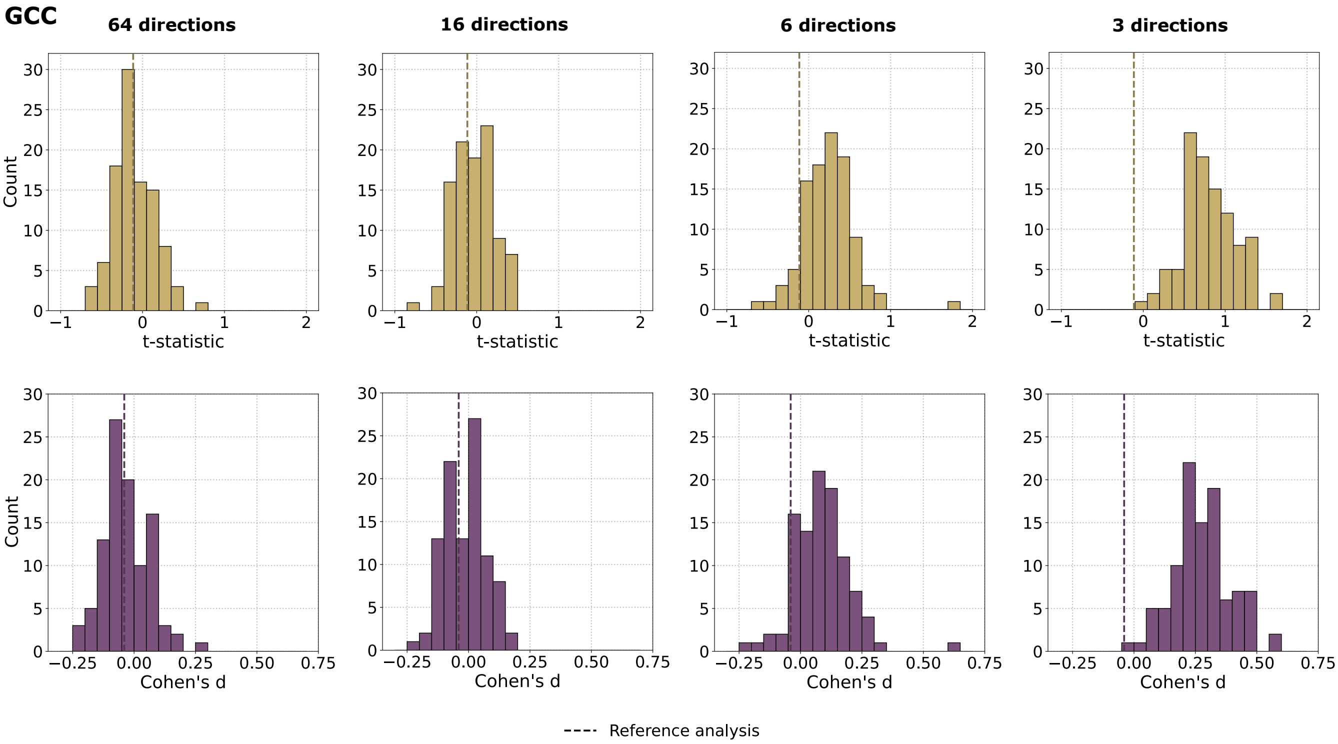


Figure S10 (continued)

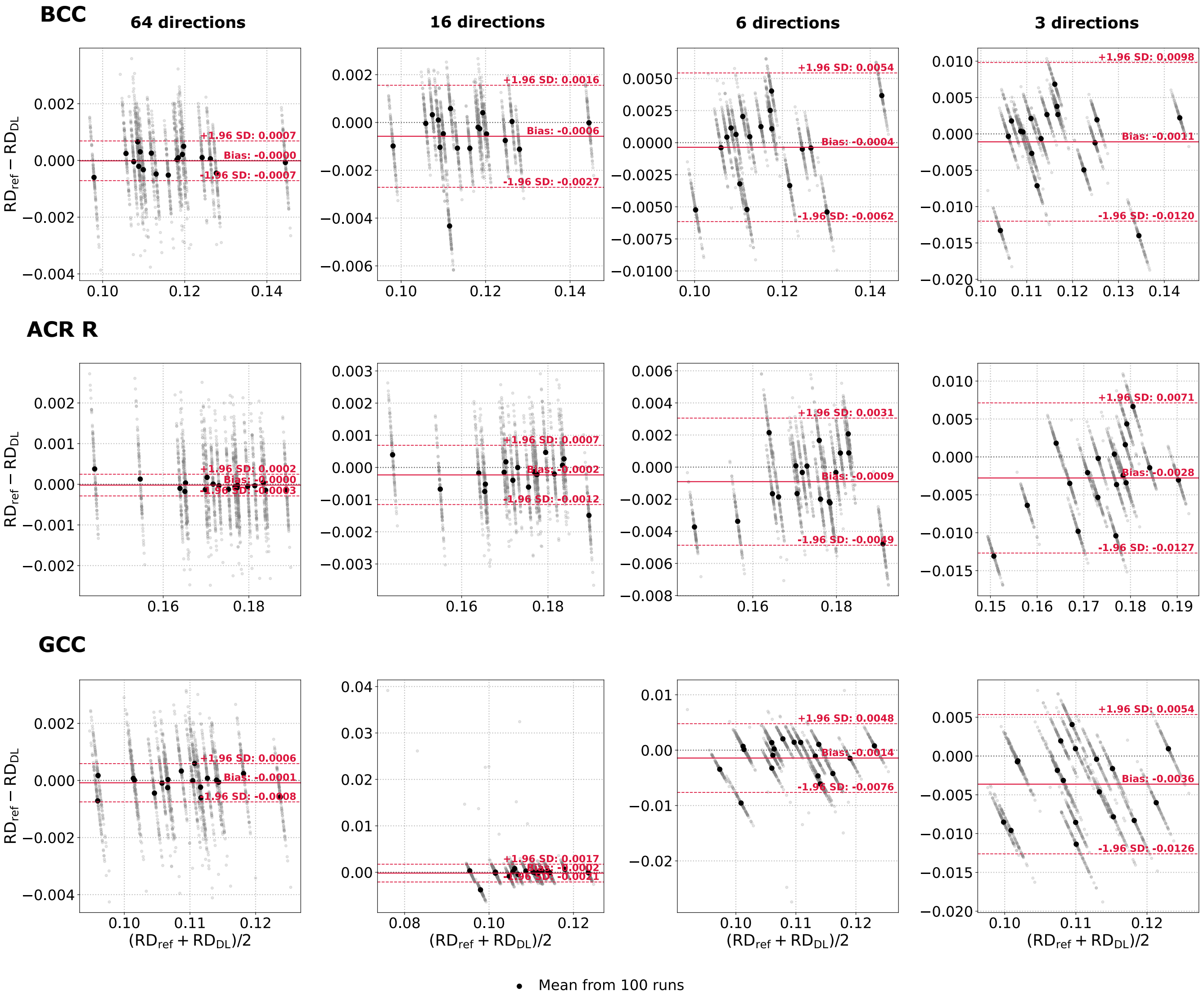


Figure S11: Bland-Altman plots of the **RD parameter** computed from the body (BCC) and genu of the corpus callosum (GCC), and right anterior corona radiata (ACR R) across all BD subjects. The reference values ($RD_{ref}$) were obtained using the NLLS estimation from an original 64-direction scheme, whereas the predicted $RD_{DL}$ values were generated using the deep learning model (**Configuration 2**) considering four different numbers of input data gradient directions (64, 16, 6, and 3). Gray points represent individual model runs (mean values over the ROI). A single black point corresponds to the average across 100 runs for a single BD patient. Solid red lines indicate the bias, while dashed red lines denote 95% limits of agreement (±1.96 standard deviations).

### *2.4. Configuration 3 (Rotated gradients)*

To assess the sensitivity of RD parameter estimated with the NLLS algorithm to gradient distribution, a rotation-based evaluation was performed. Specifically, this analysis focused exclusively on gradient distribution schemes that included $k = \{64, 48, 32, 24, 16, 6\}$ directions. The 3-direction scheme was excluded as the NLLS algorithm requires at least six diffusion gradient directions to estimate the diffusion tensor. For each gradient setup, 100 rigid rotations were generated. Rotation matrices were sampled from the special orthogonal group SO(3) using a Haar-uniform distribution. To ensure the reproducibility, all rotations were generated using a fixed random seed, resulting in a deterministic set of 100 rotation matrices shared across all experiments. Rotated gradient tables were then used to recompute RD maps using the NLLS fitting procedure (Figure S12).

The analyses conducted for Configurations 1 and 2 were also performed for Configuration 3. The relevant results are presented in Figures S13, S14, S15, S16, S17, S18, and Table S3. Overall, these results showed a different pattern compared to what was observed with the DL-based estimation. For larger numbers of gradient directions, rotating the gradient scheme had a relatively minor effect on the NLLS estimates. The resulting RD maps remained highly similar to the reference maps based on both MSE and MSSIM. Additionally, the corresponding TBSS results demonstrated low FPR and high TPR across the analyzed ROIs. The distributions of TPR and FPR exhibited significantly less variability across rotations compared to the DL runs. Similarly, the ROI-based statistical measures, including *t*-statistics, Cohen's *d*, and decision agreement with the reference analysis, remained relatively stable across rotations. The impact of the gradient distribution became more pronounced as the number of directions decreased. At 64 and 16 directions, both the image-level and statistical results remained stable across rotations. However, with only 6 directions, the NLLS estimates displayed an increase in image-level error, as evidenced by a decrease in MSSIM and an increase in MSE. This deterioration was accompanied by a substantial reduction in TPR, indicating that fewer statistically significant voxels identified in the reference analysis were reproduced. Conversely, FPR remained low for most rotations, with higher values observed for only a few rotations. The combination of reduced TPR and consistently low FPR at 6 directions, along with Bland-Altman analyses, indicates a tendency to underestimate RD compared to the reference. This behavior differs from the DL-based estimates, for which the Bland-Altman analyses revealed an increasingly positive shift in RD compared to the reference as the number of gradient directions decreased, alongside an increase in FPR. Overall, Configuration 3 demonstrates that NLLS estimation is relatively insensitive to the specific distribution of gradient directions when a sufficient number of directions is available, whereas substantial instability occurs under a significantly reduced acquisition scheme.

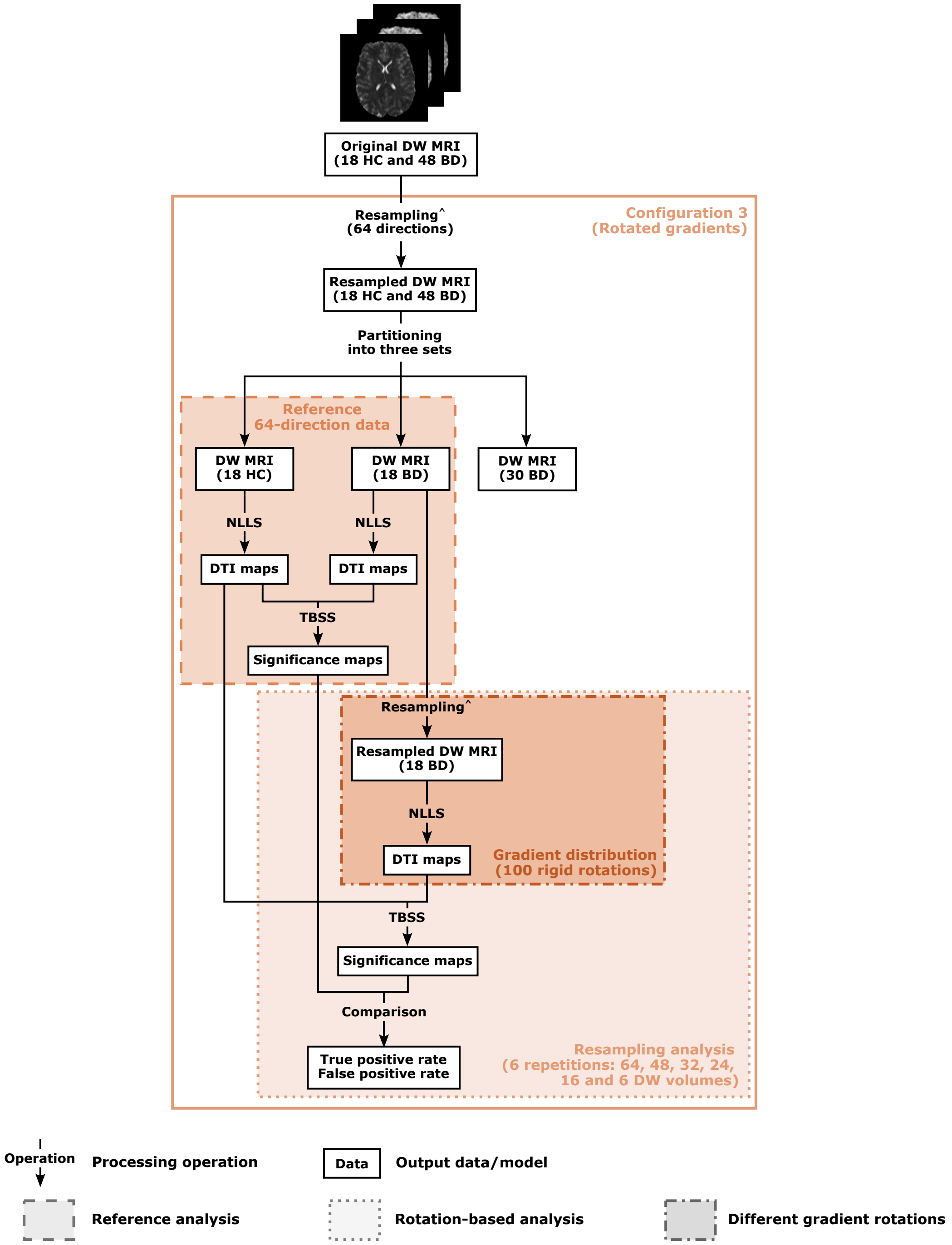


Figure S12: Overview of the **Configuration 3** (Rotated gradients). This analysis evaluated the sensitivity of DTI parameter estimation to the distribution of gradient directions using the conventional non-linear least squares (NLLS) method. The analysis focused exclusively on gradient schemes containing $k = 64, 48, 32, 24, 16, 6$ directions. The 3-direction scheme was excluded because at least six diffusion gradient directions are required for NLLS estimation of the diffusion tensor. The analysis consists of three main stages: reference analysis, rotation-based analysis, and repeated gradient rotations. The reference analysis is performed once using the unrotated 64-direction data. During this stage, the DTI maps are estimated using the conventional NLLS method, and the resulting statistical significance maps serve as the reference. The rotation-based analysis is performed separately for each of the six gradient-direction schemes. For each scheme, 100 rigid rotations of the gradient directions are generated. The rotated gradient tables are then used to recompute the DTI maps using the NLLS fitting procedure. For each rotation, statistical significance maps are obtained and compared with the reference significance maps. The agreement between the rotation-based and reference significance maps is evaluated using the true positive and false positive rates. The symbols used for resampling refer to the corresponding procedure described in Section 1. Resampling^ denotes *Data resampling (Configuration 3).*

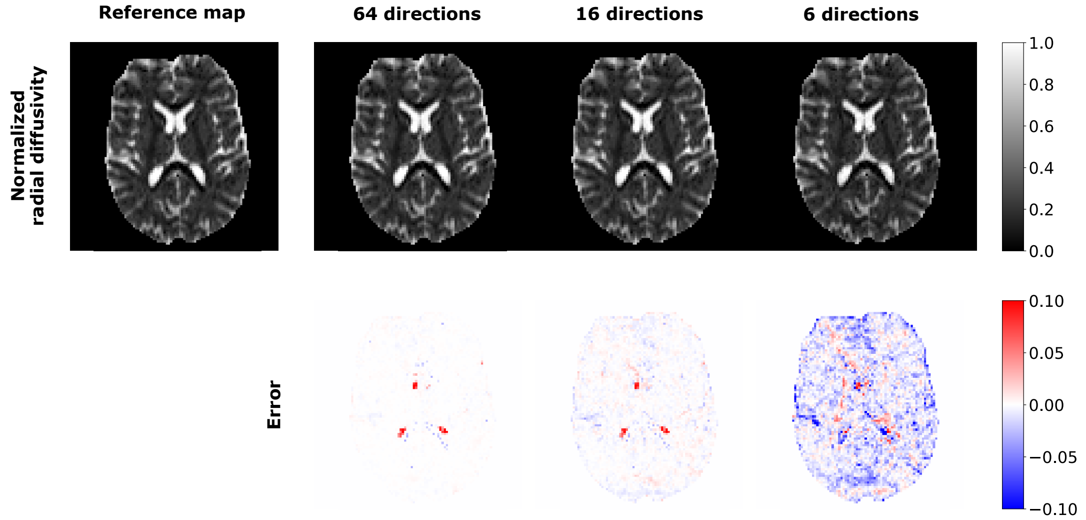


Figure S13: Visual inspection of **normalized radial diffusivity** estimated using NLLS method for a single rotation of the diffusion gradient directions and the corresponding voxel-wise differences between DL-based maps and the reference map derived from the resampled 64-direction scheme using the NLLS estimation (**Configuration 3**). Three input diffusion gradient schemes are considered: 64, 16, and 6 directions.

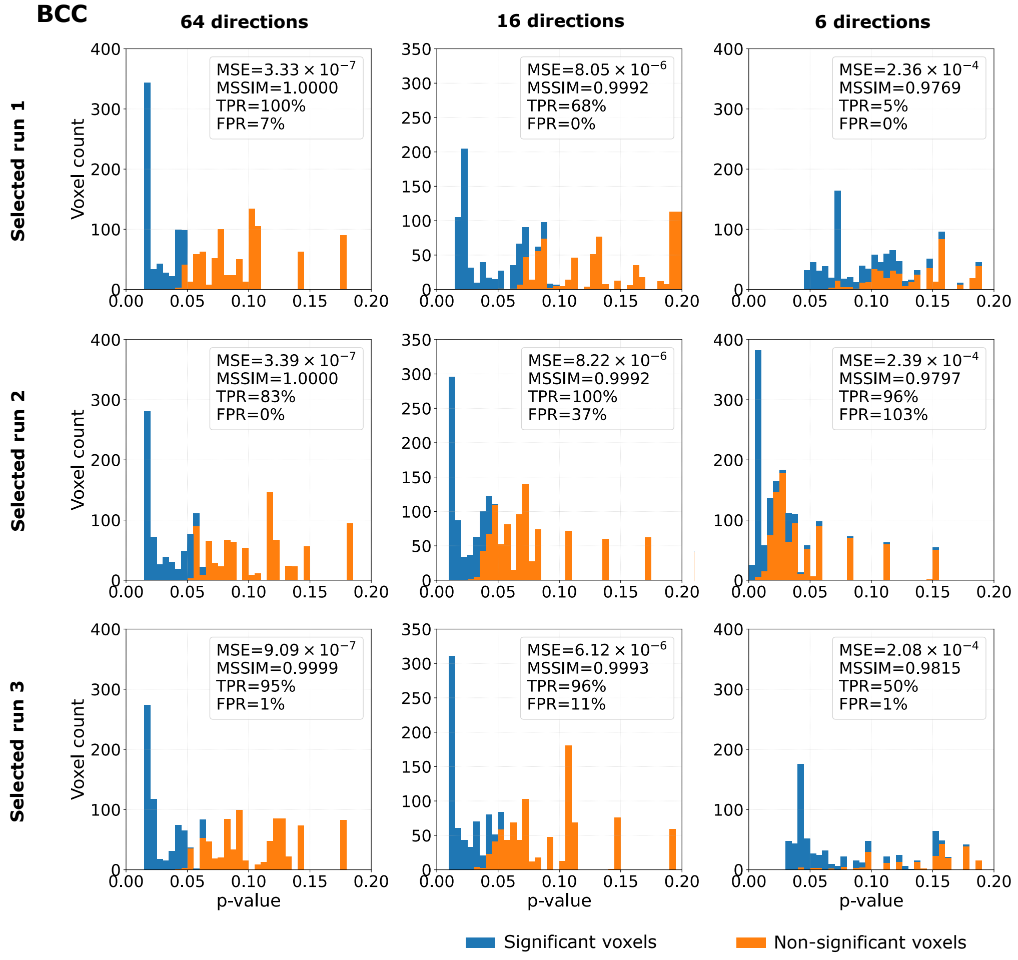


Figure S14: Histograms of voxel-wise $p$-values obtained from the tract-based spatial statistics (TBSS) analysis for the **RD parameter** estimated using the conventional NLLS method under **Configuration 3** over the skeleton of the body (BCC) and the genu (GCC) of the corpus callosum, and the right anterior corona radiata (ACR R). The figure presents three selected gradient rotations out of 100. For each rotation, three diffusion gradient distributions of input data are considered: 64, 16, and 6. Bins in blue refer to the voxels, which were statistically significant in the TBSS analysis based on NLLS-derived RD maps from the unrotated 64-direction scheme, while orange bins represent voxels that were not statistically significant in the reference analysis.

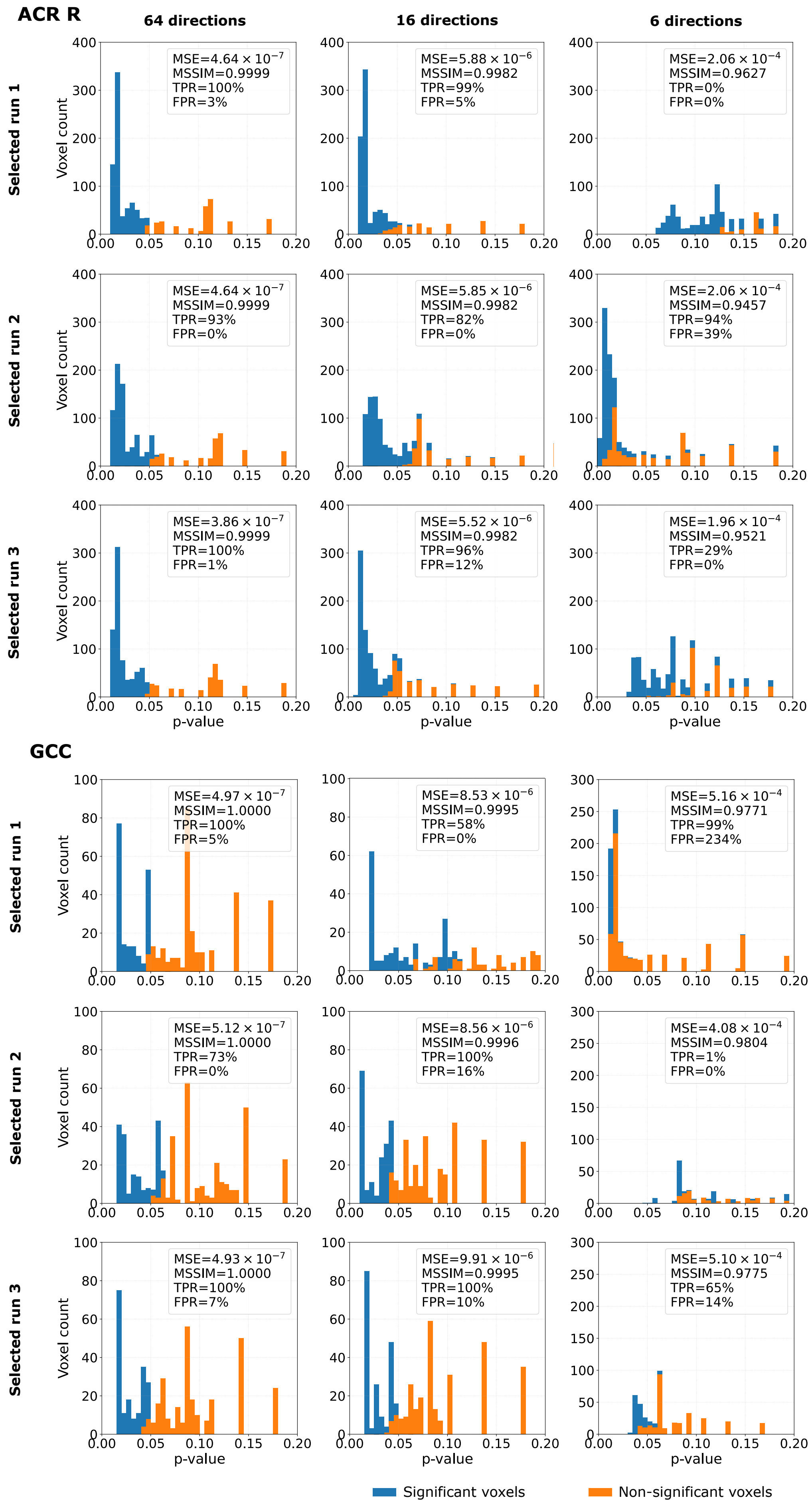


Figure S14 (continued)

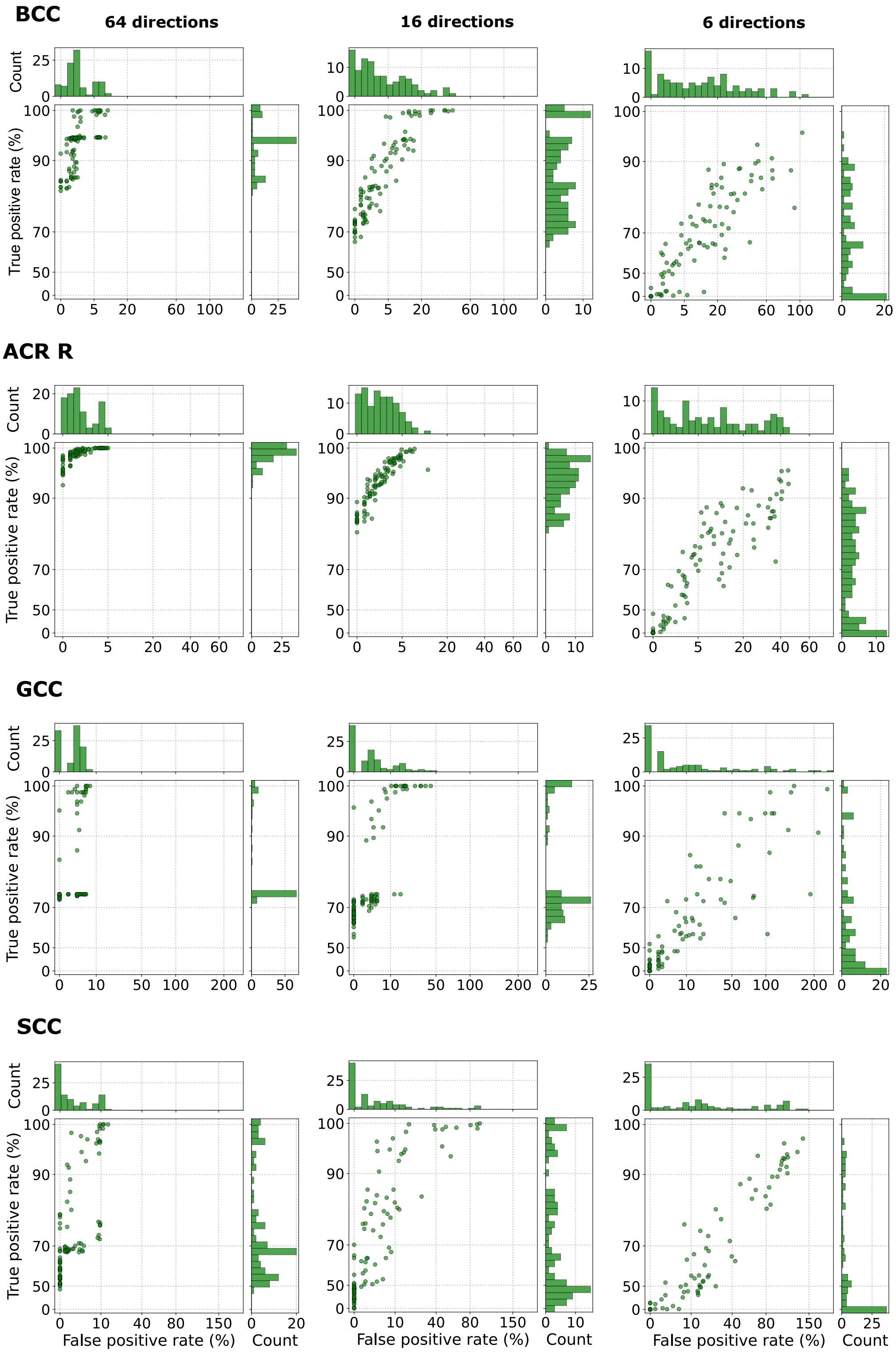


Figure S15: Relationship between the true positive rate and the false positive rate, obtained from the tract-based spatial statistics (TBSS) analysis for **RD parameter**, in the body (BCC), genu (GCC) and the splenium (SCC) of the corpus callosum, and the right anterior corona radiata (ACR R) using **Configuration 3**, with three different numbers of input data gradient directions (64, 16, and 6). Each point refers to a single gradient rotation. In total, 100 different gradient rotations were considered. The FPRs greater than 100% result from computing the values relative to the number of significant voxels identified in the reference analysis.

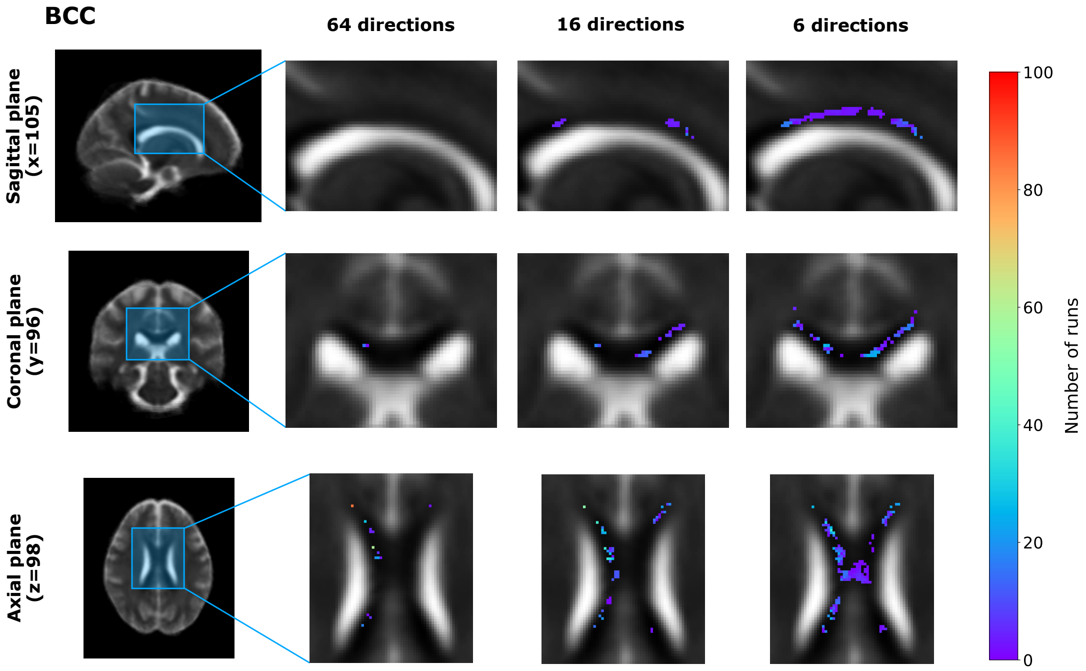


Figure S16: Visual representation of false positive voxels, obtained from the tract-based spatial statistics (TBSS) analysis for **RD parameter**, in the body of the corpus callosum (BCC) shown in the sagittal, coronal, and axial planes for three input gradient directions: 64, 16, and 6 (**Configuration 3**). False positive voxels correspond to locations identified as exhibiting statistically significant group differences in the estimated RD maps but not in the reference analysis. The color indicates the number of diffusion gradient rotations in which a given voxel was classified as a false positive.

Table S3: Decision agreement between rotation-based and reference ROI analyses across three different numbers of input data gradient directions (64, 16, and 6) (**Configuration 3**). The reference $p$-value and significance decision are reported for each ROI, while decision agreement indicates the percentage of the 100 gradient rotations that reproduced the statistical significance of the reference analysis ($p < 0.05$ or $p \geq 0.05$).

| **ROIs** | **Reference $p$-value** | **Reference decision** | **Decision agreement** | | |
|---|---|---|---|---|---|
| | | | **64 directions** | **16 directions** | **6 directions** |
| GCC | 0.9240 | Non-sig ($p \geq 0.05$) | 100% | 100% | 99% |
| BCC | 0.1910 | Non-sig ($p \geq 0.05$) | 100% | 100% | 71% |
| SCC | 0.0098 | Sig ($p < 0.05$) | 100% | 100% | 100% |
| ALIC R | 0.0192 | Sig ($p < 0.05$) | 100% | 97% | 58% |
| ACR R | 0.0700 | Non-sig ($p \geq 0.05$) | 100% | 100% | 96% |
| ACR L | 0.0494 | Sig ($p < 0.05$) | 93% | 44% | 4% |
| SCR R | 0.0852 | Non-sig ($p \geq 0.05$) | 100% | 90% | 75% |
| PCR R | 0.0954 | Non-sig ($p \geq 0.05$) | 100% | 100% | 96% |
| PTR R | 0.0012 | Sig ($p < 0.05$) | 100% | 100% | 76% |

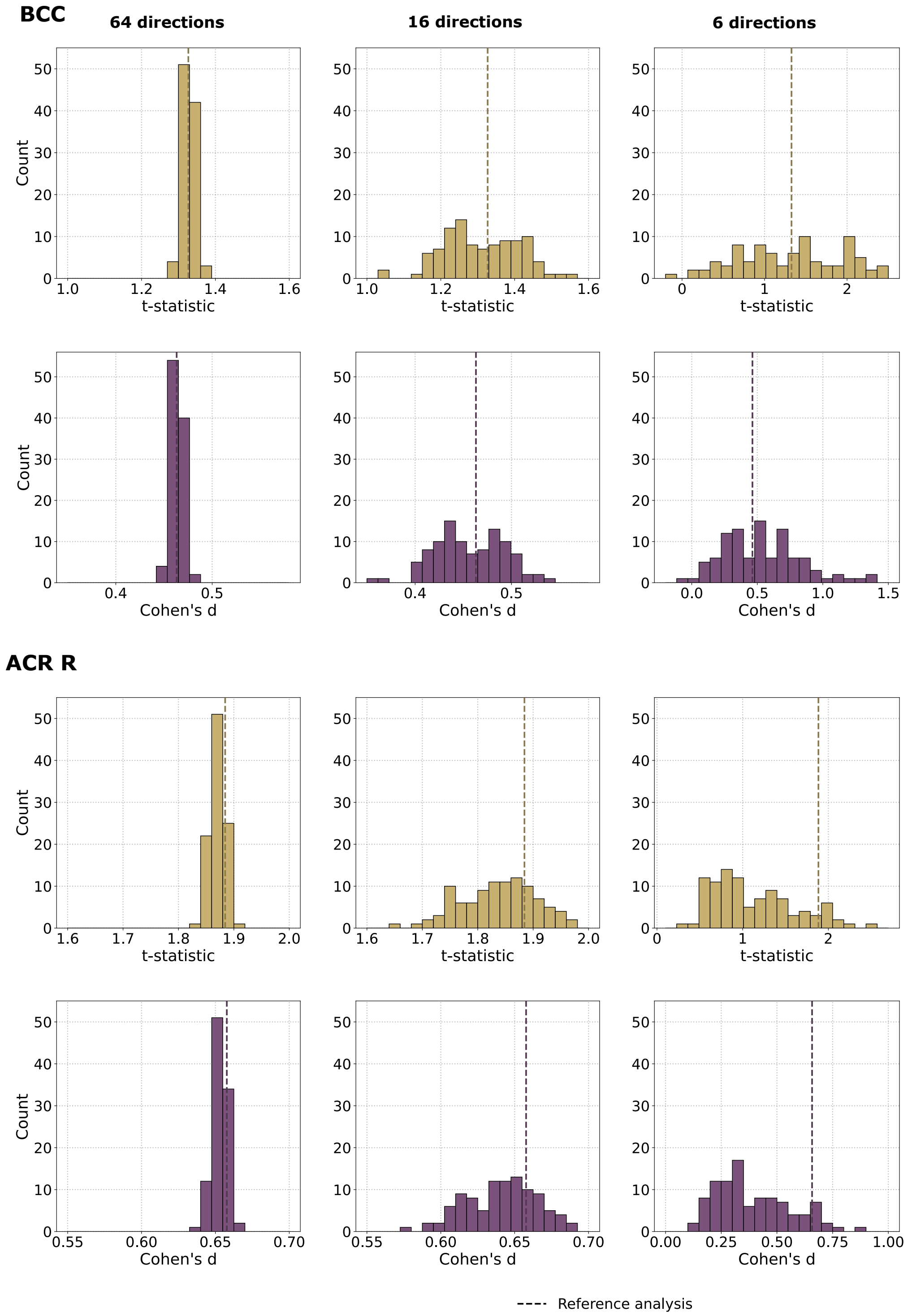


Figure S17: Distributions of $t$-statistics (top row) and Cohen's $d$ values (bottom row) in the body (BCC) and the genu (GCC) of the corpus callosum, and the right anterior corona radiata (ACR R) for the **RD parameter** estimated using the conventional NLLS method under **Configuration 3**, with three different numbers of input data gradient directions (64, 16, and 6). These statistics were calculated by comparing the mean RD values between patients with bipolar disorder (BD) and healthy controls (HC). Each histogram summarizes the results obtained from 100 gradient rotations. The dashed vertical line indicates the corresponding value from the reference analysis using the unrotated NLLS estimation.

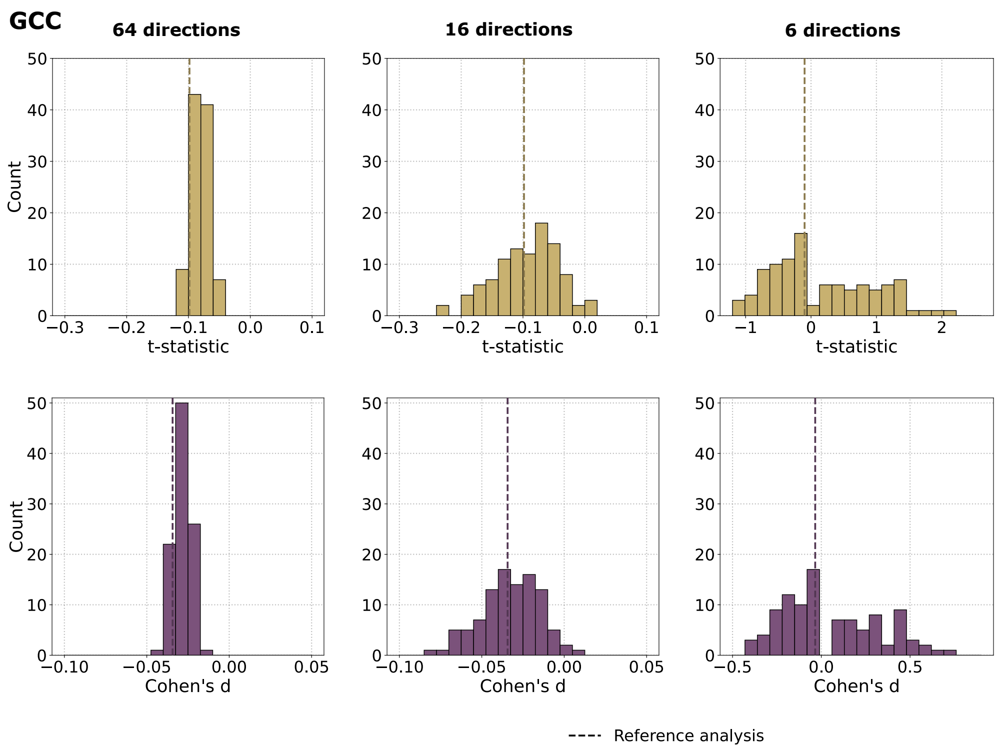


Figure S17 (continued)

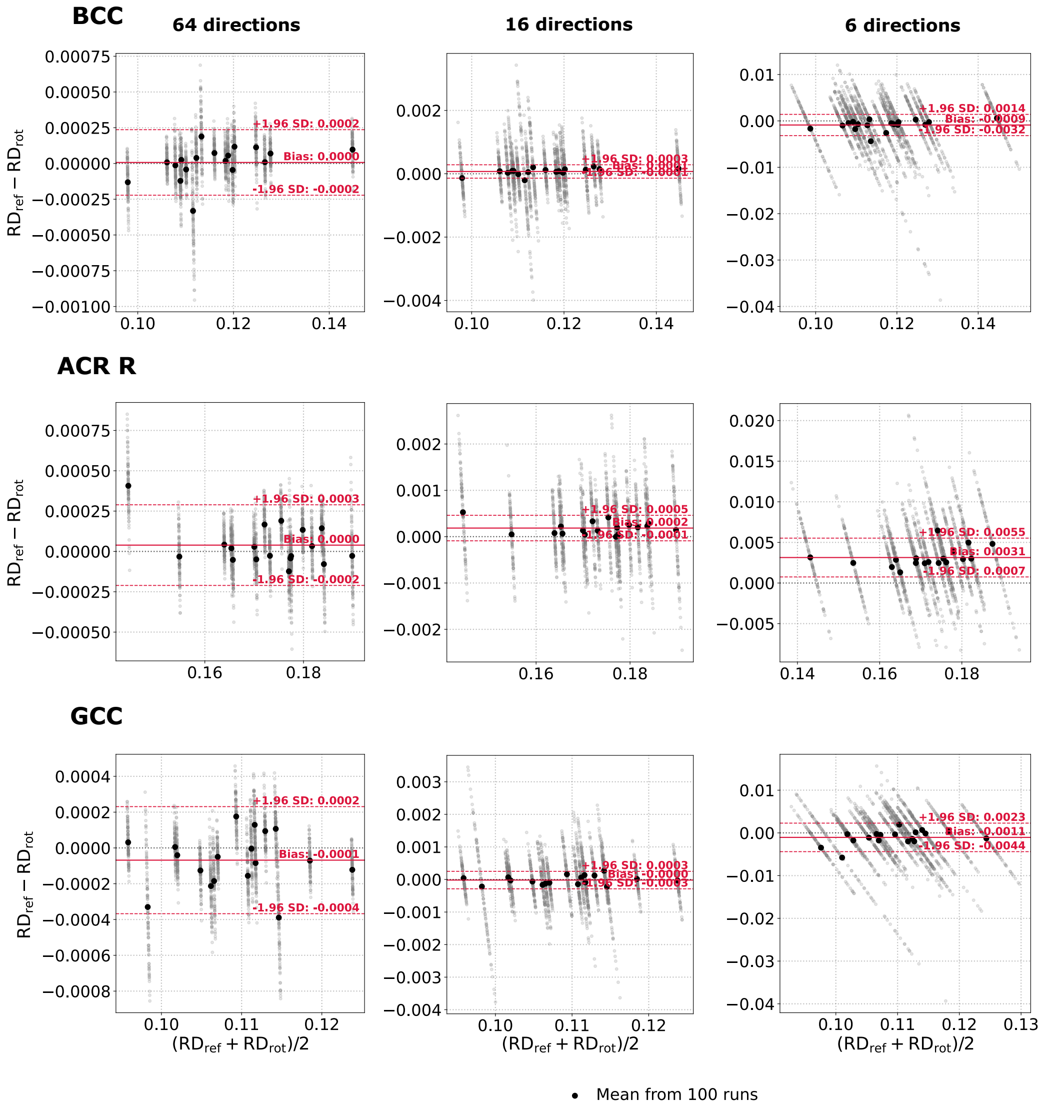


Figure S18: Bland-Altman plots of the **RD parameter** computed from the body (BCC) and genu of the corpus callosum (GCC), and right anterior corona radiata (ACR R) across all BD subjects. The reference values ($RD_{ref}$) were obtained using the NLLS estimation from an original, unrotated 64-direction scheme, whereas the $RD_{rot}$ values were estimated using NLLS for six gradient-direction schemes (64, 48, 32, 24, 16, and 6 directions) across 100 different rigid rotations of the gradient directions (**Configuration 3**). Gray points represent individual rotations (mean values over the ROI). A single black point corresponds to the average across 100 rotations for a single BD patient. Solid red lines indicate the bias, while dashed red lines denote 95% limits of agreement (±1.96 standard deviations).

### *2.5. Comparison of experimental configurations*

To evaluate whether the differences observed among the three configurations for the BCC in the main manuscript were consistent across a broader range of ROIs, we repeated the comparison for additional ones. The results are summarized in Figure S19. Overall, the same trends were evident across the additional ROIs, although the absolute values and distributions of the metrics varied by region. With a higher number of gradient directions, Configuration 3 generally produced the most stable results, demonstrating consistently high TPR and FPR close to zero. As the number of gradient directions decreased, the differences between Configurations 1 and 2, compared to Configuration 3, became more pronounced. For Configurations 1 and 2, TPR typically approached 100%, while FPR increased substantially. In contrast, Configuration 3 showed a decrease in TPR with the most reduced acquisition schemes, while FPR remained close to zero for most rotations. This pattern suggests that under highly reduced acquisitions, the two approaches exhibited different types of disagreement with the reference analysis: Configurations 1 and 2 tended to identify additional significant voxels, whereas Configuration 3 failed to replicate some of the significant voxels identified in the reference analysis.

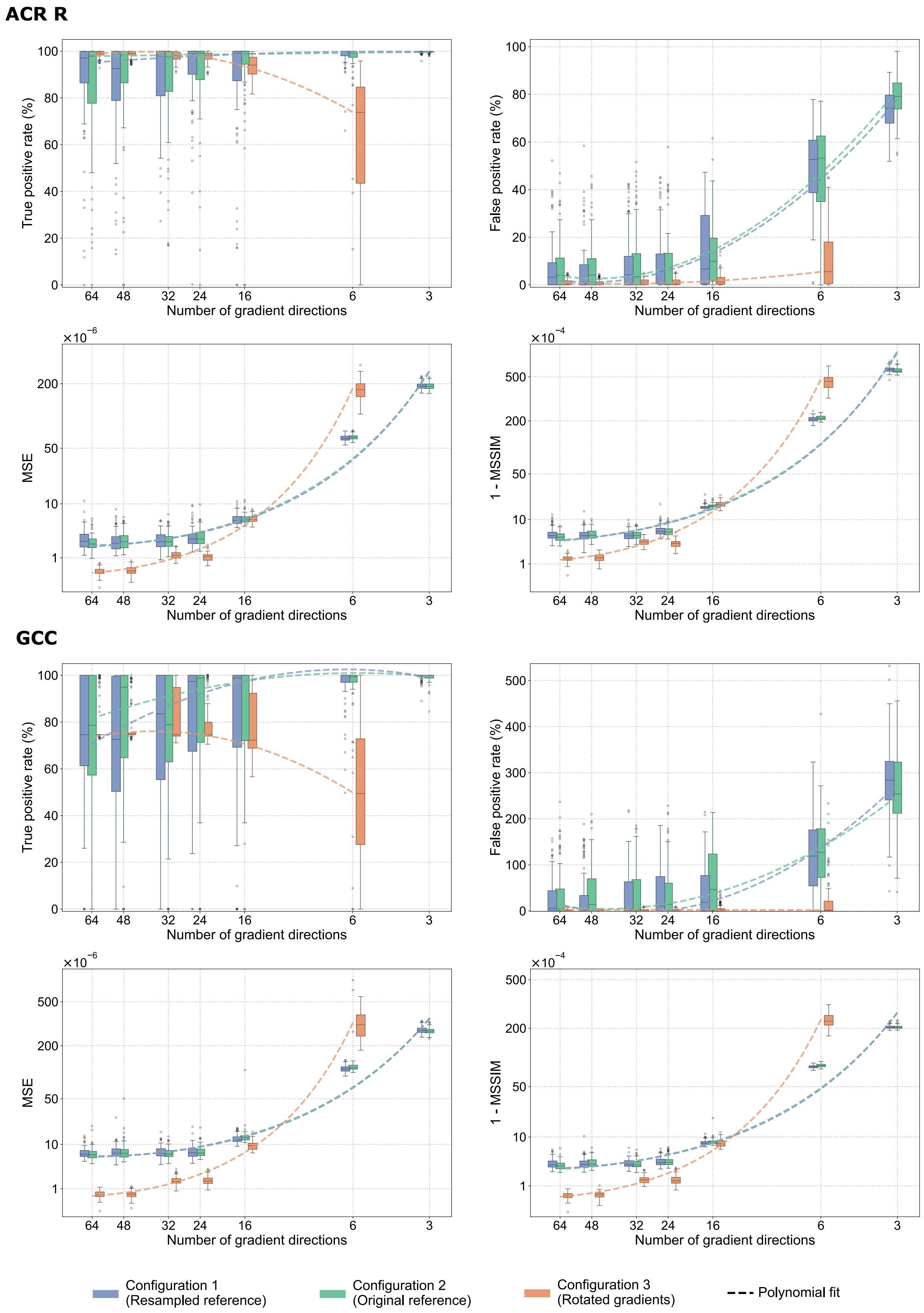


Figure S19: Boxplots presenting distributions of true positive rate, false positive rate, mean squared error (MSE), and 1−mean structural similarity index measure (1 − MSSIM) for **RD parameter** across the right anterior corona radiata (ACR R), and the genu of the corpus callosum (GCC) for the three experimental configurations: **Configuration 1 (Resampled reference)** – reference maps were derived using the NLLS from the resampled 64-direction dataset, **Configuration 2 (Original reference)** – reference maps were derived using the NLLS from the original 64-direction dataset, **Configuration 3 (Rotated gradients)** – a non-DL baseline representing RD calculated using the NLLS fitting on rotated gradient sets. Boxes indicate the interquartile range (IQR), horizontal lines denote the median, whiskers extend to 1.5 × IQR, and individual points indicate outliers. The dashed line represents a second-order polynomial regression fitted to the median values of the metrics, illustrating the overall trend of changes.

### 3. Bipolar disorder: fractional anisotropy

#### 3.1. Comparison of between-group differences between original and resampled datasets

To evaluate whether the resampling procedure influenced the identification of regions with significant between-group differences, we repeated the reference Tract-Based Spatial Statistics (TBSS) analysis described in the main manuscript using the original, non-resampled dataset. Table S4 summarizes the statistically significant ROIs identified in the original dataset. The same ROIs were found to be statistically significant in both the original and resampled datasets, with the group difference consistently indicating that the BD group had lower fractional anisotropy (FA) values than the HC group (BD < HC). These findings suggest that the resampling procedure did not change the set of ROIs identified as showing significant between-group differences, nor did it affect the direction of these effects.

Table S4: White matter regions identified by tract-based spatial statistics (TBSS) analysis of **fractional anisotropy (FA)** in patients with bipolar disorder (BD) compared with healthy controls (HC) for **Configuration 2**. The table includes only regions that show a significant increase in RD (BD > HC) relative to HC after the threshold-free cluster enhancement (TFCE) correction ($p < 0.05$). Significant voxels are reported as $n/N$ (%), where $n$ is the number of voxels showing a statistically significant difference, and $N$ is the total number of voxels within the corresponding white matter region. Peak T-stat represents the maximum $t$-statistic within the significant portion of the region. Coordinates (X, Y, Z) indicate the location of the peak voxel in MNI space. Corrected $p$-value refers to the TFCE family-wise error-corrected $p$-value at the peak voxel.

| Parameter | ROIs | Significant voxels | Peak T-stat | Coordinates (X,Y,Z) | Corrected p-value |
|---|---|---|---|---|---|
| FA (BD < HC) | GCC | 996/1787 (55.74%) | 4.516 | -8, 29, -1 | 0.0298 |
| | BCC | 2117/3245 (65.24%) | 5.355 | -6, 15, 21 | 0.0220 |
| | SCC | 607/2364 (25.68%) | 3.497 | -18, -36, 32 | 0.0352 |
| | RIC L | 306/751 (40.75%) | 3.641 | -39, -29, -2 | 0.0380 |
| | ACR R | 638/1716 (37.18%) | 3.579 | 16, 19, 29 | 0.0180 |
| | SCR R | 311/1499 (20.75%) | 3.669 | 20, -17, 39 | 0.0174 |
| | PCR R | 83/724 (11.46%) | 2.708 | 20, -27, 36 | 0.0186 |
| | PCR L | 85/732 (11.61%) | 2.948 | -19, -39, 33 | 0.0354 |
| | EC L | 180/1387 (12.98%) | 5.768 | -33, -20, -1 | 0.0382 |
| | CG R | 106/397 (26.70%) | 3.830 | 9, 15, 30 | 0.0370 |
| | SLF L | 522/1429 (36.53%) | 3.631 | -37, -22, 29 | 0.0330 |

#### 3.2. Configuration 1 (Resampled reference)

The resampled 64-direction dataset serves as the reference for the DL-based DTI. The reference FA maps were computed from this dataset using the nonlinear least-squares (NLLS) fitting procedure, and then used as pseudo-ground truth for the DL network training procedure. The reduced inputs with $k = \{48, 32, 24, 16, 6, 3\}$ gradient directions were prepared by selecting the first $k$ volumes from the resampled dataset.

Figures S20, S21, S22, S23, S24, and Table S5 display results similar to those presented in the main manuscript for RD, but focus on FA estimated using DL under Configuration 1. Overall, estimating FA appears to be more challenging than estimating RD, particularly under reduced acquisition schemes. This is evident from the higher MSE and lower MSSIM values, with the differences becoming more noticeable in the 6- and 3-direction schemes. The TBSS metrics also reveal distinct patterns for FA. Across the analyzed ROIs, the TPR values were generally less variable and closer to 100% than those observed for RD, even for the full 64-direction scheme. As the number of gradient directions decreased, the TPR approached 100% more quickly. In contrast, the FPR values were generally higher than for RD, even with 64 directions. A marked increase in FPR values, along with reduced variability, was already noticeable for 6 gradient directions, whereas a similar trend for RD was most apparent for 3 directions. Thus, although TPR values were high, FA estimates often identified additional statistically significant voxels that were not identified in the reference analysis. Supporting these findings, the distributions of ROI-based $t$-statistics and Cohen's $d$ values displayed a more significant shift toward higher values for FA compared to RD, with this shift evident for some ROIs even under the 64-direction scheme. The Bland-Altman analyses indicated a greater tendency toward lower estimated FA values relative to the reference, even with 64 directions, and this difference became more pronounced as the number of directions decreased. Despite these differences, there was substantial variability in the TBSS metrics across runs, consistent with the findings for RD.

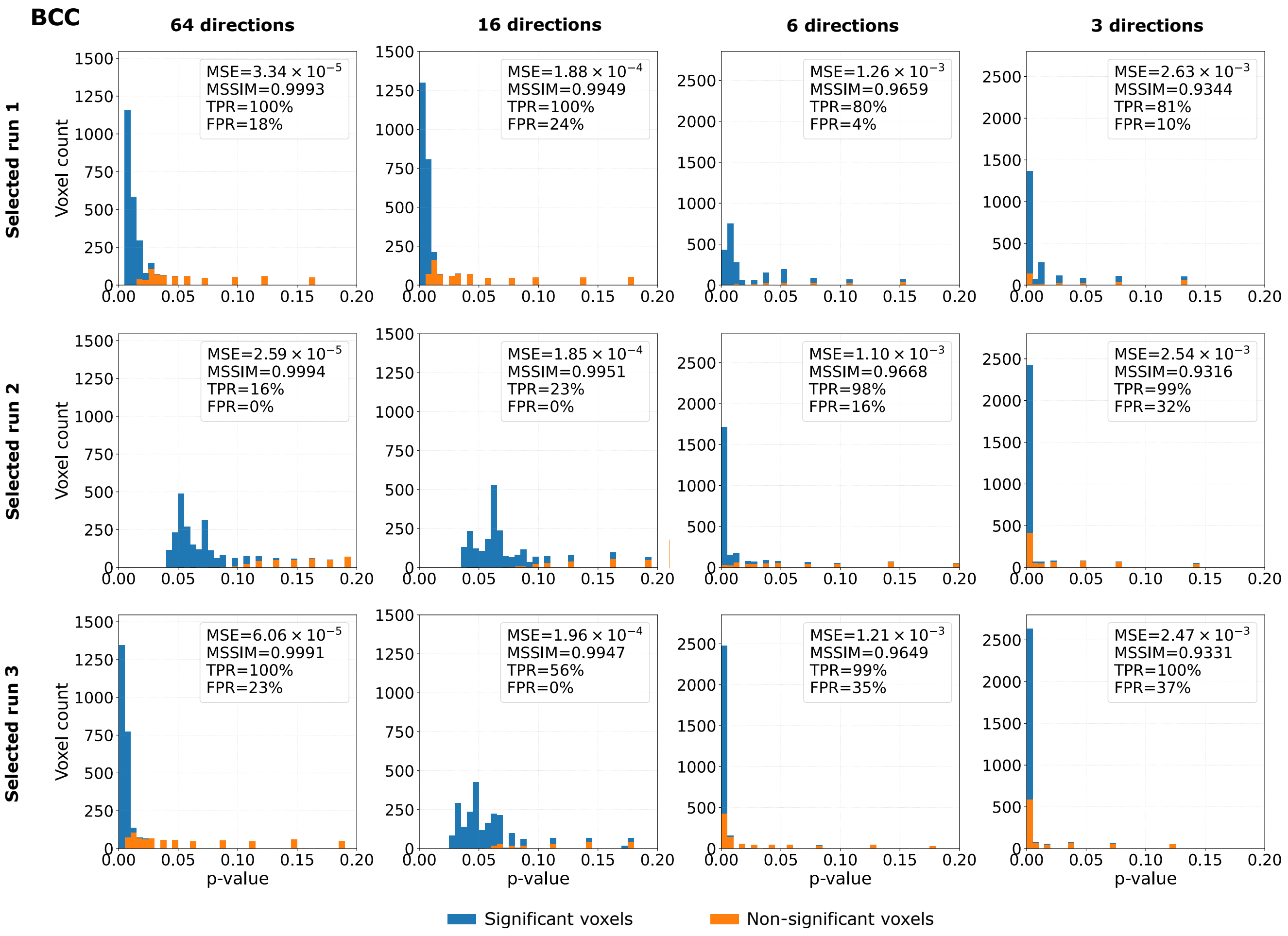


Figure S20: Histograms of voxel-wise $p$-values obtained from the tract-based spatial statistics (TBSS) analysis for the **FA parameter** estimated using a deep learning model following **Configuration 1** over the skeleton of the body of the corpus callosum (BCC) and the right anterior corona radiata (ACR R). The figure presents three selected runs over 100 runs characterized with a comparable mean squared error (MSE) and mean structural similarity index measure (MSSIM), while variable true positive and false positive rates (TPR, FPR). For each run, four diffusion gradient distributions of input data are considered: 64, 16, 6, and 3. Bins in blue refer to the voxels, which were statistically significant in the TBSS analysis based on the standard NLLS-derived DTI (no deep learning applied), while orange bins show the voxels statistically non-significant in the TBSS analysis based on NLLS-derived DTI.

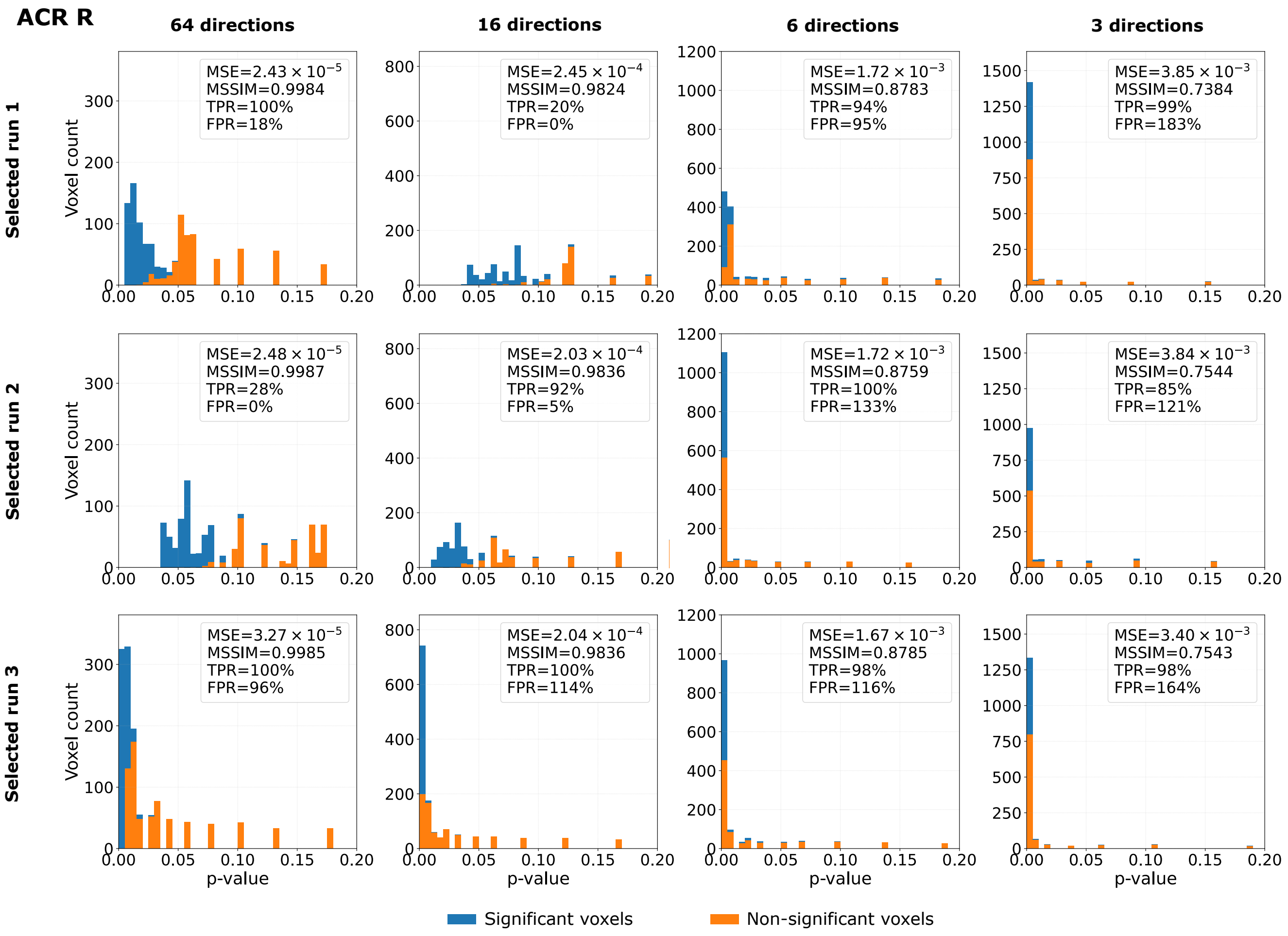


Figure S20 (continued)

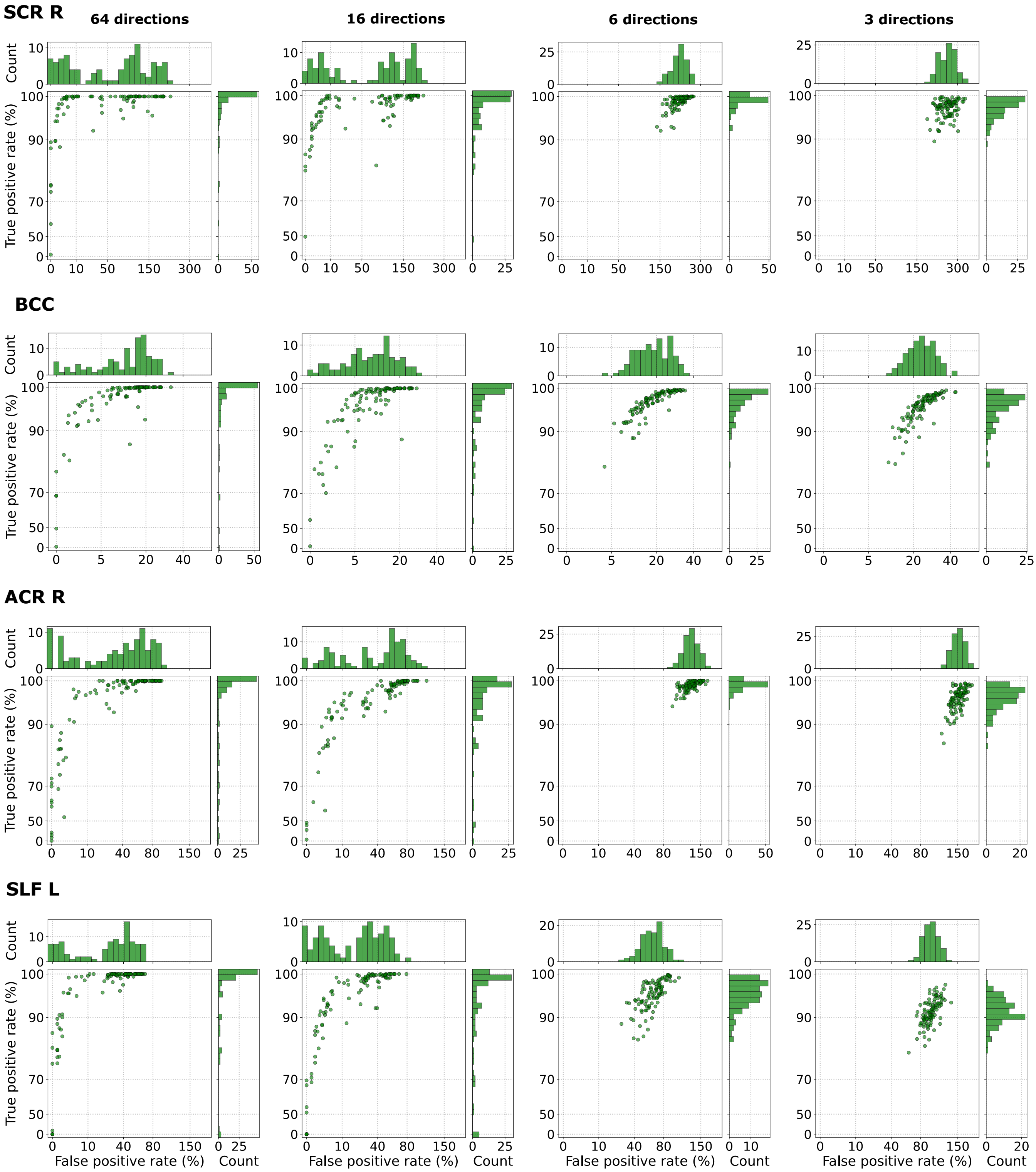


Figure S21: Relationship between the true positive rate and the false positive rate, obtained from the tract-based spatial statistics (TBSS) analysis for **FA parameter**, in the right superior corona radiata (SCR R), the body of the corpus callosum (BCC), the right anterior corona radiata (ACR R), and the left superior longitudinal fasciculus (SLF L) using **Configuration 1**, with four different numbers of input data gradient directions (64, 16, 6, and 3). Each point refers to a single run. In total, 100 runs were considered. The FPRs greater than 100% result from computing the values relative to the number of significant voxels identified in the reference analysis.

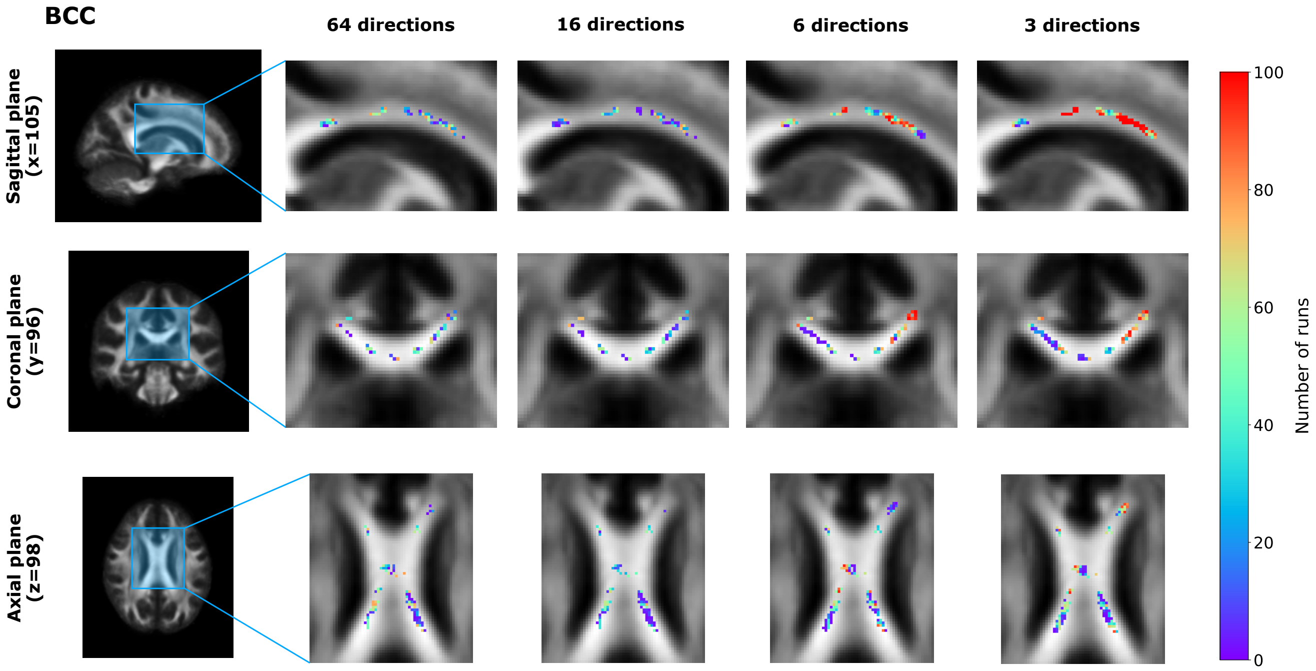


Figure S22: Visual representation of false positive voxels, obtained from the tract-based spatial statistics (TBSS) analysis for **FA parameter**, in the body of the corpus callosum (BCC) shown in the sagittal, coronal, and axial planes for four input gradient directions: 64, 16, 6, and 3 (**Configuration 1**). False positive voxels correspond to locations identified as exhibiting statistically significant group differences in the estimated FA maps but not in the reference analysis. The color indicates the number of runs in which a given voxel was classified as a false positive.

Table S5: Decision agreement between DL-based and reference ROI analyses across four different numbers of input data gradient directions (64, 16, 6, and 3) for **FA parameter** (**Configuration 1**). The reference $p$-value and significance decision are reported for each ROI, while decision agreement indicates the percentage of runs out of 100 that reproduced the statistical significance of the reference analysis ($p < 0.05$ or $p \geq 0.05$).

| ROIs | Reference $p$-value | Reference decision | Decision agreement | | | |
|---|---|---|---|---|---|---|
| | | | 64 directions | 16 directions | 6 directions | 3 directions |
| GCC | 0.0164 | Sig ($p < 0.05$) | 100% | 98% | 99% | 100% |
| BCC | 0.0048 | Sig ($p < 0.05$) | 100% | 100% | 100% | 100% |
| SCC | 0.1058 | Non-sig ($p \geq 0.05$) | 77% | 93% | 56% | 14% |
| RIC L | 0.0376 | Sig ($p < 0.05$) | 79% | 78% | 100% | 97% |
| ACR R | 0.0754 | Non-sig ($p \geq 0.05$) | 80% | 87% | 1% | 1% |
| SCR R | 0.0792 | Non-sig ($p \geq 0.05$) | 54% | 52% | 12% | 7% |
| PCR R | 0.1470 | Non-sig ($p \geq 0.05$) | 94% | 97% | 69% | 66% |
| PCR L | 0.3710 | Non-sig ($p \geq 0.05$) | 100% | 100% | 87% | 88% |
| EC L | 0.1156 | Non-sig ($p \geq 0.05$) | 94% | 97% | 42% | 7% |
| CG R | 0.1090 | Non-sig ($p \geq 0.05$) | 84% | 92% | 23% | 0% |
| SLF L | 0.0222 | Sig ($p < 0.05$) | 98% | 94% | 97% | 99% |

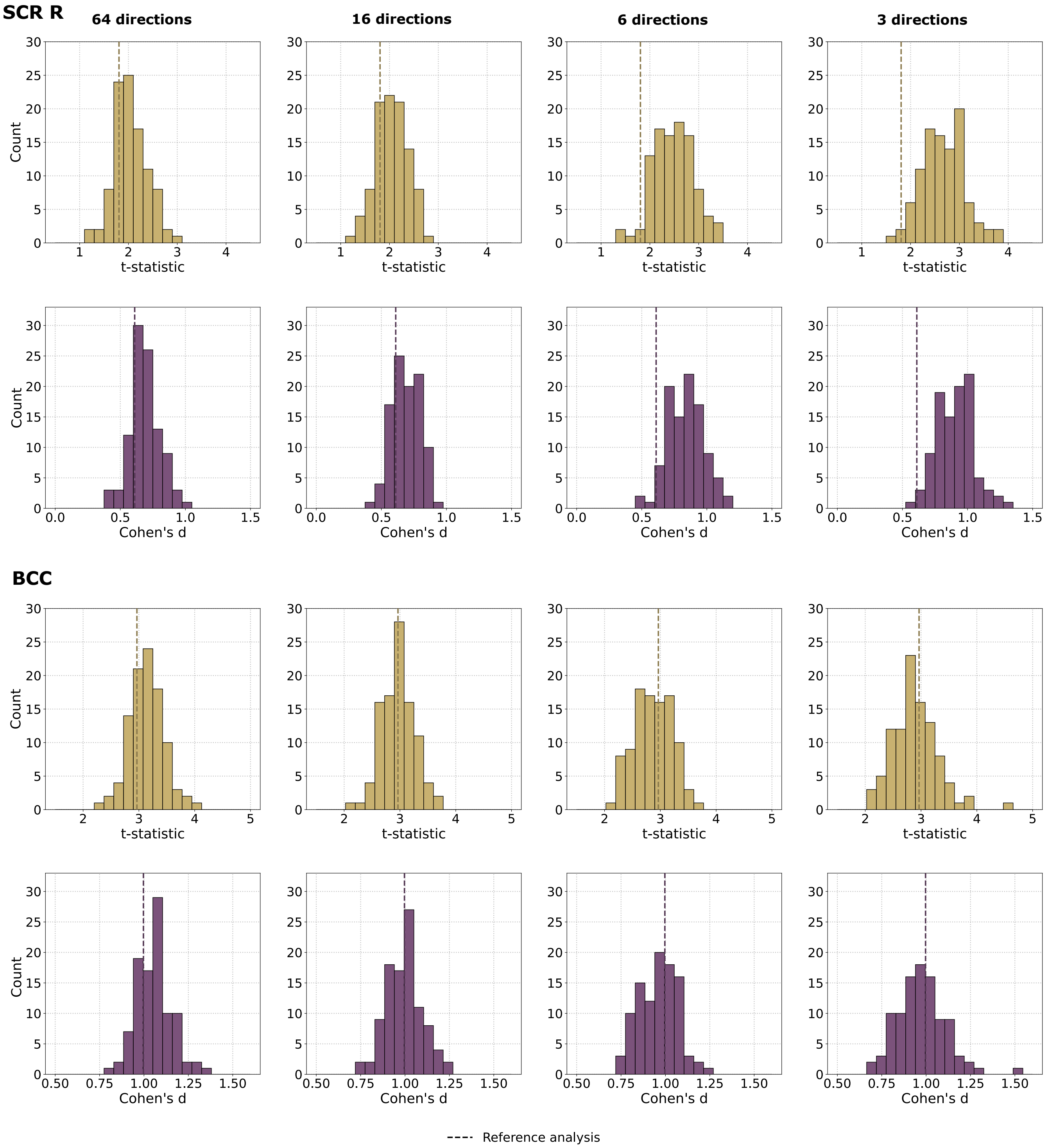


Figure S23: Distributions of $t$-statistics (top row) and Cohen's $d$ values (bottom row) in the right superior corona radiata (SCR R), the body of the corpus callosum (BCC), and the right anterior corona radiata (ACR R) for the **FA parameter** estimated using a deep learning model, with four different numbers of input data gradient directions (64, 16, 6, and 3) (**Configuration 1**). These statistics were calculated by comparing the mean FA values between patients with bipolar disorder (BD) and healthy controls (HC). Each histogram summarizes the results obtained from 100 runs. The dashed vertical line indicates the corresponding value from the reference analysis using the NLLS estimation.

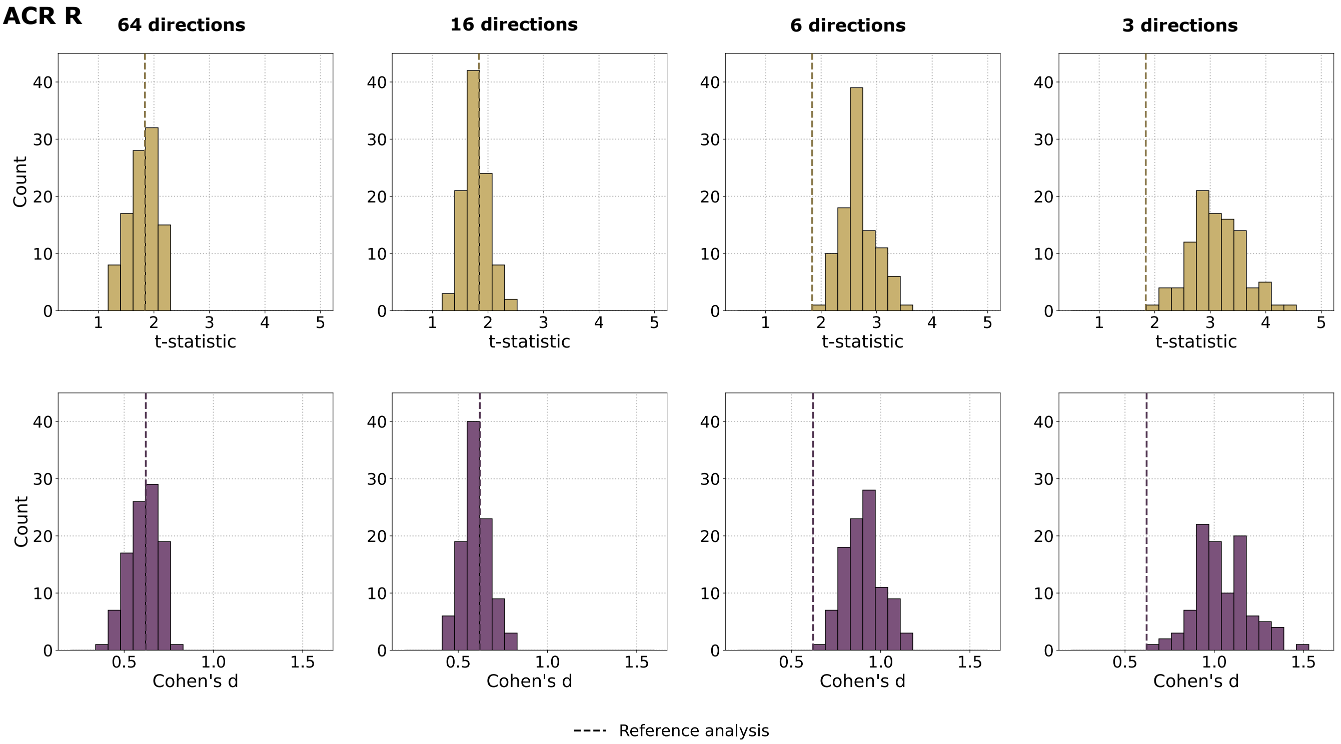


Figure S23 (continued)

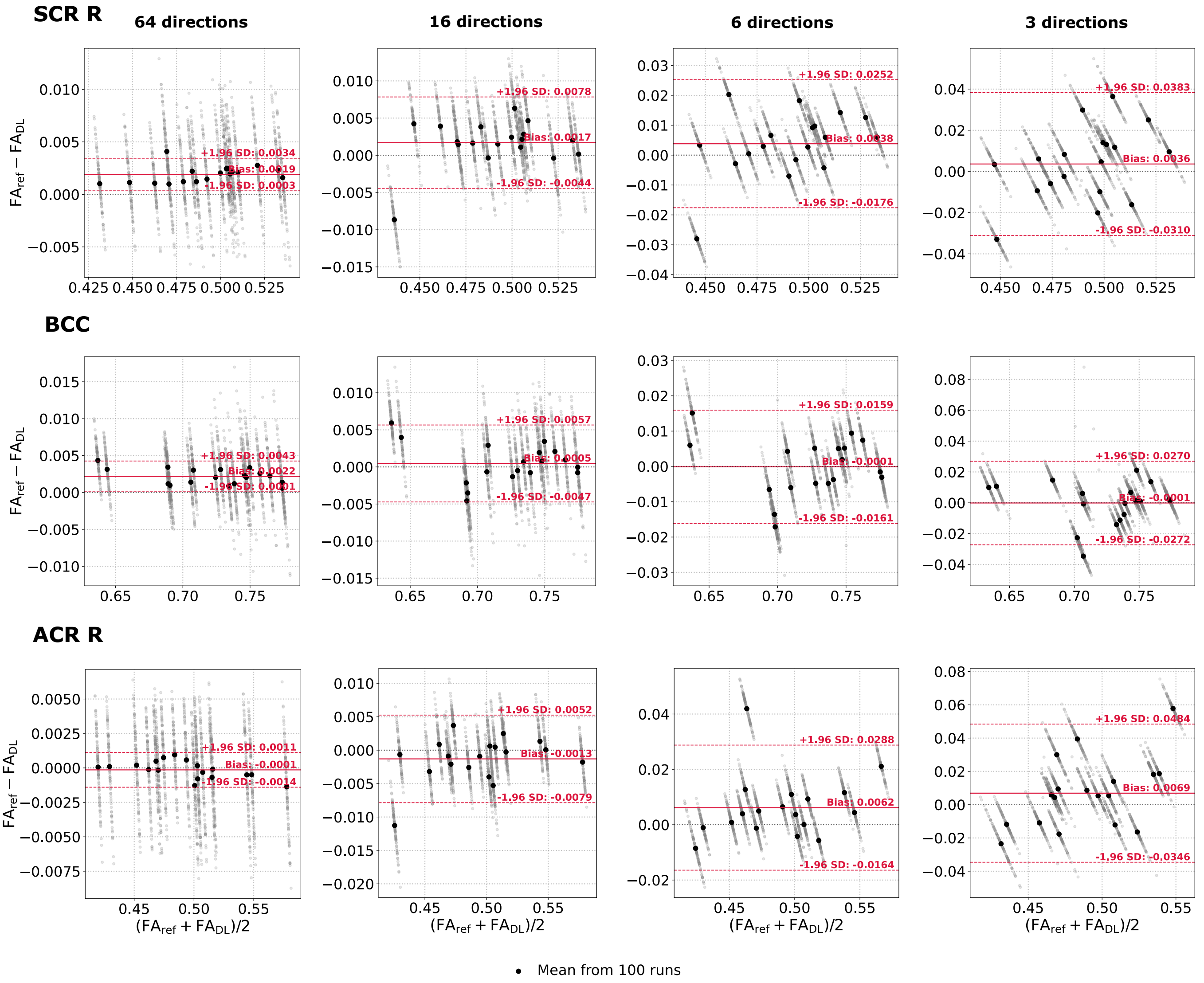


Figure S24: Bland-Altman plots of the **FA parameter** computed from the right superior corona radiata (SCR R), the body of the corpus callosum (BCC), and the right anterior corona radiata (ACR R) across all BD subjects. The reference values ($FA_{ref}$) were obtained using the NLLS estimation from a resampled 64-direction scheme, whereas the predicted $FA_{DL}$ values were generated using the deep learning model (**Configuration 1**) considering four different numbers of input data gradient directions (64, 16, 6, and 3). Gray points represent individual model runs (mean values over the ROI). A single black point corresponds to the average across 100 runs for a single BD patient. Solid red lines indicate the bias, while dashed red lines denote 95% limits of agreement (±1.96 standard deviations).

### 4. Schizophrenia: radial diffusivity

#### *4.1. Between-group difference*

To understand whether the observed trends were unique to bipolar disorder or also applicable to other conditions, we conducted a similar analysis on a group of individuals with schizophrenia (SCH). First, we examined the differences in white matter microstructure between SCH and HC using TBSS. The results of our analysis are summarized in Table S6, which also defines the abbreviations used for the ROIs.

Our findings revealed that the SCH group exhibited significantly higher RD compared to the HC group (SCH > HC). The most pronounced changes were observed in the GCC, BCC, and SCC. Additionally, we noted widespread differences in several areas, including the ALIC R and L, ACR R and L, SCR R and L, PCR R and L, PTR R and L, RIC R and L, SS R and L, EC R and L, and SLF R and L. These findings align with previous studies that have reported white matter microstructural abnormalities in schizophrenia (Seal et al., 2008; Scheel et al., 2013; Holleran et al., 2014; Rigucci et al., 2016; Zeng et al., 2016; Fu et al., 2019; Lee et al., 2020; Gao et al., 2025; Mamah et al., 2025).

Table S6: White matter regions identified by tract-based spatial statistics (TBSS) analysis of **radial diffusivity (RD)** in patients with schizophrenia (SCH) compared with healthy controls (HC) for **Configuration 1**. The table includes only regions that show a significant increase in RD (SCH > HC) in SCH relative to HC after the threshold-free cluster enhancement (TFCE) correction ($p < 0.05$). Significant voxels are reported as $n/N$ (%), where $n$ is the number of voxels showing a statistically significant difference, and $N$ is the total number of voxels within the corresponding white matter region. Peak T-stat represents the maximum $t$-statistic within the significant portion of the region. Coordinates (X, Y, Z) indicate the location of the peak voxel in MNI space. Corrected $p$-value refers to the TFCE family-wise error-corrected $p$-value at the peak voxel.

| Parameter | ROIs | Significant voxels | Peak T-stat | Coordinates (X,Y,Z) | Corrected p-value | Studies reporting similar findings |
|---|---|---|---|---|---|---|
| RD (SCH > HC) | GCC | 1326/1785 (74.29%) | 4.544 | 15, 30, 16 | 0.0050 | Scheel et al. (2013); Holleran et al. (2014); Gao et al. (2025) |
| | BCC | 2091/3190 (65.55%) | 5.214 | -9, 8, 25 | 0.0044 | Scheel et al. (2013); Holleran et al. (2014); Fu et al. (2019); Lee et al. (2020); Gao et al. (2025) |
| | SCC | 1213/2400 (50.54%) | 5.206 | 18, -45, 11 | 0.0232 | Holleran et al. (2014); Fu et al. (2019); Lee et al. (2020); Gao et al. (2025) |
| | ALIC R | 548/804 (68.16%) | 4.729 | 19, 6, 13 | 0.0088 | Scheel et al. (2013) |
| | ALIC L | 418/834 (50.12%) | 3.664 | -23, -2, 17 | 0.0044 | Scheel et al. (2013); Lee et al. (2020); Gao et al. (2025) |
| | PLIC R | 229/877 (26.11%) | 2.325 | 28, -22, 11 | 0.0272 | Holleran et al. (2014) |
| | RIC R | 346/755 (45.83%) | 3.598 | 30, -22, 5 | 0.0266 | Scheel et al. (2013); Lee et al. (2020) |
| | RIC L | 96/771 (12.45%) | 3.573 | -33, -23, -2 | 0.0114 | Lee et al. (2020) |
| | ACR R | 1092/1660 (65.78%) | 5.094 | 27, 20, 21 | 0.0078 | Scheel et al. (2013); Fu et al. (2019); Gao et al. (2025) |
| | ACR L | 1494/1777 (84.07%) | 5.852 | -21, 16, 29 | 0.0044 | Scheel et al. (2013); Fu et al. (2019); Gao et al. (2025) |
| | SCR R | 441/1456 (30.29%) | 4.022 | 17, 15, 30 | 0.0050 | Scheel et al. (2013); Gao et al. (2025) |
| | SCR L | 396/1434 (27.62%) | 4.399 | -23, 9, 19 | 0.0042 | Scheel et al. (2013); Lee et al. (2020); Gao et al. (2025) |
| | PCR R | 194/727 (26.69%) | 3.712 | 25, -41, 26 | 0.0256 | Scheel et al. (2013) |
| | PCR L | 199/713 (27.91%) | 3.537 | -20, -47, 31 | 0.0116 | Scheel et al. (2013); Fu et al. (2019); Lee et al. (2020) |
| | PTR R | 808/1142 (70.75%) | 4.672 | 34, -62, 6 | 0.0194 | Scheel et al. (2013); Lee et al. (2020); Gao et al. (2025) |
| | PTR L | 690/1064 (64.85%) | 4.024 | -38, -46, 3 | 0.0132 | Fu et al. (2019) |
| | SS R | 156/564 (27.66%) | 3.408 | 41, -31, -6 | 0.0246 | Scheel et al. (2013); Zeng et al. (2016); Fu et al. (2019) |
| | SS L | 184/509 (36.15%) | 3.554 | -42, -37, -10 | 0.0132 | Scheel et al. (2013); Zeng et al. (2016) |
| | EC R | 229/1255 (18.25%) | 3.724 | 16, 20, -11 | 0.0130 | Seal et al. (2008); Scheel et al. (2013); Holleran et al. (2014); Lee et al. (2020) |
| | EC L | 828/1396 (59.31%) | 4.021 | -33, 0, 0 | 0.0114 | Seal et al. (2008); Scheel et al. (2013) |
| | SLF R | 618/1431 (43.19%) | 4.203 | 34, -42, 32 | 0.0164 | Scheel et al. (2013); Holleran et al. (2014); Fu et al. (2019); Mamah et al. (2025) |
| | SLF L | 156/1479 (10.55%) | 4.217 | -35, 7, 22 | 0.0226 | Rigucci et al. (2016); Zeng et al. (2016); Fu et al. (2019); Mamah et al. (2025) |

GCC – genu of the corpus callosum
BCC – body of the corpus callosum
SCC – splenium of the corpus callosum
ALIC R and L – right and left anterior limb of the internal capsule
PLIC R – right posterior limb of the internal capsule
RIC R and L – right and left retrolenticular part of the internal capsule
ACR R and L – right and left anterior corona radiata
SCR R and L – right and left superior corona radiata
PCR R and L – right and left posterior corona radiata
PTR R and L – right and left posterior thalamic radiation
SS R and L – right and left sagittal stratum
EC R and L – right and left external capsule
SLF R and L – right and left superior longitudinal fasciculus

#### 4.2. Configuration 1 (Resampled reference)

The resampled 64-direction dataset serves as the reference for the DL-based DTI. The reference RD maps were computed from this dataset using the nonlinear least-squares (NLLS) fitting procedure, and then used as pseudo-ground truth for the DL network training procedure. The reduced inputs with $k = \{48, 32, 24, 16, 6, 3\}$ gradient directions were prepared by selecting the first $k$ volumes from the resampled dataset.

Figures S25, S26, S27, S28, S29, S30, and Table S7 present results similar to those in the main manuscript for the BD, but specifically for the schizophrenia and the RD estimated using DL under Configuration 1. Overall, the results showed trends akin to those observed in the BD group. The visual assessment indicated that the estimated RD maps closely resembled the reference maps across the evaluated acquisition schemes. As anticipated, the voxel-wise differences increased as the number of gradient directions decreased. The TBSS analysis showed notable variability in TPR and FPR across different runs, despite relatively stable MSE and MSSIM values. This variability was evident even when the full set of 64 gradient directions was available and became more pronounced as the number of directions was reduced. Under limited acquisition schemes, TPR generally remained high or approached 100%, while FPR increased, indicating that additional voxels were classified as significant. The distributions of these metrics followed trends similar to those observed for BD, although their absolute values varied across ROIs. The ROI-based statistical analyses revealed corresponding variability in the estimated t-statistics and Cohen's d values across independent training repetitions. As the number of gradient directions decreased, the estimated statistical effects tended to deviate from the reference results. The decision agreement analysis similarly indicated that independent runs did not consistently reproduce the reference significance decisions. The Bland-Altman analyses also demonstrated increasing discrepancies between the estimated RD values and the reference values as the number of gradient directions declined. In comparison with BD, the schizophrenia analysis identified statistically significant group differences across a greater number of ROIs. Nevertheless, the overall relationships between image-level similarity, TBSS performance, and variability across different runs remained consistent between the two groups. Thus, the schizophrenia results further support the observation that the variability in downstream statistical outcomes is not unique to the BD group or to the specific ROIs examined in the main manuscript.

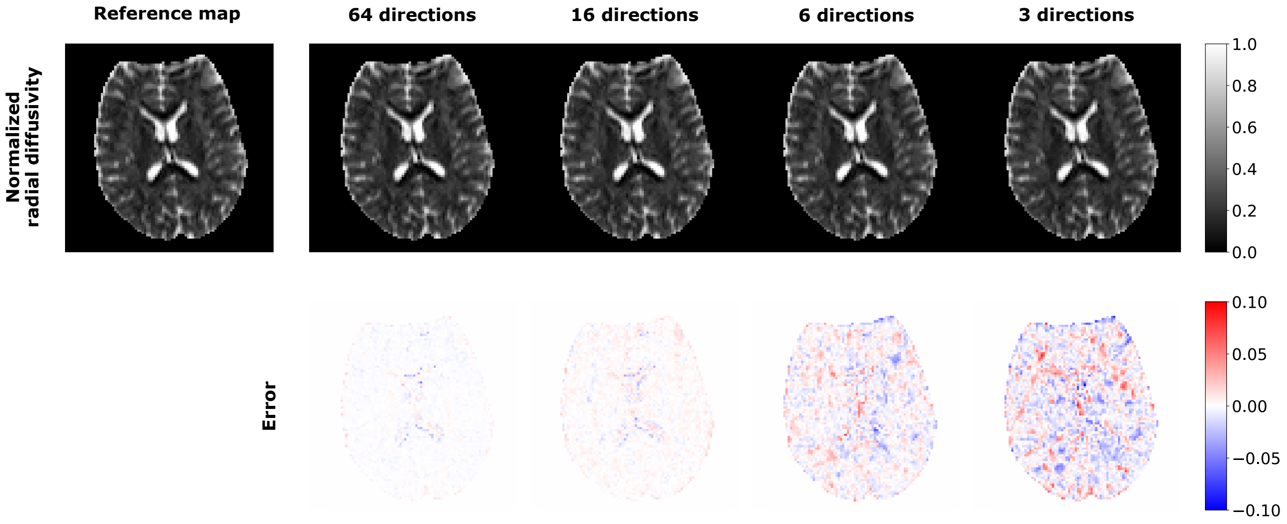


Figure S25: Visual inspection of **normalized radial diffusivity** estimated using single runs of the deep learning model and the corresponding voxel-wise differences between DL-based maps and the reference map derived from the resampled 64-direction scheme using the NLLS estimation (**Configuration 1**). Four input diffusion gradient schemes are considered for DL-based estimation: 64, 16, 6, and 3 directions.

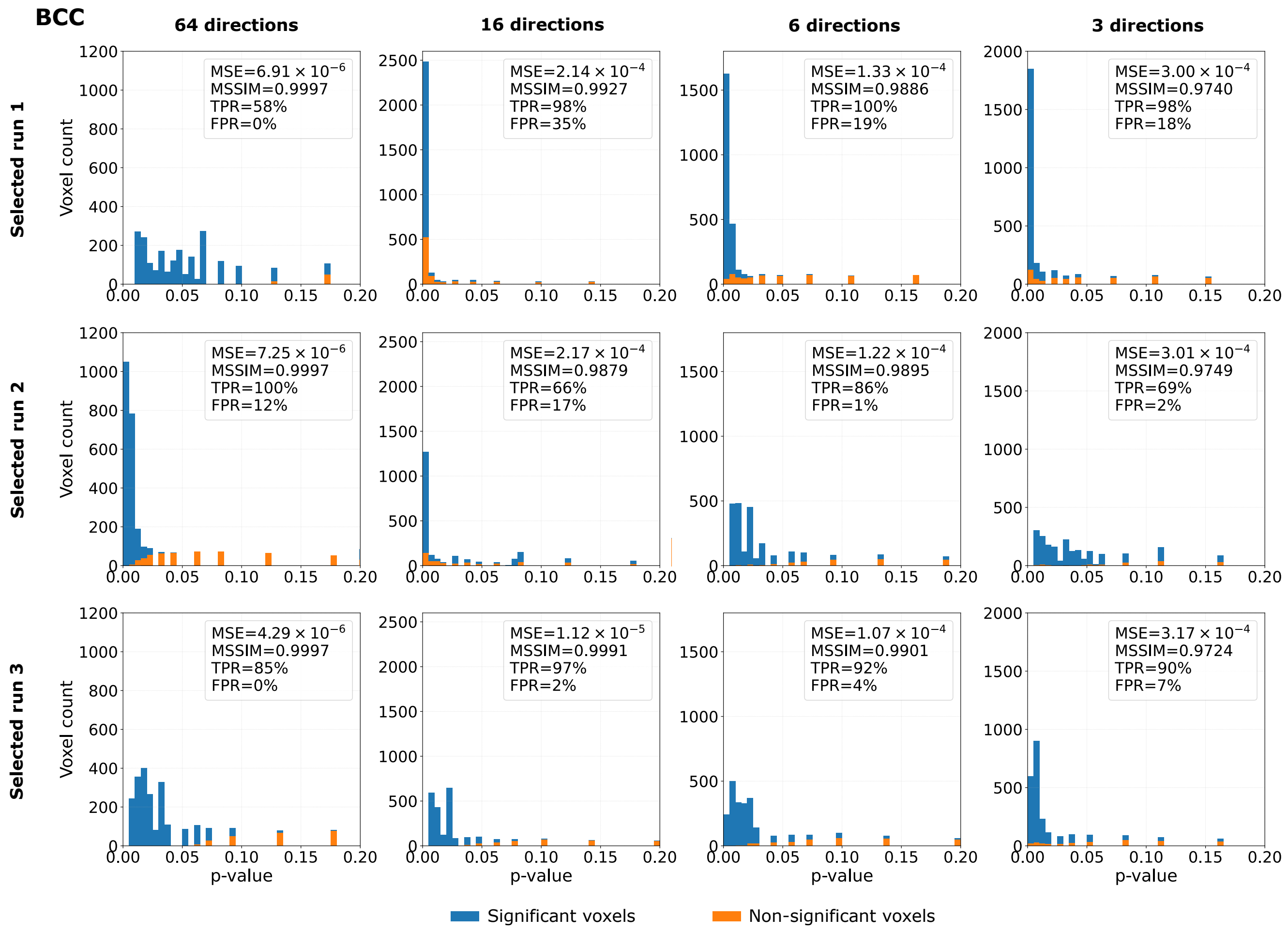


Figure S26: Histograms of voxel-wise $p$-values obtained from the tract-based spatial statistics (TBSS) analysis for the **RD parameter** estimated using a deep learning model following **Configuration 1** over the skeleton of the body (BCC) and splenium (SCC) of the corpus callosum and the left superior longitudinal fasciculus (SLF L). The figure presents three selected runs over 100 runs characterized with a comparable mean squared error (MSE) and mean structural similarity index measure (MSSIM), while variable true positive and false positive rates (TPR, FPR). For each run, four diffusion gradient distributions of input data are considered: 64, 16, 6, and 3. Bins in blue refer to the voxels, which were statistically significant in the TBSS analysis based on the standard NLLS-derived DTI (no deep learning applied), while orange bins show the voxels statistically non-significant in the TBSS analysis based on NLLS-derived DTI.

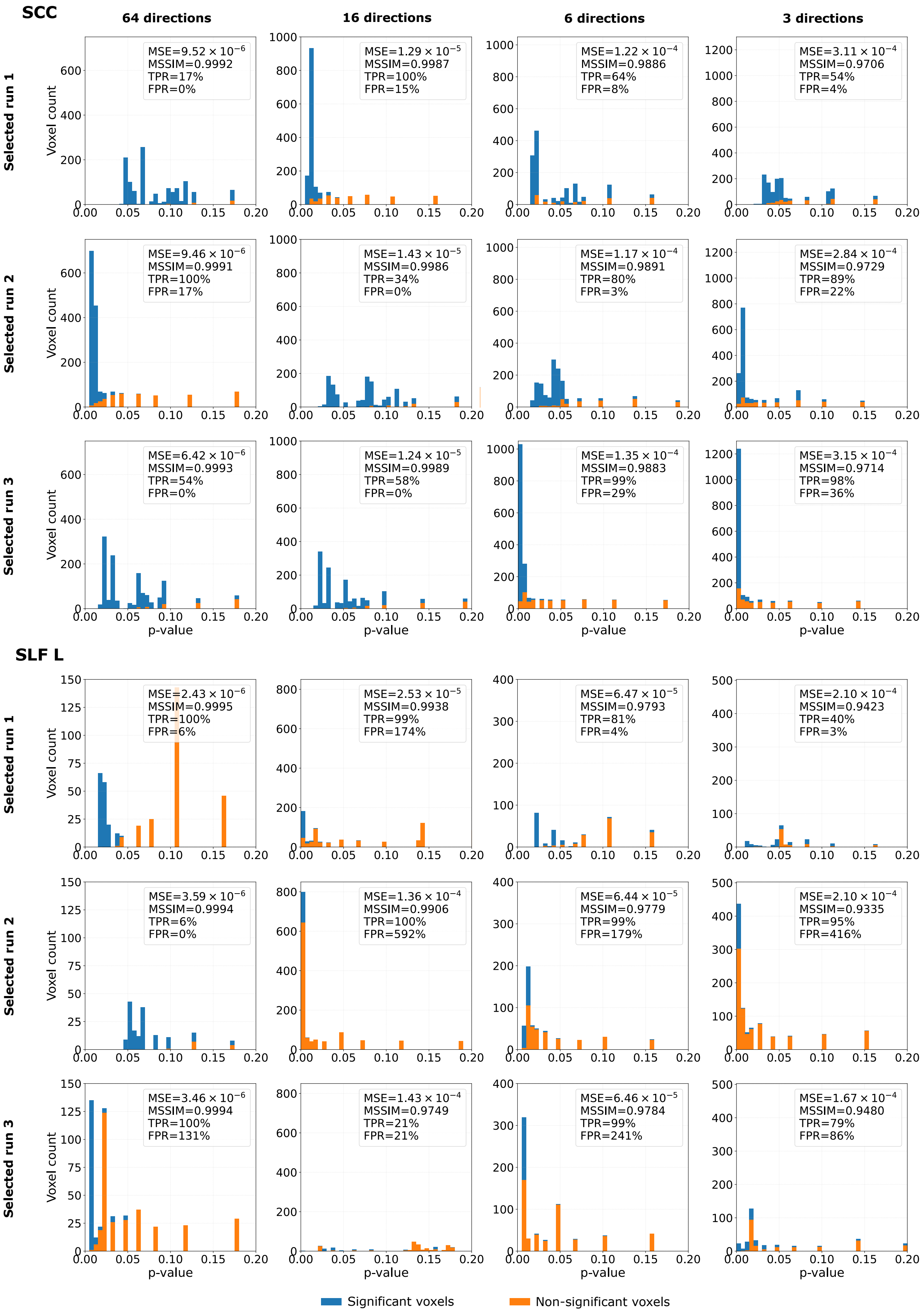


Figure S26 (continued)

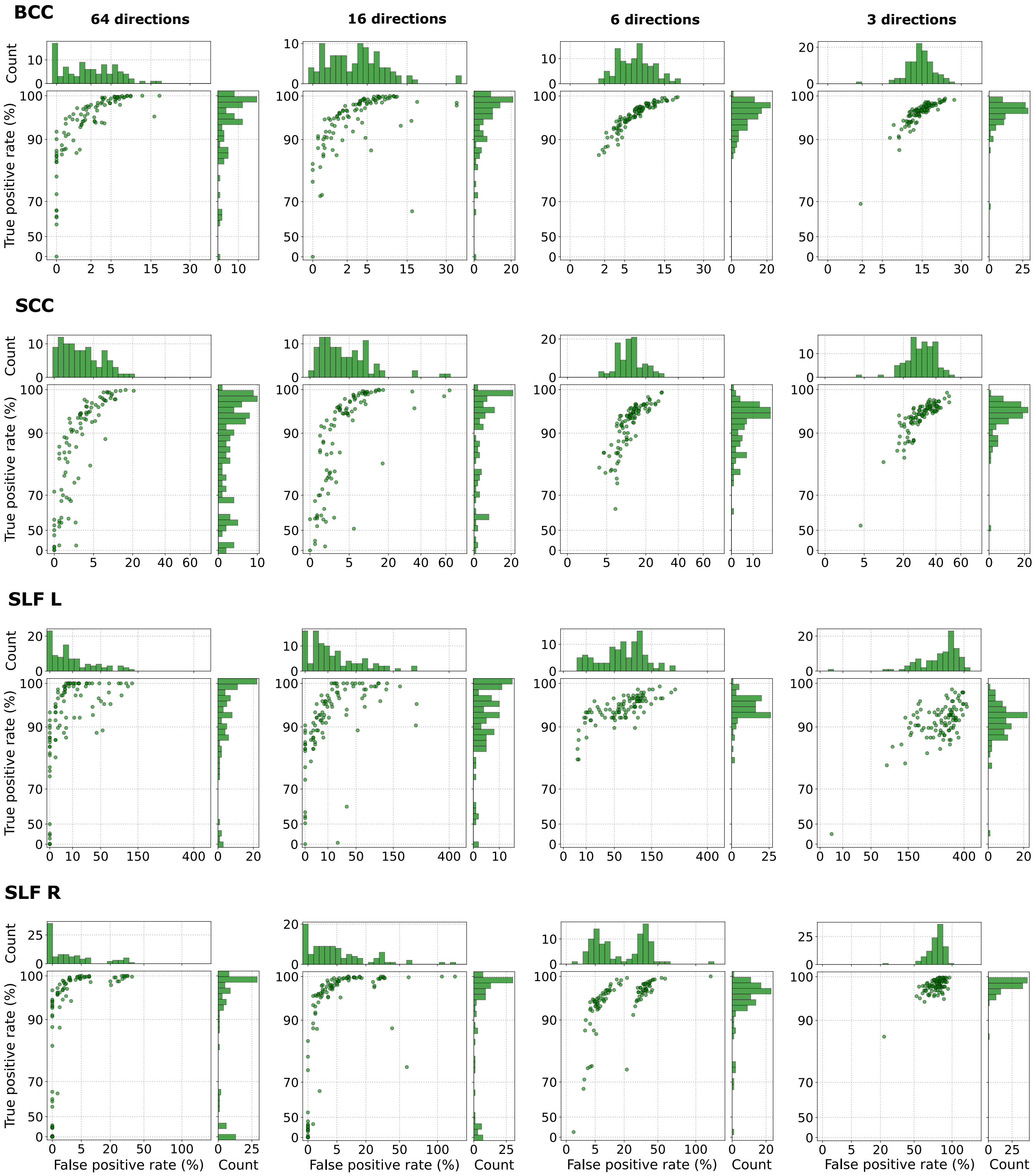


Figure S27: Relationship between the true positive rate and the false positive rate, obtained from the tract-based spatial statistics (TBSS) analysis for **RD parameter**, in the body (BCC) and splenium (SCC) of the corpus callosum and the left (SLF L) and right (SLF R) superior longitudinal fasciculus using **Configuration 1**, with four different numbers of input data gradient directions (64, 16, 6, and 3). Each point refers to a single run. In total, 100 runs were considered. The FPRs greater than 100% result from computing the values relative to the number of significant voxels identified in the reference analysis.

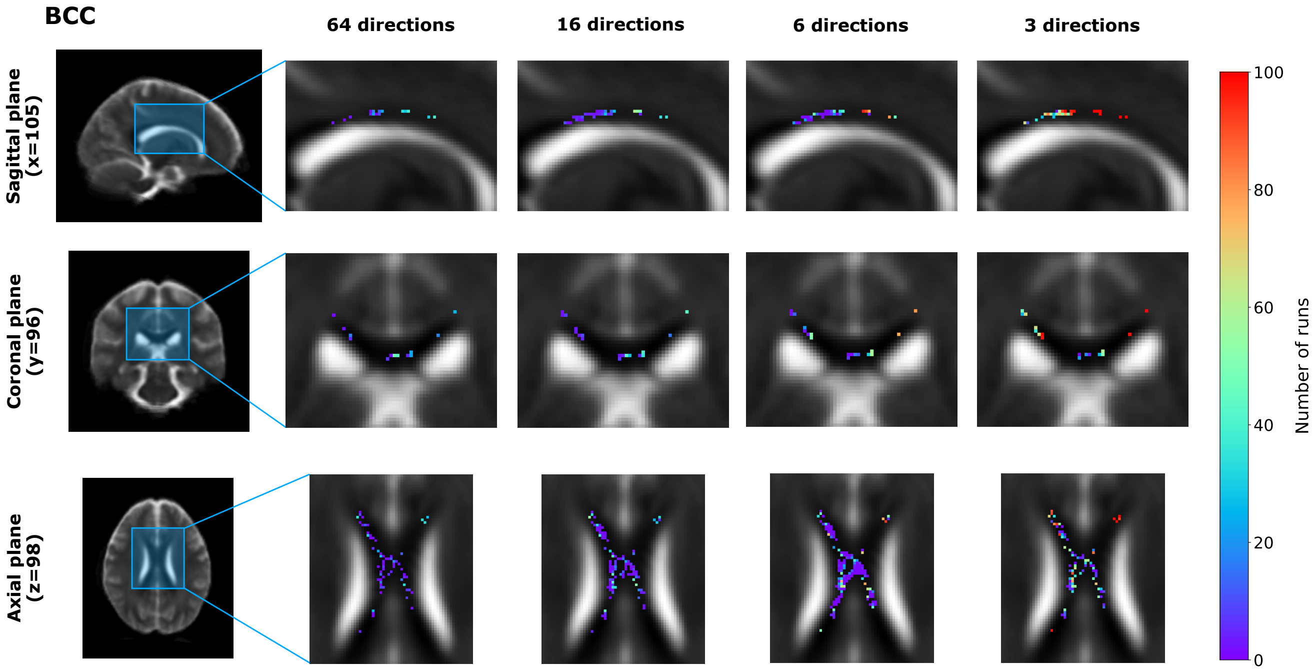


Figure S28: Visual representation of false positive voxels, obtained from the tract-based spatial statistics (TBSS) analysis for **RD parameter**, in the body of the corpus callosum (BCC) shown in the sagittal, coronal, and axial planes for four input gradient directions: 64, 16, 6, and 3 (**Configuration 1**). False positive voxels correspond to locations identified as exhibiting statistically significant group differences in the estimated RD maps but not in the reference analysis. The color indicates the number of runs in which a given voxel was classified as a false positive.

Table S7: Decision agreement between DL-based and reference ROI analyses across four different numbers of input data gradient directions (64, 16, 6, and 3) (**Configuration 1**). The reference $p$-value and significance decision are reported for each ROI, while decision agreement indicates the percentage of runs out of 100 that reproduced the statistical significance of the reference analysis ($p < 0.05$ or $p \geq 0.05$).

| **ROIs** | **Reference $p$-value** | **Reference decision** | **Decision agreement** | | | |
|---|---|---|---|---|---|---|
| | | | **64 directions** | **16 directions** | **6 directions** | **3 directions** |
| GCC | 0.0052 | Sig ($p < 0.05$) | 99% | 100% | 100% | 100% |
| BCC | 0.0348 | Sig ($p < 0.05$) | 69% | 74% | 95% | 99% |
| SCC | 0.0478 | Sig ($p < 0.05$) | 47% | 49% | 79% | 98% |
| ALIC R | 0.0206 | Sig ($p < 0.05$) | 89% | 89% | 93% | 99% |
| ALIC L | 0.0756 | Non-sig ($p \geq 0.05$) | 68% | 58% | 26% | 1% |
| PLIC R | 0.5684 | Non-sig ($p \geq 0.05$) | 98% | 95% | 98% | 99% |
| RIC R | 0.0448 | Sig ($p < 0.05$) | 60% | 53% | 72% | 98% |
| RIC L | 0.3974 | Non-sig ($p \geq 0.05$) | 99% | 98% | 99% | 18% |
| ACR R | 0.0020 | Sig ($p < 0.05$) | 100% | 100% | 100% | 100% |
| ACR L | 0.0008 | Sig ($p < 0.05$) | 100% | 100% | 100% | 100% |
| SCR R | 0.3520 | Non-sig ($p \geq 0.05$) | 99% | 95% | 97% | 96% |
| SCR L | 0.4404 | Non-sig ($p \geq 0.05$) | 99% | 96% | 97% | 83% |
| PCR R | 0.1350 | Non-sig ($p \geq 0.05$) | 98% | 96% | 96% | 51% |
| PCR L | 0.0610 | Non-sig ($p \geq 0.05$) | 63% | 46% | 18% | 0% |
| PTR R | 0.0068 | Sig ($p < 0.05$) | 99% | 100% | 100% | 99% |
| PTR L | 0.0096 | Sig ($p < 0.05$) | 99% | 99% | 100% | 99% |
| SS R | 0.0258 | Sig ($p < 0.05$) | 96% | 97% | 100% | 98% |
| SS L | 0.0788 | Non-sig ($p \geq 0.05$) | 90% | 69% | 43% | 15% |
| EC R | 0.4152 | Non-sig ($p \geq 0.05$) | 99% | 96% | 95% | 72% |
| EC L | 0.0100 | Sig ($p < 0.05$) | 100% | 98% | 100% | 99% |
| SLF R | 0.0540 | Non-sig ($p \geq 0.05$) | 43% | 30% | 29% | 1% |
| SLF L | 0.6162 | Non-sig ($p \geq 0.05$) | 99% | 98% | 99% | 100% |

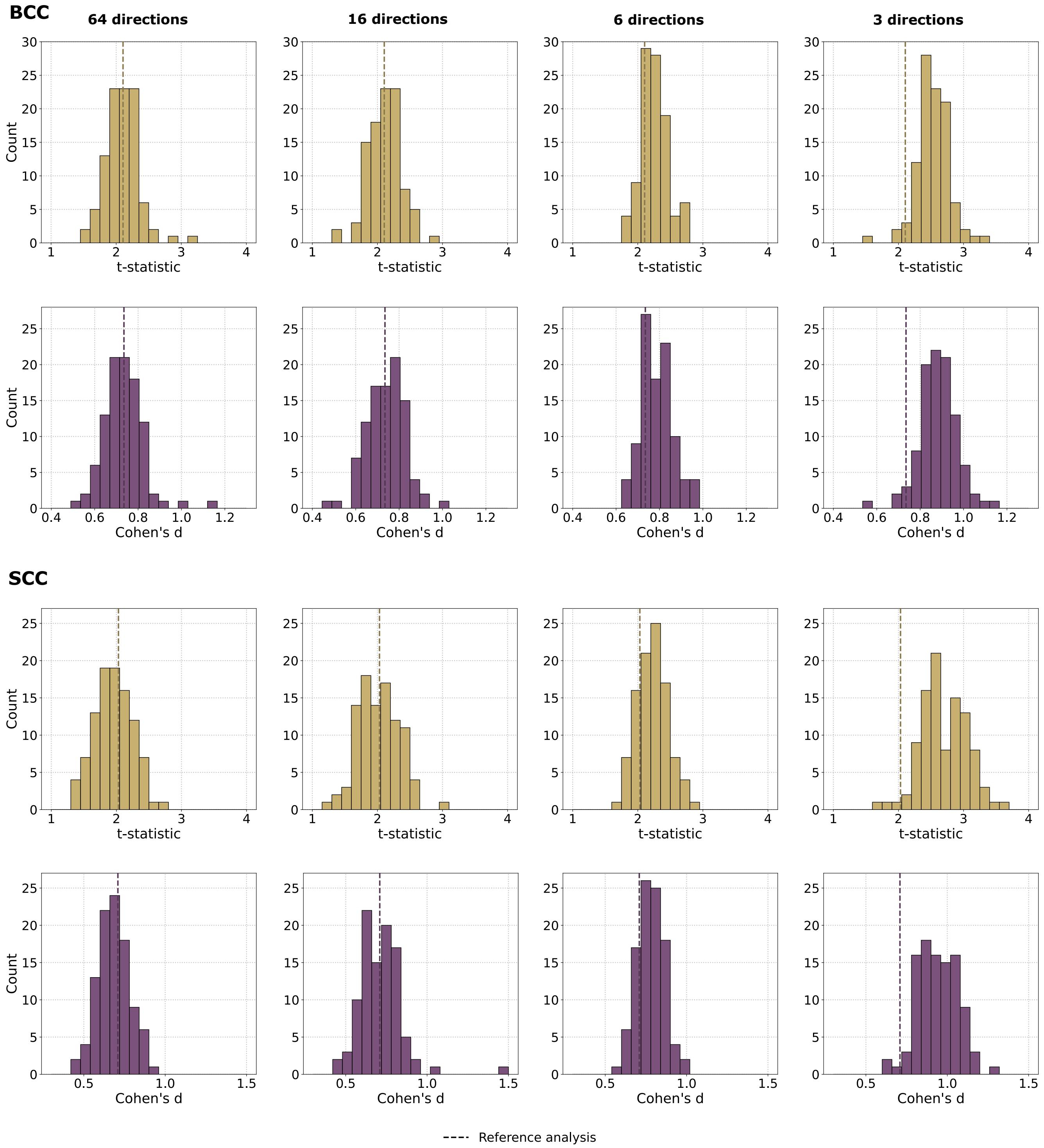


Figure S29: Distributions of $t$-statistics (top row) and Cohen's $d$ values (bottom row) in the body (BCC) and splenium (SCC) of the corpus callosum and the left superior longitudinal fasciculus (SLF L) for the **RD parameter** estimated using a deep learning model, with four different numbers of input data gradient directions (64, 16, 6, and 3) (**Configuration 1**). These statistics were calculated by comparing the mean RD values between patients with schizophrenia (SCH) and healthy controls (HC). Each histogram summarizes the results obtained from 100 runs. The dashed vertical line indicates the corresponding value from the reference analysis using the NLLS estimation.

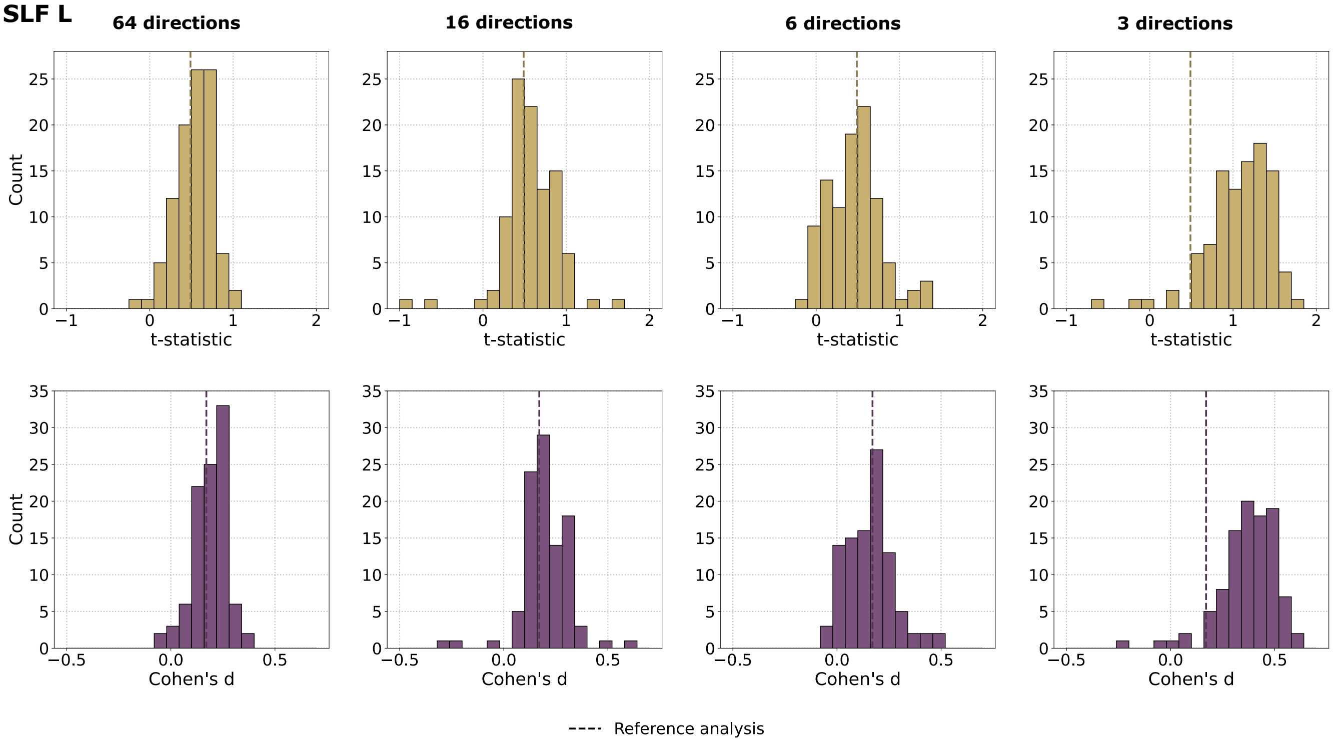


Figure S29 (continued)

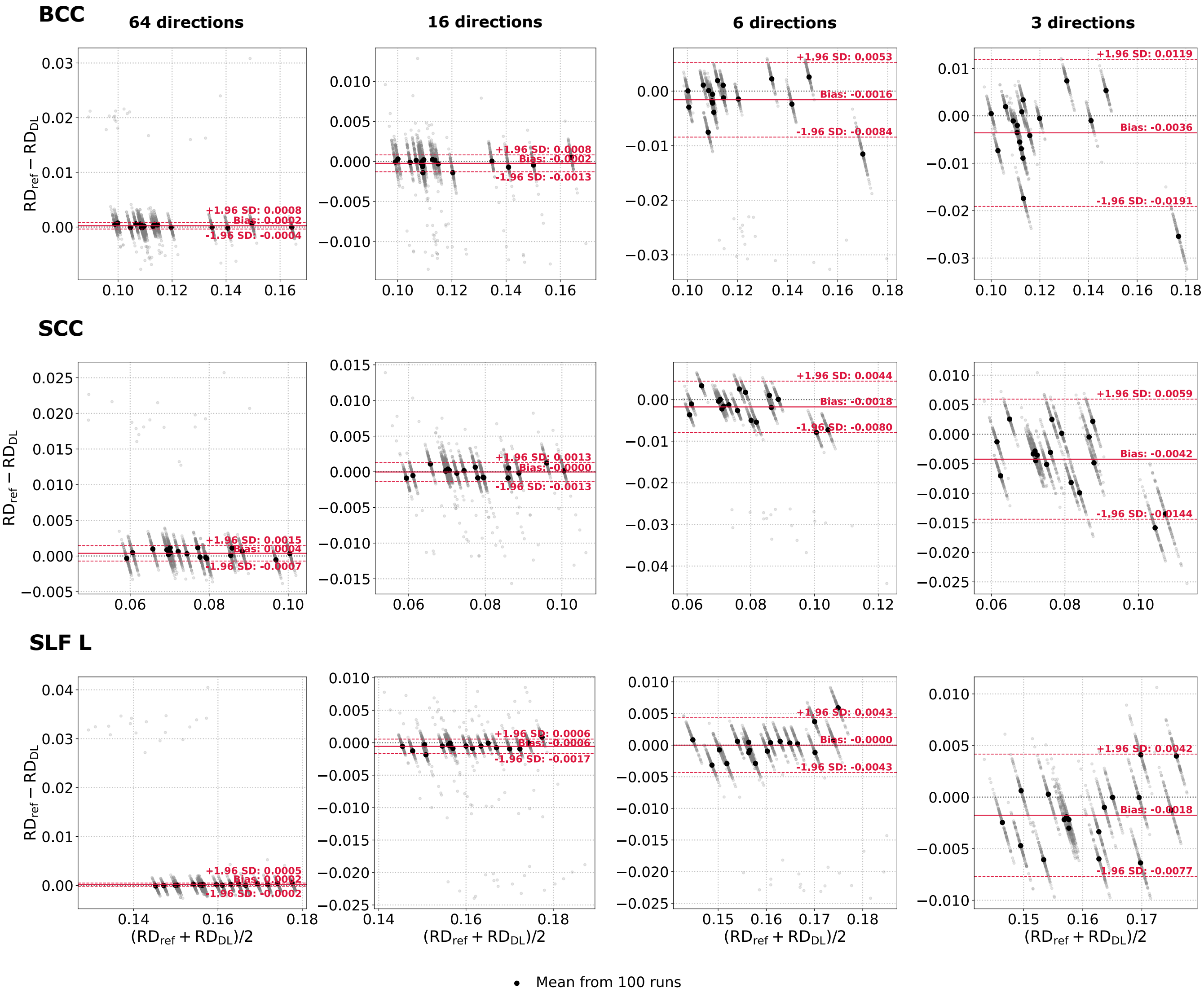


Figure S30: Bland-Altman plots of the **RD parameter** computed from the body (BCC) and splenium (SCC) of the corpus callosum and the left superior longitudinal fasciculus (SLF L) across all SCH subjects. The reference values ($RD_{ref}$) were obtained using the NLLS estimation from a resampled 64-direction scheme, whereas the predicted $RD_{DL}$ values were generated using the deep learning model (**Configuration 1**) considering four different numbers of input data gradient directions (64, 16, 6, and 3). Gray points represent individual model runs (mean values over the ROI). A single black point corresponds to the average across 100 runs for a single SCH patient. Solid red lines indicate the bias, while dashed red lines denote 95% limits of agreement (±1.96 standard deviations).

### References


Caruyer, E., Lenglet, C., Sapiro, G., Deriche, R., 2013. Design of multishell sampling schemes with uniform coverage in diffusion mri. Magnetic resonance in medicine 69, 1534–1540.

Descoteaux, M., Angelino, E., Fitzgibbons, S., Deriche, R., 2007. Regularized, fast, and robust analytical Q-ball imaging. Magn Reson Med 58, 497–510.

Fu, G., Zhang, W., Dai, J., Liu, J., Li, F., Wu, D., Xiao, Y., Shah, C., Sweeney, J.A., Wu, M., et al., 2019. Increased peripheral interleukin 10 relate to white matter integrity in schizophrenia. Frontiers in neuroscience 13, 52.

Gao, J., Yu, D., Yin, M., Li, J., Zhang, X., Tang, X., Zhang, X., 2025. Distinct white matter abnormalities and cognitive impairments in deficit schizophrenia: A cross-sectional diffusion tensor imaging study. Journal of Psychiatric Research 181, 381–390.

Holleran, L., Ahmed, M., Anderson-Schmidt, H., McFarland, J., Emsell, L., Leemans, A., Scanlon, C., Dockery, P., McCarthy, P., Barker, G.J., et al., 2014. Altered interhemispheric and temporal lobe white matter microstructural organization in severe chronic schizophrenia. Neuropsychopharmacology 39, 944–954.

Jones, D.K., Horsfield, M.A., Simmons, A., 1999. Optimal strategies for measuring diffusion in anisotropic systems by magnetic resonance imaging. Magnetic Resonance in Medicine 42, 515–525.

Lee, D.K., Lee, H., Park, K., Joh, E., Kim, C.E., Ryu, S., 2020. Common gray and white matter abnormalities in schizophrenia and bipolar disorder. PLoS One 15, e0232826.

Mamah, D., Patel, A., Chen, S., Wang, Y., Wang, Q., 2025. Diffusion basis spectrum imaging of white matter in schizophrenia and bipolar disorder. Brain Imaging and Behavior 19, 1002–1017.

Rigucci, S., Santi, G., Corigliano, V., Imola, A., Rossi-Espagnet, C., Mancinelli, I., De Pisa, E., Manfredi, G., Bozzao, A., Carducci, F., et al., 2016. White matter microstructure in ultra-high risk and first episode schizophrenia: a prospective study. Psychiatry Research: Neuroimaging 247, 42–48.

Scheel, M., Prokscha, T., Bayerl, M., Gallinat, J., Montag, C., 2013. Myelination deficits in schizophrenia: evidence from diffusion tensor imaging. Brain Structure and Function 218, 151–156.

Seal, M.L., Yücel, M., Fornito, A., Wood, S.J., Harrison, B.J., Walterfang, M., Pell, G.S., Pantelis, C., 2008. Abnormal white matter microstructure in schizophrenia: a voxelwise analysis of axial and radial diffusivity. Schizophrenia research 101, 106–110.

Zeng, B., Ardekani, B.A., Tang, Y., Zhang, T., Zhao, S., Cui, H., Fan, X., Zhuo, K., Li, C., Xu, Y., et al., 2016. Abnormal white matter microstructure in drug-naive first episode schizophrenia patients before and after eight weeks of antipsychotic treatment. Schizophrenia research 172, 1–8.
